# Genetics of fasting indices of glucose homeostasis using GWIS unravels tight relationships with inflammatory markers

**DOI:** 10.1101/496802

**Authors:** Iryna O. Fedko, Michel G. Nivard, Jouke-Jan Hottenga, Liudmila Zudina, Cross Consortia Pleiotropy (XC-Pleiotropy) Group, CHARGE Inflammation working group, Zhanna Balkhiyarova, Daniel I. Chasman, Santhi Ganesh, Jie Huang, Mike A. Nalls, Christopher J. O’Donnell, Guillaume Paré, Paul M. Ridker, Meta-Analyses of Glucose and Insulin-related traits Consortium (MAGIC) Investigators, Reedik Mägi, Marika Kaakinen, Inga Prokopenko, Dorret I. Boomsma

## Abstract

**Purpose:** Homeostasis Model Assessment of β-cell function and Insulin Resistance (HOMA-B/-IR) indices are informative about the pathophysiological processes underlying type 2 diabetes (T2D). Data on both fasting glucose and insulin levels are required to calculate HOMA-B/-IR, leading to underpowered Genome-Wide Association studies (GWAS) of these traits.

**Methods:** We overcame such power loss issues by implementing Genome-Wide Inferred Statistics (GWIS) approach and subsequent dense genome-wide imputation of HOMA-B/-IR summary statistics with SS-imp to 1000 Genomes project variant density, reaching an analytical sample size of 75,240 European individuals without diabetes. We dissected mechanistic heterogeneity of glycaemic trait/T2D loci effects on HOMA-B/-IR and their relationships with 36 inflammatory and cardiometabolic phenotypes.

**Results:** We identified one/three novel HOMA-B (*FOXA2*)/HOMA-IR (*LYPLAL1, PER4,* *PPP1R3B*) loci. We detected novel strong genetic correlations between HOMA-IR/-B and Plasminogen Activator Inhibitor 1 (PAI-1, *r*_*g*_=0.92/0.78, P=2.13×10^-4^/2.54×10^-3^). HOMA-IR/-B were also correlated with C-Reactive Protein (*r*_*g*_=0.33/0.28, P=4.67×10^-3^/3.65×10^-3^). HOMA-IR was additionally correlated with T2D (*r*_*g*_=0.56, P=2.31×10^-9^), glycated haemoglobin (*r*_*g*_=0.28, P=0.024) and adiponectin (*r*_*g*_=-0.30, P=0.012).

**Conclusion:** Using innovative GWIS approach for composite phenotypes we report novel evidence for genetic relationships between fasting indices of insulin resistance/beta-cell function and inflammatory markers, providing further support for the role of inflammation in T2D pathogenesis.

## Introduction

Type 2 diabetes (T2D) represents the fastest growing non-communicable disease epidemic world-wide^1^. Fasting hyperglycaemia is one of the criteria used for type 2 diabetes (T2D) diagnosis, and fasting insulin (FI) levels are indicative of insulin sensitivity in peripheral tissues, but neither measure provides mechanistic insights into insulin secretion and action^2^. Homeostasis Model Assessment of β-cell function (HOMA-B) and Insulin Resistance (HOMA-IR) can be derived from FG and FI concentrations and are two commonly used fasting state glycaemic indices elucidating pathophysiological processes in T2D^3-11^. HOMA-B reflects the function of β-cells in terms of their ability to secrete insulin, whereas HOMA-IR is a surrogate measure of insulin sensitivity.

Pathophysiology of T2D and metabolic syndrome, as well as epidemiological and genetic studies suggest that there is a shared aetiology between cardiometabolic phenotypes, variability of glycaemic traits in individuals without diabetes, and T2D^9,12-14^. Sedentary lifestyle and major risk factors of T2D induce low-grade inflammation, which consequently leads to overt diabetes in individuals with more pronounced and prolonged inflammatory response^15^ and insulin resistance. However, genetic relationships between these traits are not established. Moreover, metabolic syndrome is a pro-thrombotic state due to the inhibition of the fibrinolytic pathway, another proposed risk factor of T2D related to insulin resistance and inflammation^16,17^, but little is known about their shared genetic risk factors. The shared genetic aetiology within cluster of cardiometabolic traits was only recently reported through estimation of genetic correlations^18,19^.

Our understanding of biological processes shared between β-cell function and insulin resistance, reflected by HOMA-B/-IR, and a range of epidemiologically related traits and diseases, such as T2D, could be informed by dissecting the patterns of their genetic relationships. Genome-wide association studies (GWAS) have identified over 70 loci associated with fasting glucose (FG) and/or FI levels in non-diabetic European descent individuals^20^. However, to date, only ten HOMA-B loci, including *ADCY5, ARAP1(STARD10), DGKB, FADS1, GLIS3, GCK, G6PC2, MTNR1B, TCF7L2,* and *SLC30A8*, and three HOMA-IR loci (*GCKR, IGF1/SC4MOL, TCERG1L)* have been described in GWA studies in European/African American descent populations^21-25^. This is mostly attributable to underpowered studies, since HOMA-B and HOMA-IR calculations require assessment of both FG and FI in the same individual. If either of these two primary fasting measures is missing for an individual, their HOMA indices cannot be derived and such individuals do not contribute to GWAS of HOMA indices. Similarly, missing FG or FI measurement in an entire cohort precludes involvement of such study in GWAS meta-analyses of HOMA indices (**Figure 1**). As a consequence, HOMA-B/-IR GWAS meta-analyses usually feature dramatically smaller sample sizes compared to FG/FI GWAS. Previously published HOMA-B/-IR discovery analyses by the Meta-Analyses of Glucose and Insulin-related traits Consortium (MAGIC) suffered from about 20% sample size and power losses for analytically inferred indices^21^, which has held up the progress of understanding the molecular mechanisms behind insulin secretion and sensitivity composite measures through locus discovery in recent years^26,27;28,29^. The Genome Wide Inferred Statistics (GWIS) method provides an approximation of the GWA summary statistics for any derived variable that is a function of primary phenotypes, when such statistics, means and covariances of the constituent primary phenotypes are available or can be approximated with reasonable precision^30^. Unlike GWAS, the GWIS can accommodate information from individuals or cohorts, where any of the primary phenotypes is assessed. Moreover, the GWIS accommodates any degree of overlap between individuals from studies contributing data for primary traits. In this study, we applied GWIS methodology^30^ and derived the summary statistics for HOMA indices based on recent FG and FI GWAS meta-analysis summary statistics from the MAGIC (*Lagou et al., manuscript in preparation*).

**Figure 1.**
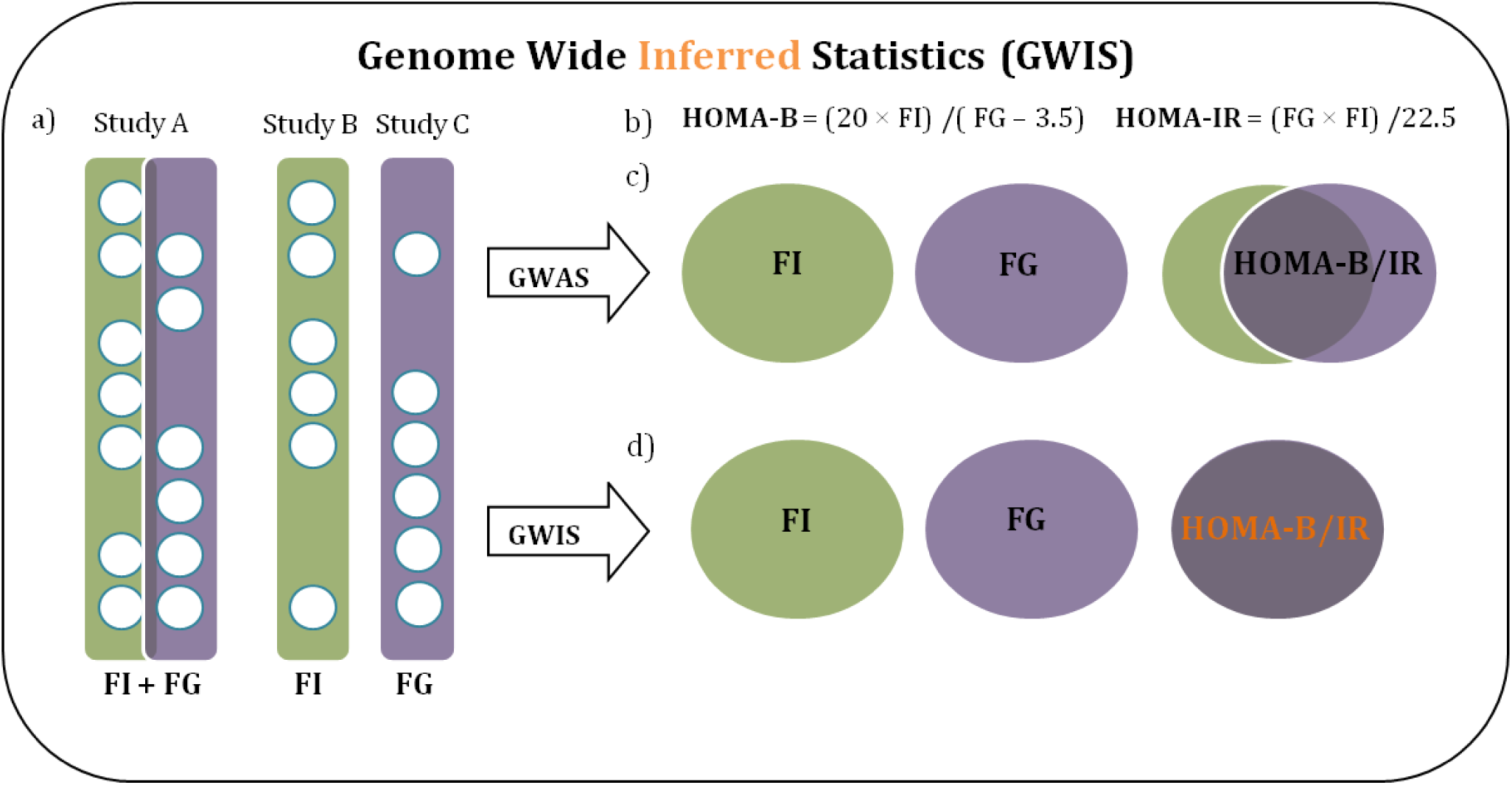
Comparison between GWAS meta-analysis of primary phenotypes and GWIS of derived phenotypes. Input phenotypes are fasting glucose (FG) and fasting insulin (FI), output phenotypes are Homeostasis Model Assessment of β-cell function (HOMA-B) and insulin resistance (HOMA-IR). a) Examples of how FG and FI measures could partially overlap within one cohort (e.g. Study A), whereas some cohorts could only have measures of FG (Study C) or FI (Study B). Study A could participate in future GWAS by computing HOMA-B/IR for overlapping individuals, although the sample size will be decreased in comparison to original FG and FI phenotypes. Study B and C cannot contribute to HOMA-B/IR GWAS, thus reducing the sample size of GWAS meta-analysis of HOMA-B/IR to a larger extent. b) Formulae to compute HOMA-B/IR from FG and FI measures. c) Conventional GWAS approach, where HOMA-B/IR would be computed based on individual cohort studies, undergoing the reduction of the sample size, if FG and FI are not measured in the same individual. GWAS summary statistics would be then computed on reduced sample size. d) GWIS approach, where FG and FI meta-analyses summary statistics are computed separately with maximum sample size available. HOMA-B/-IR summary statistics are then calculated from FG and FI meta-analyses results. The sample size of derived phenotypes are computed as geometric mean between FG and FI sample sizes.

The *aims* of this study were three-fold: (i) to infer analytically HOMA-B/-IR GWIS based on the FG/FI GWAS meta-analysis summary statistics from the MAGIC consortium and evaluate the sensitivity of GWIS methodology; (ii) to use the obtained summary statistics to define the effects of T2D, FG/FI and HbA1_C_ loci on insulin secretion and action through their effects on HOMA-B/-IR; and (iii) to explore the genetic relationships between HOMA-B/-IR, cardiometabolic, and inflammatory traits.

## Methods

### Phenotypes

Homeostasis Model Assessment of β-cell function (HOMA-B) and Insulin resistance (HOMA-IR) are calculated from the Fasting Glucose and Fasting Insulin measures by formulae:

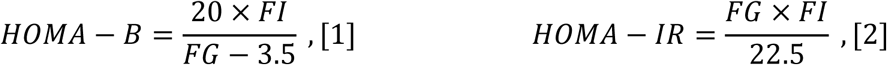

where FG is measured in mmol/l and FI is in mU/l units^31^.

### GWIS for HOMA-B/-IR

To approximate the HOMA-B/-IR GWAS summary statistics, we applied a recently developed GWIS approach^30^. We obtained the summary statistics from the latest GWAS meta-analysis of FG and FI performed by the MAGIC in up to 88,320/64,090 individuals and 40/33 studies respectively (**Supplementary Materials and Methods**) (*Lagou et al., manuscript in preparation*). Studies genotyped on Metabochip were not included, so that inferred summary statistics were appropriate to analyse with LD score regression (LDSC)^32^. In the MAGIC meta-analysis FI was measured in pmol/l and natural log transformed, FG was measured in mmol/l with a cut-off at 7mmol/l. The standard HOMA formulae require untransformed FG/FI measures and use mU/l units for FI. We adapted the HOMA formulae to compute a GWAS summary statistics for ln(HOMA-IR) and ln(HOMA-B) given summary statistics for FG and ln(FI), measured in the units described above:

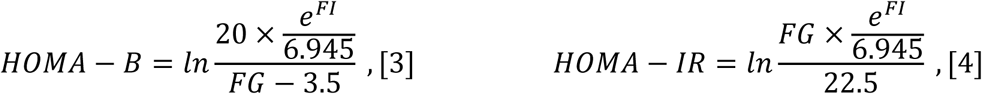

where division by 6.945 was introduced to convert FI from pmol/l to mU/l units. The GWIS method requires, in addition to genome wide summary statistics for FG and FI, the population mean of FG and FI, the phenotypic correlation between FG and FI and sample overlap across the studies, included in both meta-analyses studies, to correct for dependence between the FG and FI GWAS (details in **Supplemental Materials and Methods** and example scripts in **Data Access**). Overall, we calculated the effect estimates (β’s) for HOMA-B and HOMA-IR for each SNP_*i*,_ *i*=1,…,M, where M is the number of SNPs. According to the GWIS method β_i_ is computed as a sum of HOMA-B or HOMA-IR functions respectively (versions of the formulae given above) derived using population means and published summary statistics for FG and FI, which are evaluated for 1 and 2 copies of effect alleles and weighted by the estimated frequencies of effect alleles in FI and FG GWAS summary statistics (**Supplementary Materials and Methods**). For the calculation of the standard errors (SEs) of the GWIS-inferred effect estimates of HOMA-B and HOMA-IR we used the Delta method accounting for the sample overlap (**Supplementary Materials and Methods**). We compared the magnitude and direction of the effects in genome-wide significant loci in published and inferred analysis and confirmed power gain, especially for HOMA-IR, when compared to previous analysis. (**Tables S1 and S2**, **Figures S1-S4**).

We run the LDSC^33^ between the GWIS-inferred and published summary statistics for HOMA-B and HOMA-IR, to obtain the LDSC intercept for the GWIS-inferred HOMA-B/-IR and to estimate the genetic correlation between inferred and published data. We used LDSC intercept to correct the inferred summary statistics for any inflation (**Supplementary Materials and Methods**).

### Genome-wide imputation using summary statistics

We implemented a novel methodology developed within SS-imp software tool that enables genome-wide imputation from denser reference panels^34^ and imputed the analytically inferred HOMA-B/-IR GWIS to 1000 Genomes project variant density and compared the findings with the lower variant density directly inferred GWIS.

### Functional and regulatory elements enrichment analysis

We applied the GARFIELD^35^ tool v2 on the meta-analysis results to assess enrichment of the HOMA-B/HOMA-IR associated variants within functional and regulatory features. GARFIELD enables checking for genic annotations, chromatin states, DNaseI hypersensitive sites, transcription factor (TF) binding sites, FAIRE-seq elements and histone modifications, among others, in a number of publicly available cell lines. We considered significant enrichment to be present if the GWIS signal and the functional annotation signal significantly co-localized, i.e. *P*_GWIS_<5×10^-8^ and *P*_enrichment_<1.2×10^-5^ after correction for 2,040 annotations.

### Genetic correlation between HOMA indices and other phenotypes

To evaluate the shared genetic aetiology, we applied the LDSC approach^33^ to HOMA-B/-IR and publicly available meta-analysis summary statistics for T2D, 13 glycaemic and 17 cardiometabolic phenotypes (sample size, ethnicity, reference and source of the data presented in **Table S3**). We defined an extended group of cardiometabolic phenotypes and related traits by including chronic kidney disease (CKD) and its markers (Estimated glomerular filtration rate (eGFR) based on creatinine and cystatin C)^36^; systolic and diastolic blood pressure (SBP/DBP) and hypertension (HTN)^37^ to explore the genetic relationships between cardiometabolic traits and HOMA-B/-IR. We expanded our analysis to five inflammatory markers, including adiponectin^38^, plasminogen activator inhibitor 1 (PAI-1)^39^, C-reactive protein (CRP)^40^, intercellular adhesion molecule 1 (ICAM-1)^41^, white blood cell counts (WBC)^42^). The five inflammation phenotypes were obtained with permission of Cross Consortia Pleiotropy (XC-Pleiotropy) Group (**Table S3**).

## Results

### GWAS for HOMA-B/-IR

We inferred analytically the HOMA-B/-IR GWAS summary statistics using the summary statistics of HapMap reference panel-imputed FG/FI GWAS meta-analyses of 40/33 studies by the MAGIC (*Lagou et al., in preparation*). We imputed them to 1000 Genomes phase 3 (2013-05-02) reference panel thus reaching ∼11.2M autosomal variants for up to 75,240 non-diabetic individuals of European descent. We detected a novel HOMA-B locus at *FOXA2* gene (rs5029909, β(SE)=-0.044(0.0079), P=2.70×10^-08^), which overlapped with an established FG-association (rs6113722, R^2^=0.69 in Europeans, 1000G)^13^. We also confirmed associations at 10 established HOMA-B/FG loci (**Figure S5a, Table 1**)^13,21,43^. The HOMA-IR GWIS provided three novel loci at *LYPLAL1/SLC30A10, PER4, PPP1R3B* and confirmed established loci at *GCKR* and *IGF1*^21^ (**Figure S5b, Table 1**). Imputation using new tool SS-imp^34^ for summary statistics to 1000G density highlighted small difference from the lead variants at HOMA loci imputed to HapMap2 reference panel (**Table S4**). New lead 1000G-imputed variants at *TCF7L2, ARAP1[STARD10]* for HOMA-B, and at *GCKR* (rs11336847) for HOMA-IR, were in strong (r^2^>0.8) LD, and at *G6PC2* (rs580670 for HOMA-B in moderate (r^2^>0.6) LD with HapMap2 lead variants (**Table1**).

**Table 1.**
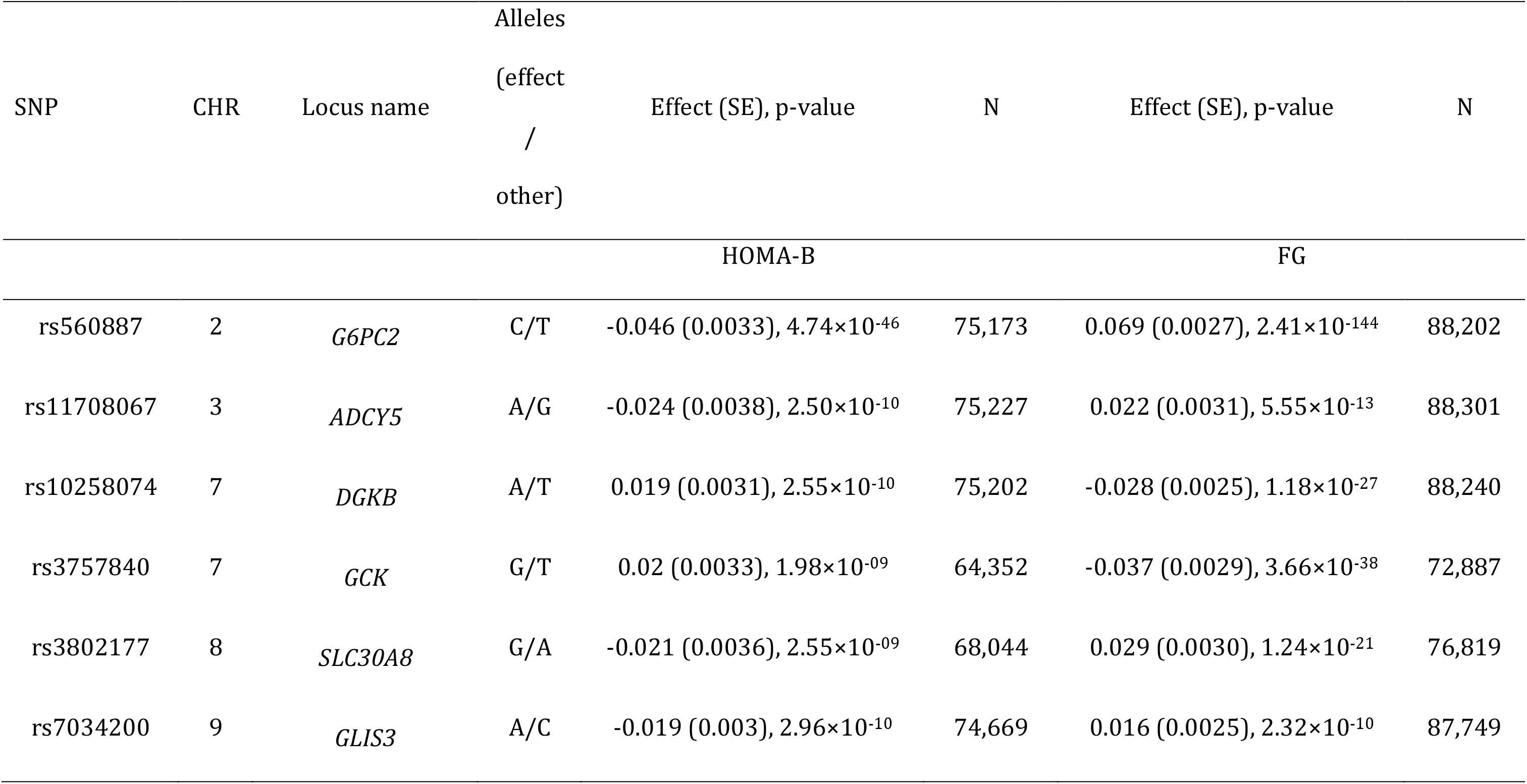
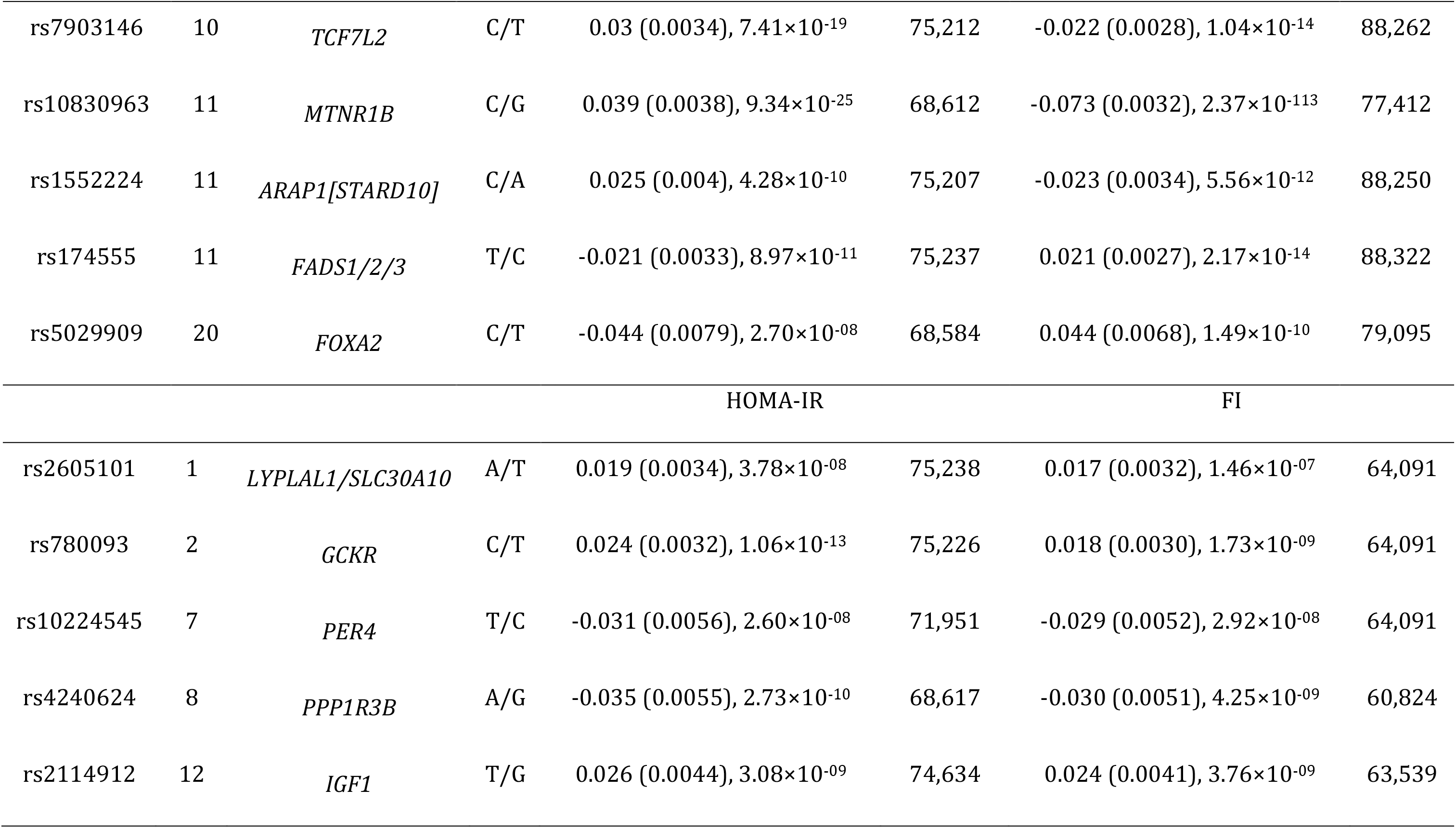
Comparison of HOMA-B/-IR and FG/FI effects across genome-wide significant loci

GWIS requires FG/FI population mean estimates to infer summary statistics for HOMA-B/-IR, which vary across studies. We performed a set of sensitivity analyses, where we allowed mean estimates to fluctuate between reasonable values. These analyses indicated that within 1 standard deviation (*SD)* from the two means (*M(SD)* = 5.25(0.34) mmol/l for FG and *M(SD)*=60.40(19.45) pmol/l for FI) 8 out of 11 HOMA-B loci were genome-wide significant in 100% of cases, except for *GCK* (72%), *SLC30A8* (86%) and *FOXA2* (55%) with slight p-value drops to 4.7×10^-7^, 1.2×10^-7^ and 2.7×10^-7^ respectively; and all 5 HOMA-IR loci were genome-wide significant irrespective of FG/FI means variation (**Figure S6**).

### Effects of established T2D, FG/FI and HbA1_C_ loci on HOMA-B/-IR

We investigated the effects of established 102 T2D, 50 FG/FI and 57 HbA1_C_^44^ loci on indices of β-cell function and insulin resistance (**Figure 2, Tables S5-S7**)^20,45-48^. As previous studies were performed using a number of reference panel imputations, the number of established T2D/FG/FI/HbA1_C_ SNPs that map to HOMA-B/-IR GWIS results is smaller than the number of currently reported SNPs in the literature (R^2^>=0.9 for the proxy SNPs). The effects of T2D loci on HOMA indices followed a grouping previously defined by us^29^. The largest group of loci associated at least nominally with HOMA-B and primarily leading to reduced insulin secretion included 20 T2D loci. Some of these loci have at least a nominal effect on improved insulin sensitivity which likely reflects compensatory mechanisms observed in individuals without known T2D used for the HOMA analyses, as previously discussed^29^. The effects of T2D loci on HOMA index of insulin resistance (**Figure 2b**) clearly demonstrate that, in addition to *IRS1, GCKR, KLF14* and *PPARG*, a number of more recently defined T2D loci, including *PEPD, ANKRD55, BCL2, ARL15* and *GRB14* are related to this pathophysiological process. *FTO, MC4R* and *NRXN3* are established body mass index (BMI) loci^49^ and therefore their observed effects on fasting insulin resistance are likely to be driven through their primary effects on adiposity.

**Figure 2.**
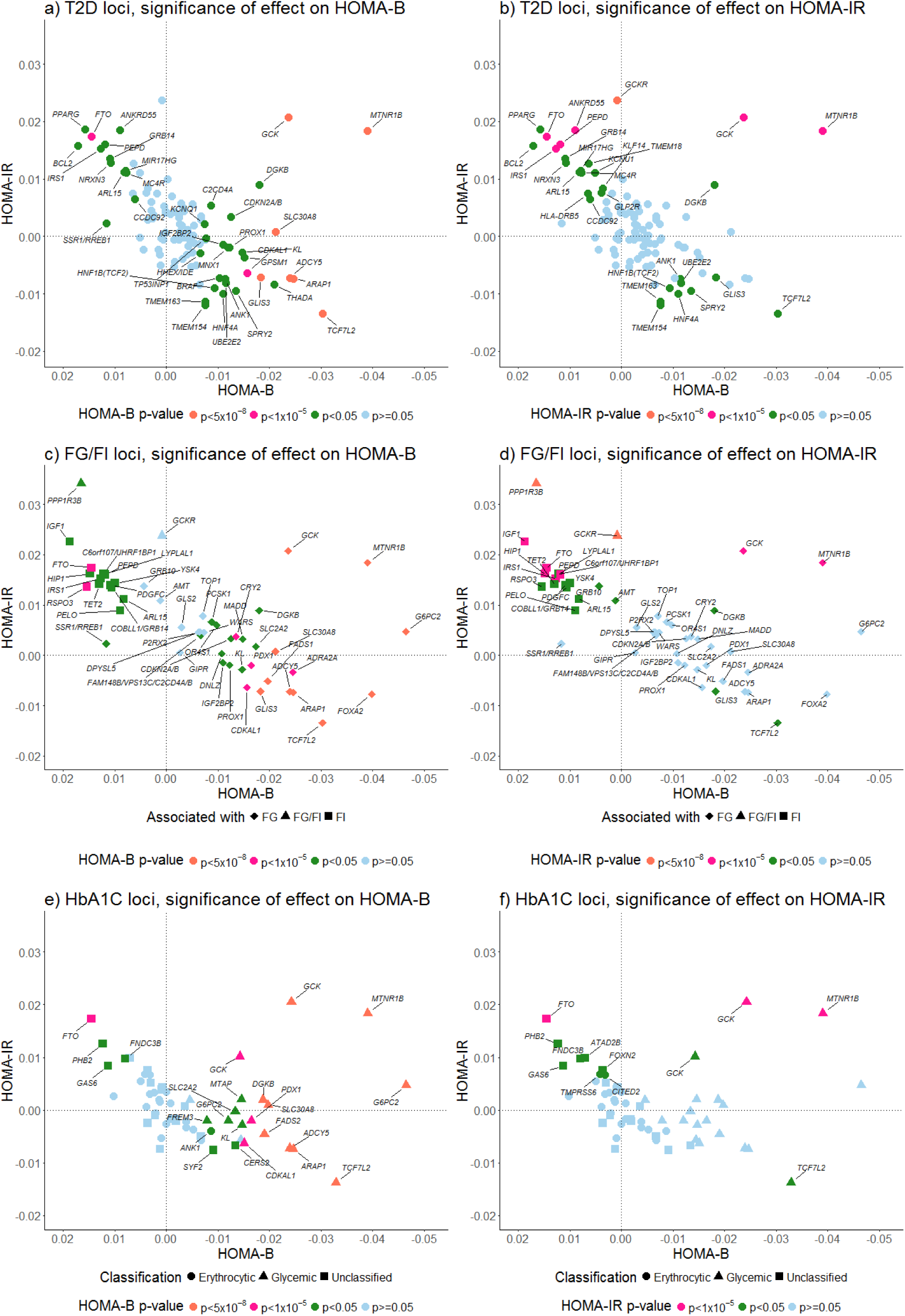
Effects of established T2D (a,b), FG/FI (c,d) and HbA1_C_ (e,f) loci on HOMA-B/-IR. Legend: X-axis and Y-axes represent the direction of loci effects on β-cell function and insulin resistance as reflected by HOMA-B/-IR, respectively. Note, *DUSP9* T2D locus is not included on the Figure 2 (a,b) as it resides on X chromosome. Effects of HOMA-B/-IR SNPs correspond to the risk alleles of either T2D, FG/FI or HbA1_C_ SNPs. The largest group of 20 T2D loci (*DGKB, CDKN2A/B, C2CD4A, TP53INP1, HHEX/IDE, UBE2E2, ANK1, SPRY2, TMEM163, TMEM154, HNF4A, PROX1, THADA, BRAF, MNX1, HNF1B(TCF2), GPSM1, IGF2BP2, KCNQ1, KL*) is associated at least nominally with HOMA-B and is primarily leading to reduced insulin secretion. The strongest effects on HOMA-B were observed for the loci characterised by the reduced β-cell function (*SLC30A8, ADCY5, TCF7L2, GLIS3* and *CDKAL1),* reduced proinsulin production and insulin secretion at *ARAP1(STARD10)*, and those related to fasting hyperglycaemia (*MTNR1B* and *GCK*).

Most of the established FG loci were at least nominally associated with the reduced β-cell function, as reflected by negative effect estimates for HOMA-B (**Figure 2c**), while FI loci were associated with decreased insulin sensitivity, i.e. provided positive effect estimates for HOMA-IR values (**Figure 2d**). Ten FG loci were genome-wide significant for HOMA-B (**Figure 2c**). FG level-increasing alleles at eight overlapping T2D loci (**Figure 2a**) represented by hyperglycaemia (*MTNR1B, GCK),* β-cell *(SLC30A8, PROX1, ADCY5, GLIS3, TCF7L2),* and proinsulin (*ARAP1[STARD10])* loci, as well as the FG only loci that do not have established associations with T2D (*G6PC2, FOXA2)* were associated with reduced β-cell function through their primary effect on the set point of glucose. All FI loci were at least nominally associated with increased insulin resistance detected by HOMA-IR (**Figure 2d**). At FI loci, we detected at least nominal effects on HOMA-B, which might be related to left truncation for FG values for these analyses which introduced seeming improvement in β-cell function^29^. The T2D effect allele in *GCKR* locus reflects solely the decreased insulin action, i.e. normal functioning of β-cells in the presence of increased insulin resistance (**Figure 2b**)^50,51^.

Nine of eleven genome-wide significant HOMA-B loci and four loci of suggestive significance (*GCK, PDX1, CDKAL1, FTO*) have established effects on HbA1_C_ levels^44^ (**Figure 2e**). The HbA1_C_ loci have been classified into erythrocytic or glycaemic, but a number of them remain unclassified. We suggest that five unclassified HbA1c loci (*PHB2, FNDC3B, GAS6, SYF2, CERS2*) are glycaemic as they exert nominal effects on HOMA-B. Similarly, the abovementioned *PHB2, FNDC3B, GAS6* as well as *ATA2B* and *FOXN2* are nominally associated with HOMA-IR and therefore are likely glycaemic.

### Enrichment in functional elements for HOMA-B/-IR associations

The enrichment analysis using the GARFIELD software did not yield statistically significant co-localization signals neither for HOMA-B nor HOMA-IR. We observed suggestive evidence (*P*<0.05) for an enrichment of the HOMA-B associated variants within several pathophysiologically relevant tissues, including blood, liver and brain. Both HOMA-B and HOMA-IR associated variants showed also suggestive evidence for enrichment in a number of annotations within K562 (human immortalised myelogenous leukemia cell line) and HepG2 (human immortalised liver carcinoma cell line) cells, including transcription start cites, weak enhancers among chromatin states, histone modifications, transcription factor binding cites and FAIRE-seq elements (**Figure S7**).

### Genetic relationships between HOMA indices and other phenotypes

LDSC^33^ was applied to estimate the genome-wide genetic correlation between HOMA-B/-IR and 36 other phenotypes falling into five broad domains (**Table S8**), including, T2D, 13 glycaemic traits, four blood lipids, four obesity traits, nine phenotypes indicative of T2D complications and five inflammation markers (**Figure 3**). HOMA-B showed positive and strong (r_g_=0.68[SE=0.07], P=4.33×10^-21^) genetic correlation with HOMA-IR in our data. We identified the strongest novel genetic correlation (FDR–adjusted to correct for multiple testing at P<0.05) in the domain of inflammatory markers, specifically between HOMA-IR and PAI-1 (r_g_=0.92, P=2.13×10^-4^), usually observed at higher levels in people with obesity and metabolic syndrome^39^. Similarly, HOMA-B directly correlated with PAI-1 (r_g_=0.78, P=2.54×10^-3^). CRP positively correlated with both HOMA indices (r_gHOMA-B/-IR_=0.28/0.33, P_HOMA-B/-IR_=3.65×10^-3^/4.67×10^-3^), while adiponectin showed significant inverse correlation with HOMA-IR only (r_g_=-0.30, P=0.012).

**Figure 3.**
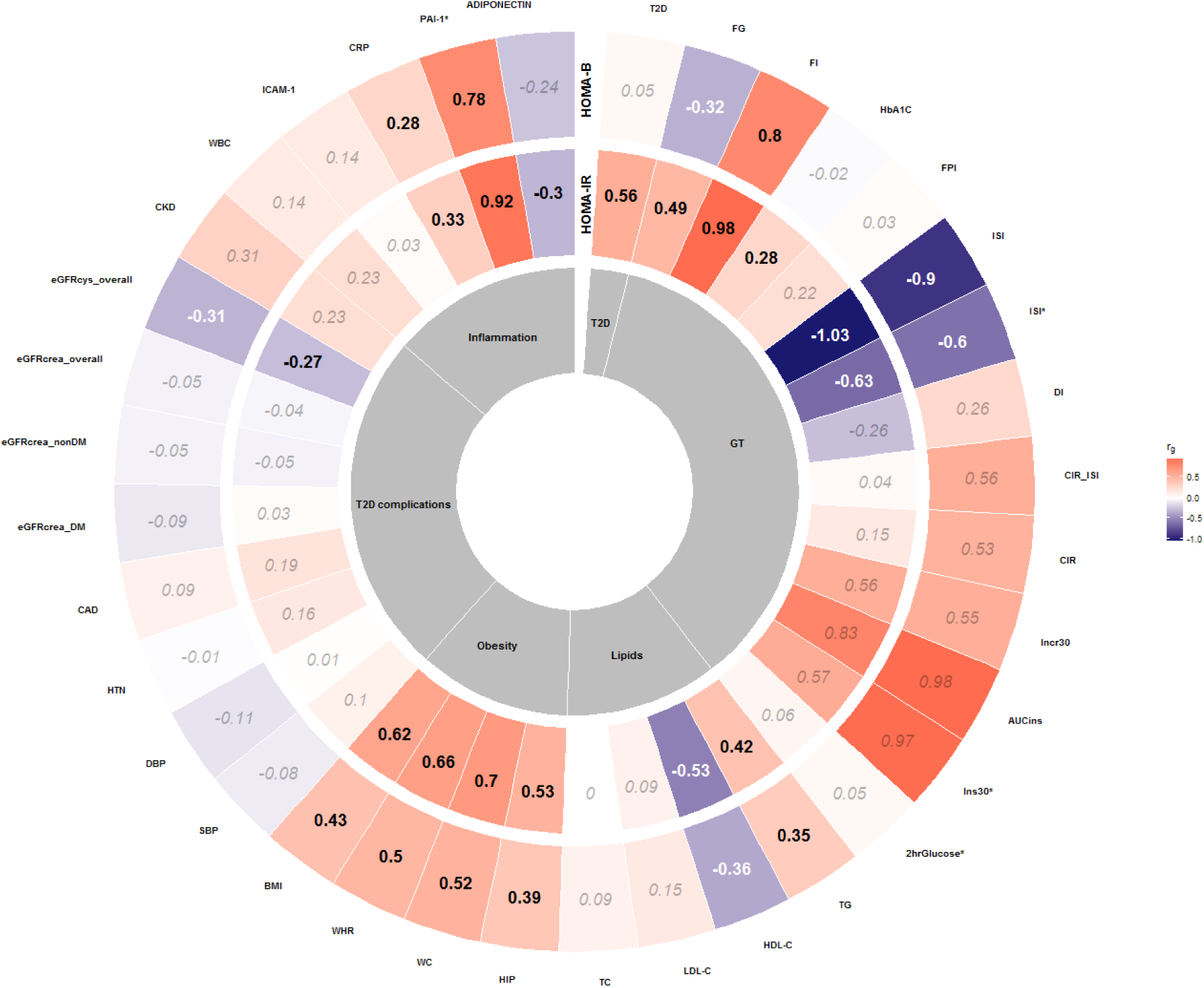
Genetic correlations (r_g_) between HOMA-B/-IR and 13 glycaemic, 17 cardiometabolic traits, five inflammation markers, and T2D. Genetic correlations reaching statistical significance (FDR–adjusted P < 0.05) are indicated in **bold black** in contrast to non-significant correlations reported in *italic grey*. An asterisk indicates BMI adjustment for specific phenotype. Phenotype abbreviations: type 2 diabetes (T2D), fasting glucose (FG), fasting insulin (FI), glycated haemoglobin (HbA1_C_), fasting pro-insulin (FPI), insulin sensitivity index (ISI), disposition index (DI), corrected insulin response (CIR), corrected insulin response adjusted for Insulin Sensitivity Index (CIR_ISI), incremental insulin at 30 min (Incr30), area under the curve of insulin levels during OGTT (AUCIns), insulin response at 30 min (Ins30), 2 hour Glucose (2hrGlucose), triglycerides (TG), high density lipoprotein cholesterol (HDL-C), low density lipoprotein cholesterol (LDL-C), total cholesterol (TC), hip circumference (HIP), waist circumference (WC), waist to hip ratio (WHR), body mass index (BMI), systolic blood pressure (SBP), diastolic blood pressure (DBP), hypertension (HTN), coronary artery disease (CAD), eGFR creatinine diabetes mellitus (eGFRcrea_DM), eGFR creatinine non diabetes mellitus (eGFRcrea_nonDM), eGFR creatinine all (eGFRcrea_overall), eGFR cystatin C all (eGFRcys_overall), chronic kidney disease (CKD), white blood cell counts (WBC), intercellular adhesion molecule 1 (ICAM-1), C-reactive protein (CRP), plasminogen activator inhibitor 1 (PAI-1), GT – glycaemic traits.

For HOMA-IR, we saw significant correlations with FI (r_g_=0.98, P=<0.001), FG (r_g_=0.49, P=1.37×10^-9^), HbA1_C_ (r_g_=0.28, P=0.02) and T2D (r_g_=0.56, P=2.31×10^-9^), in accordance with epidemiological observations. The genetic correlation between HOMA-B from our largest to date GWAS confirmed initial observations^18^ on the strong genetic correlation with FI (r_g_=0.80, P=8.77×10^-72^). In accordance with previous reports, the correlation with FG (r_g_=-0.32, P=0.05) and relationship with T2D (r_g_=0.05, P=0.71) were not significant after multiple testing corrections^18,52^. Other indices of glucose homeostasis, including the insulin sensitivity index, ISI, without adjustment for BMI, were inversely correlated with both HOMA indices (r_gHOMA-B/-IR_=-0.90/-1.04, P_HOMA-B/-IR_=1.20×10^-5^/2.71×10^-8^). We did not find significant genetic correlations between HOMA-B/-IR and other glycaemic traits, which is likely due to small GWAS sample sizes for those phenotypes (**Table S8**). Triglycerides (TG) and all obesity traits were directly correlated with both HOMA-B/-IR, similarly to a previous report that used enrichment analysis approaches^13^. We found no significant genetic correlations between HOMA-B/-IR and traits in the domain of T2D complications with the exception of Estimated Glomerular Filtration Rate (eGFR) defined from cystatine C and HOMA-B (r_gHOMA-B/IR_=-0.31/-0.27, P_HOMA-B/IR_=0.03/0.05).

### Effects of fasting glycaemic trait loci on PAI-1 variability

We followed up the observation from genome-wide genetic correlation estimates by evaluation of the effects of FG/FI loci on PAI-1 and HOMA-B and HOMA-IR, respectively (**Figure 4**). While only four FG loci (*PPP1R2B, MTNR1B, CRY2* and *GCKR*) showed at least nominally significant effects on PAI-1^ref39^, the direction of their effects was variable. Interestingly, among FI loci, at least half of the loci had nominal association with PAI-1. At only two loci (*PELO* and *GCKR*), FI increasing allele was associated with lower PAI-1 levels, *GCKR* variant effect being in line with its established mutational mechanism leading at the same time to higher FG and lower triglycerides levels^51^. At the same time, among four established PAI-1 loci, rs1801282-A (coding SNP, Pro12Ala) at *PPARG* is an established T2D risk^53^ and insulin resistance^29^ variant leading to higher PAI-1 levels.

**Figure 4.**
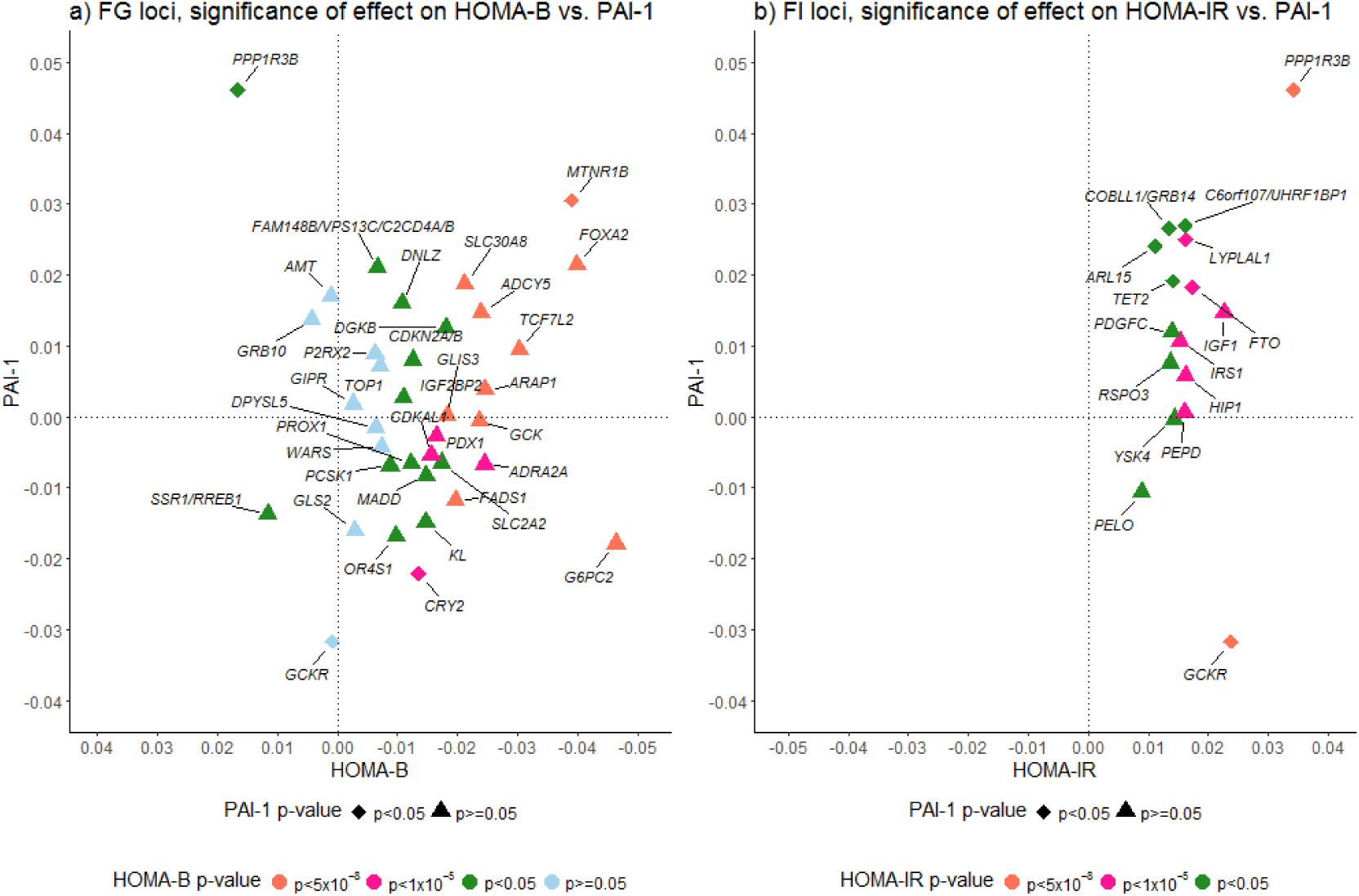
Effects of established FG/FI (a,b) loci onPAI-1 levels and HOMA-B (a)/HOMA-IR (b). The X-axis represents the direction of loci effects on PAI-1, while the Y-axis shows effects β-cell function and insulin resistance as reflected by HOMA-B/-IR, respectively.

## Discussion

We report the largest GWAS to date of HOMA-B and HOMA-IR indices in non-diabetic individuals of European origin, inferred analytically from FG and FI GWAS meta-analysis summary statistics and imputed using novel methodology^34^ to 1000 Genomes Project variant density. The application of GWIS methodology brought the total count of HOMA-B/-IR loci to 11/5, respectively, and brought the analytical sample size up to ∼75,240 individuals. This GWIS-based analysis revealed one novel HOMA-B and three novel HOMA-IR loci. We highlighted the strongest novel positive genetic correlations between non-glycaemic trait PAI-1 and HOMA-B/-IR. We also reported novel positive genetic relationships between CRP and HOMA-B/-IR, as well as inverse relationships between adiponectin and HOMA-IR. We demonstrated that GWIS of both HOMAs produced more precise SNP effect parameter estimates, and thus gained power, compared to the previous GWAS^21^. The results of LDSC^33^ analysis confirmed that the gain in power in GWIS did not arise from the introduction of spurious inflation due to population stratification or other biases.

Novel HOMA-B/-IR loci, including *FOXA2, LYPLAL1/SLC30A10, PPP1R3B*, are established for FG or FI, while HOMA-IR locus at *PER4* encoding for Period Circadian Clock 3 Pseudogene is novel. The lead variant rs10224545 at *PER4* also showed a genome-wide significant association with FI in our data. This region was not covered on Illumina Metabochip and therefore was not detected by a larger study from the MAGIC investigators^13^. Similarly to *MTNR1B*^23,54^ and *CRY2*^21^ loci, both affecting FG levels and related to indices of β-cell function, the association with insulin resistance at *PER4* gene provides an additional link between circadian rhythmicity and energy metabolism^55^. Overall, nine HOMA-B loci at *ADCY5, DGKB, GCK, SLC30A8, GLIS3, TCF7L2, ARAP1, LYPLAL1/SLC30A10* and *MTNR1B* and one HOMA-IR locus at *GCKR* are also established as contributing to the risk of T2D^47^. Among the remaining, two lead variants (rs174555, rs4240624 at *FADS1/2/3* and *PPP1R3B,* respectively) are only nominally (P<0.05) associated with T2D risk^47^. Other HOMA-B loci, such as *G6PC2,* are established for glycated haemoglobin (HbA1_C_) levels, in addition to FG, or for a wide range of glycaemic, lipid, metabolomic traits, such as variants within *FADS1*^refs21,56,57^. Among novel HOMA-IR loci, *LYPLAL1/SLC30A10* is an established FI (rs2820436, R^2^=1)^ref43^ and obesity locus known for its effects on waist-to hip ratio (WHR) in women (rs2820443, R^2^=0.44)^ref58^, while associations at *PPP1R3B* are established for several glycaemic traits, including FI (rs983309, r2=0.75; rs2126259, r2=0.85), FG (rs983309, r2=0.75) and 2-hour postprandial glucose, 2hrGlu (rs11782386, r2=0.60).

High insulin and glucose levels, insulin resistance and β-cell dysfunction characterize the pathophysiology of T2D. However, the exact mechanism of genetic interrelationships between the metabolic processes and disease is still unclear. Amongst the established T2D loci^45,46,48^ we investigated in relation to HOMA indices GWIS results (**Tables S5-S7**), only 41/30 have at least nominal effect on HOMA-B/-IR variability, respectively, highlighting an incomplete overlap between genetic variants affecting normal glucose homeostasis and processes involved in T2D pathogenesis. Insulin resistance and β-cell function may have distinct impact on susceptibility to T2D^28^, and mechanistically T2D loci can be related to a specific biological process affecting insulin secretion, resistance or processing^59^. Effects of T2D loci follow the main subdivisions into β-cell, hyperglycaemia, proinsulin and insulin resistance groups^29^. The group of established T2D loci that influence insulin resistance or fasting insulin has been under-characterised, most probably, due to low power of respective endophenotype GWAS meta-analyses^21,60^. Empowered by analytically inferred HOMA-IR GWAS, we demonstrated that the group on insulin resistance loci is larger than described before and comparable to that of β-cell loci. As expected from epidemiological studies, FG loci also affect HOMA-B values (**Figure S5**), whereas FI variants have most prominent effect on HOMA-IR.

We confirmed the evidence of shared genetic effects between HOMA-B and T2D risk at several loci, manifested by inverse relationships^28,29^. However, the effect of left truncation for FG levels to define non-diabetic individuals in general population leads to seemingly improved β-cell function through positive HOMA-B values for the insulin resistance loci. In fact, we did not observe any genetic correlation between HOMA-B and T2D using summary statistics from the latest GWAS meta-analyses. In parallel, we however demonstrated that HOMA-B loci, associated with a decrease in β-cell function, exert their effects by both increasing and decreasing insulin resistance (**Figure 2a-b**).

Inflammation markers PAI-1 and CRP showed strong positive genetic correlation with both HOMA-B/-IR measures. The correlation of these markers was stronger with fasting insulin resistance. CRP is a marker of increased cardiovascular risk^61,62^ and PAI-1 is associated with CAD^63^ and myocardial infarction^64^, both high risk factors in T2D. Importantly, PAI-1 is a fibrinolytic and inflammatory marker, which was proposed and confirmed in epidemiological studies as an independent risk factor for T2D^65,66^. In our study among non-glycaemic traits, PAI-1 showed the strongest genetic correlation with fasting insulin resistance, a pathophysiological process leading to T2D. Moreover, when we looked at the individual effects of FG/FI variants on PAI-1 variability, variant alleles associated with higher FI grouped around increased levels of PAI-1, while there was no clear pattern of such an effect for FG variants. From pathophysiological point of view, the clustering of hyperinsulinaemia, dysglycemia, dyslipidemia, and hypertension as cardiovascular risk endophenotypes in T2D occurs around insulin resistance, and in the presence of elevated PAI-1 levels, leads to fibrinolytic dysfunction, increased thrombotic risk^17,67^, and induces local or systemic low-grade inflammation. Taken together, evidence from our study provides novel genetic support for the need to dissect in better detail these deficient mechanisms in the early pathogenesis of T2D.

Noteworthy, we observed no significant genetic correlation between HOMA-B/-IR and CAD, hypertension, SBP and DBP, which could indicate that the associations between HOMA-B/-IR and PAI-1 and CRP are strictly due to inflammatory pathways involved in T2D risk. Contrary to expectations from epidemiological studies, we neither observed correlation between adiponectin and HOMA-B for genetic effects using the GWAS summary statistics, whilst the respective genetic correlation with HOMA-IR may be through its association with obesity, since adiponectin is secreted in adipose tissue. Alternatively, it can be due to its involvement in inflammatory processes in T2D pathogenesis^68^. Analysis of CKD and its marker eGFR based on creatinine did not yield statistically significant genetic correlations; only eGFR based on cystatine C for HOMA-B did so. The latter could reflect the association between cystatine and metabolic syndrome^69^, whose definition includes T2D risk factors such as insulin resistance and abnormal fat distribution^70^.

Finally, we reported novel significant positive genetic correlation between HOMA-IR and T2D, and our estimates of genetic correlations are concordant with established genome-wide relationships between HOMA-B and FG/FI/BMI/T2D and HOMA-IR and FG/FI/BMI^18^. The positive genetic correlations for TG and negative for high density lipoprotein cholesterol (HDL-C) with HOMA-B/-IR reflect the conditions of diabetic dyslipidaemia, characterised by increased levels of TG and decreased levels of HDL cholesterol^71^. The role of obesity and adiposity^43,72^, in particular, in the risk of T2D is well established and was reflected in significant genetic correlations between HOMA-B/-IR and obesity related traits, such as BMI, WHR, waist circumference (WC), hip circumference (HIP), in our study. In addition, adiposity loci *FTO, MC4R* and *NRXN3* were at least nominally associated with HOMA-B/-IR in our analysis not adjusted for BMI.

Our study provides additional evidence for the GWIS method as a powerful tool for future GWAS studies of analytically derived phenotypes. Our work also suggests that GWAS meta-analysis of summary statistics is a useful source of information for follow-up analyses, including inferences about genetic correlations and mechanistic characterisation of the specific trait pathophysiology through their effects on other related phenotypes.

## Data Access

The GWAS summary statistics for the MAGIC investigators study of FG and FI and the present study results will be deposited on the consortium web site www.magicinvestigators.org.

## Acknowledgments

IOF is supported by Biobanking and Biomolecular Resources Research Infrastructure (BBMRI-NL, NWO 184.021.007). I.P. is funded by the World Cancer Research Fund (WCRF UK) and World Cancer Research Fund International (2017/1641), the Wellcome Trust (WT205915), and the European Union’s Horizon 2020 research and innovation programme (DYNAhealth, project number 633595). MGN is supported by Royal Netherlands Academy of Science Professor Award (PAH/6635) granted to D.I. Boomsma. Mike A. Nalls’ participation is supported by a consulting contract between Data Tecnica International and the National Institute on Aging, NIH, Bethesda, MD, USA. Data on glycaemic traits have been contributed by MAGIC investigators and have been downloaded from www.magicinvestigators.org.

## Author contributions

Conceived and designed the experiments: I.P. and D.I.B.; Analysed the data: Meta-analyses: R.M., D.I.C., S.G., J.H., M.A.N., C.J.O., G.P., P.M.R., MAGIC Investigators and CHARGE Inflammation working group; analytical derivation of HOMA-B/HOMA-IR GWIS summary statistics: I.O.F, M.G.N; central analysis group: I.O.F, L.Z, M.K., M.G.N, I.P., and XC-Pleiotropy Group. Wrote the first draft of the manuscript: I.O.F., J.J.H., M.G.N, I.P. and D.I.B. Contributed to the writing of the manuscript: I.O.F., M.G.N., J.J.H., L.Z, Z.B., D.I.C., S.G., J.H., M.A.N., C.J.O., G.P., P.M.R., M.K., R.M., I.P., D.I.B..

