## Supplementary Materials and Methods for "Genetics of fasting indices of glucose homeostasis using GWIS unravels tight relationships with inflammatory markers"

*MAGIC studies (subset) contributing to each primary phenotype meta-analysis*

*FG:* AGES, ALSPAC, AMISH, ARIC, ASCOT, BLSA, BSN, CHS, COLAUS, CROATIA, DECODE, DGI, ERF, FAMHS, FENLAND, FHS, FRENCHADULTCONTROL, FRENCHADULTOBESE, FRENCHYOUNGCONTROL, FRENCHYOUNGOBESE, FUSION, GENOA, GENOMEUTWINS, HABC, HEALTH2000, INCHIANTI, KORA, KORCULA, LIFELINES, NFBC66, ORCADES, PREVEND, PROCARDIS, RS, SARDINIA, SORBS, SPLIT, SUVIMAX, TWINSUK, TYROL.

*FI:* AGES, ALSPAC, AMISH, ARIC, BLSA, BSN, CHS, COLAUS, CROATIA, DECODE, DGI, ERF, FAMHS, FENLAND, FHS, FRENCHADULTCONTROL, FRENCHADULTOBESE, FRENCHYOUNGOBESE, FUSION, GENOA, GENOMEUTWINS, HABC, HEALTH2000, INCHIANTI, KORCULA, NFBC66, ORCADES, PREVEND, PROCARDIS, RS, SARDINIA, SORBS, TWINSUK.

*Fasting Glucose (FG) and Fasting Insulin (FI) GWAS and pre-GWIS quality control (QC)*

Summary statistics for FG and FI GWAS meta-analyses were obtained with permission of MAGIC (the Meta-Analyses of Glucose and Insulin-related traits Consortium). For detailed description of the above-mentioned cohorts, genotyping, imputation and QC on the individual study level see (*Lagou et al., manuscript in preparation, www.magicinvestigators.org*). In the individual samples FG and FI were measured in mmol/L and pmol/L respectively. Before performing a GWAS FI was natural logarithm transformed. Genotyping was performed on either Illumina or Affymetrix platforms and imputed to the HapMap II CEU reference panel[1]. Studies, genotyped on specifically designed arrays, such as Metabochip were excluded. Meta-analysis was performed in the software GWAMA[2].

We started with 2,728,451 SNPs for FG and 2,717,287 SNPs for FI, which passed QC in the separate GWAS. First, we compared allele frequencies between HapMap II CEU[1] and separate FG and FG analysis results. SNPs with allele frequency differences more than 20% were excluded from each study results (66 for FG and 36 for FI). Second, we selected 2,715,654 overlapped SNPs and checked them for reference allele concordance and frequency. 2,715,004 SNPs had concordant alleles and 647 SNPs had swapped reference and not reference allele. We aligned 647 alleles, their beta coefficients and frequencies. 3 SNPs were flipped by strand, we excluded them from the analysis. SNPs, which had an allele frequency difference between the GWASs of FG and FI of more than 10% were excluded (N = 4,299). The resulting set contained 2,711,352 SNPs. Data management and analyses were performed in R version 3.2.5[3], easyQC software[4] was used to compare FG/FI SNP frequencies with HapMap II CEU reference set[1].

*GWIS of HOMA-B and HOMA-IR*

Throughout the Supplementary Materials and Methods we will refer to the previous meta-analysis of HOMA-B/-IR[5] as 'published' and the results of the present study and analyses as to 'inferred' meta-analysis of HOMA-B/-IR.

*Allele effect estimates*

The GWIS-inferred[6] effect estimates (βs) for HOMA-B and HOMA-IR for $SNP_{i}$, $i$=1,…,M were calculated using the following formulae:

$$\beta_{HOMA-B/-IR} = AF1 \times \beta 1 + AF2 \times \beta 2 \,, [1]$$

AF1 and AF2 are relative population frequencies and β1 and β2 are HOMA-B/-IR functions (main text formulas [3,4]) evaluated for 1 and 2 copies of effect allele respectively. For both HOMA-B/-IR AF1 and AF2 are the same:

$$AF1 = \frac{2\times p\times(1-p)}{2\times p\times(1-p)+p^2}, [2]$$

$$AF2 = \frac{p^2}{2\times p\times(1-p)+p^2}, [3]$$

where $p$ is the effect allele frequency, calculated as a weighted mean of FG and FI effect allele frequencies, expressed as:

$$p = \frac{N1\times EAF_{FI}+N2\times EAF_{FG}}{N1+N2}, [4]$$

where N1 and N2 are the sample sizes of FI and FG respectively for $SNP_i$.

For HOMA-B β1 and β2 for each $SNP_i$ are represented by:

$$\beta 1 = ln\frac{20\times\frac{e^{(M_{FI}+\beta_{FI})}}{6.945}}{(M_{FG}+\beta_{FG})-3.5} - ln\frac{20\times\frac{e^{M_{FI}}}{6.945}}{M_{FG}-3.5}, [5]$$

$$\beta 2 = (ln\frac{20\times\frac{e^{(M_{FI}+2\times\beta_{FI})}}{6.945}}{(M_{FG}+2\times\beta_{FG})-3.5} - ln\frac{20\times\frac{e^{M_{FI}}}{6.945}}{M_{FG}-3.5})/2, [6]$$

For HOMA-IR β1 and β2 for each $SNP_i$ are represented by:

$$\beta 1 = ln\frac{(M_{FG}+\beta_{FG})\times\frac{e^{(M_{FI}+\beta_{FI})}}{6.945}}{22.5} - ln\frac{M_{FG}\times\frac{e^{M_{FI}}}{6.945}}{22.5}, [7]$$

$$\beta 2 = (ln\frac{(M_{FG}+2\times\beta_{FG})\times\frac{e^{(M_{FI}+2\times\beta_{FI})}}{6.945}}{22.5} - ln\frac{M_{FG}\times\frac{e^{M_{FI}}}{6.945}}{22.5})/2, [8]$$

$M_{FG/FI}$ are the estimated population means (5.252857 mmol/l and 60.40195 pmol/l for FG/FI respectively) computed as mean of means across meta-analyses studies (from which mean estimates were available). $M_{FG/FI}$ are used instead of intercept terms in GWIS[6]. $\beta_{FG/FI}$ are the effect estimates for a $SNP_i$ from GWAS meta-analyses of FG and FI respectively. During calculation of β values 35 SNPs resulted in missing values (NAs) for HOMA-B because the logarithm of negative value is not defined.

*Standard errors*

For the calculation of the standard errors (SEs) of the GWIS-inferred[6] summary statistics of HOMA-B/-IR we used the Delta method implemented in R package *msm* to account for the estimated sample overlap[7]. We estimated the sample overlap between FG and FI by using two strategies: 1) 'Global', assuming the sample overlap is the same across all SNPs. In this case cross-trait LDSC[8] intercept from FG and FI genetic correlation analysis (before the GWIS QC) reflects the sample overlap between FG and FI meta-analyses and could be used as a correction factor while calculating SEs. 2) 'Local', assuming varying sample overlap across SNPs. To compute the cross-trait LDSC[8] intercept per each SNP we performed the following steps:

a) We run the LDSC[8] between FG and FI to obtain the LDSC intercept. The latter is represented by the formulae

$$gcov\_int = \frac{N_S \times r}{\sqrt{N_1 \times N_2}}, [9]$$

where $N_s$=samples overlap, *r* is the phenotypic correlation between the two traits and $N_1$ and $N_2$ are the sample sizes for traits 1 and 2 (here FI and FG) from each meta-analysis. The same estimate of *gcov_int* was used as correction factor when assuming 'global' sample overlap (*gcov_int* = 0.2606).

b) In order to compute 'local' $gcov\_int_i$ for each $SNP_i$ using formula [9] we would need to know $Ns_i$, $N1_i$ and $N2_i$ per each $SNP_i$ as well as phenotypic correlation between FG and FI (*r)*. $N1_i$ and $N2_i$ are known. $Ns_i$ is the unknown sample overlap for each $SNP_i$ and therefore we approximated it as min($N1_i$,$N2_i$).

As only summary statistics were available and we were not able to estimate phenotypic correlation *r* from the actual phenotype data, we rearranged formula [9] to derive *r* from LDSC intercept as follows:

$$r = gcov\_int \times \sqrt{N_1 \times N_2}/N_s \text{ , [10]}$$

where *gcov_int* is the cross-trait LDSC intercept from step a), $N_1$=*minimum* sample size for FI, and $N_2$=*minimum* sample size for FG meta-analyses across all SNPs. Note, $N_1$ and $N_2$ were obtained after an initial run with LDSC[8], which removes SNPs with low N and suggests reliable minimum values for the traits ($N_1$<42727.3333333, $N_2$<58876.6666667). As sample overlap between FG and FI is unknown, we approximated $N_s$=min($N_1$,$N_2$),

c) Finally, we calculated a vector of cross-trait LDSC intercepts $gcov\_int_i$ per $SNP_i$ *i*=1,…,M using the same formulae [9] adjusted for each $SNP_i$ as follows:

$$gcov\_int_i = \frac{Ns_i \times r}{\sqrt{N1_i \times N2_i}}, \text{ [11],}$$

where we used the estimated phenotypic correlation *r* from the previous step b). The values of computed cross-trait intercept vector ranged from 0.02846 to 0.3059 with the mean=0.2634 and median=0.2606.

*Sample size*

We approximated a sample size for each $SNP_i$ of the inferred HOMA-B/-IR meta-analysis results, as a geometric mean between FG (N1) and FI (N2) sample sizes:

$$N = \sqrt{N1 \times N2}, \text{ [12]}$$

*Post-GWIS HOMA-B/-IR QC*

We computed the genetic correlation $r_g$ between published and inferred HOMA-B/-IR (both ‘global’ or ‘local’ corrected versions) using LDSC[8] (the expectation, when the same trait is analyzed, is that the genetic correlation equals 1) and found similar results (**Table S9**). The comparison of summary statistics distributions between published and inferred HOMAs demonstrated that standard errors (SEs) decreased in current analyses compared to the previously determined validating the fact that GWIS results gain in power over previous GWAS efforts (**Figures S8 – S18**)[5]. We also compared the distribution of summary statistics of the two versions of inferred and published HOMA-B/-IR aiming to select 'global' or 'local' corrected version. The distribution between the 'global' and 'local' correction versions of HOMA-B/-IR results were also very similar (**Figures S8 – S19**), therefore we proceeded with the 'local' correction as it is assumed to be more precise. However, to re-iterate we did not observe a difference between 'global' and 'local' correction approaches in current analysis and given the similarity of summary statistics they both could have been used in further analysis (**Figure S19**).

From genetic correlation analysis the LDSC intercepts for inferred HOMA-B/-IR summary statistics were also available (**Table S9**). The LDSC intercepts were estimated as 1.0493/1.033 for HOMA-B/-IR 'local' corrections, confirming the gain in power in GWIS did not arise from inflation due to imprecise approximation. However, as LDSC intercepts exceeded 1 and could indicate residual population stratification or misspecification of the sample overlap, we additionally adjusted SEs for excess in the LDSC intercept:

$$SE_{adj} = \frac{\sqrt{(SE \times \sqrt{N})^2 \times intercept}}{\sqrt{N}}, [13]$$

where for a particular $SNP_i$ SE is the standard error from inferred HOMA-B/-IR summary statistics, N is the sample size and $(SE \times \sqrt{N})^2$ is the variance. P-values were computed based on chi-square distribution with 1 degree of freedom.

The comparison of variants with low N and MAF between the published and inferred GWAS showed a marginally lower concordance between SEs for low N and MAF (**Figures S9, S13-S15, S18**) confirming it is sensible to apply QC filters for minimal sample size and MAF. As difference in SEs between inferred and published meta-analysis results gets larger when N decreases (**Figures S13-S15,S18**) we excluded SNPs with low N (278,542/278,577 SNPs with N<35,000 for HOMA-B/-IR, among which 113/113 were filtered out based on MAF<0.01 filter). The total number of SNPs after QC was 2,432,775 for HOMA-B/-IR. LocusZoom[9] was used to build region plots for the established loci (**Figures S20-S21**).

*Effects of established T2D, FG/FI and $HbA1_C$ loci on HOMA-B/-IR*

We plotted the effects of HOMA-B/-IR at established T2D, FG/FI and $HbA1_C$ loci in Figure 2. Effects of HOMA-B/-IR SNPs were aligned to the risk allele of either T2D, FG/FI or $HbA1_C$ SNPs. If loci between T2D and FG/FI overlapped, but SNPs did not, we selected one SNP per locus. First, we considered loci from previous efforts[10,11], then added novel 1000G detected loci from the two recent T2D studies[12,13] and $HbA1_C$[14]. For SNPs, which were not present in HOMA-B/-IR results, we obtained the proxies via LDlink (R2 > 0.9)[15].

**Table S1.** Comparison of HOMA-B effects across genome-wide significant loci between inferred and published GWAS

| SNP | CHR | Locus name | HOMA-B inferred | | | HOMA-B published | |
|---|---|---|---|---|---|---|---|
| | | | Alleles (effect / other) | Effect (SE), p-value | N | Alleles (effect / other) | Effect (SE), p-value |
| rs560887 | 2 | *G6PC2* | C/T | -0.046 (0.0033), $4.74\times10^{-46}$ | 75,173 | T/C | 0.040 (0.0036), $7.67\times10^{-29}$ |
| rs11708067 | 3 | *ADCY5* | A/G | -0.024 (0.0038), $2.50\times10^{-10}$ | 75,227 | A/G | -0.016 (0.0043), $1.77\times10^{-04}$ |
| rs10258074 | 7 | *DGKB* | A/T | 0.019 (0.0031), $2.55\times10^{-10}$ | 75,202 | A/T | 0.022 (0.0033), $2.99\times10^{-11}$ |
| rs3757840 | 7 | *GCK* | G/T | 0.02 (0.0033), $1.98\times10^{-09}$ | 64,352 | T/G | -0.017 (0.0038), $7.96\times10^{-06}$ |
| rs3802177 | 8 | *SLC30A8* | G/A | -0.021 (0.0036), $2.55\times10^{-09}$ | 68,044 | A/G | 0.016 (0.0038), $1.96\times10^{-05}$ |
| rs7034200 | 9 | *GLIS3* | A/C | -0.019 (0.003), $2.96\times10^{-10}$ | 74,669 | A/C | -0.016 (0.0033), $1.67\times10^{-06}$ |
| rs7903146 | 10 | *TCF7L2* | C/T | 0.03 (0.0034), $7.41\times10^{-19}$ | 75,212 | T/C | -0.020 (0.0038), $1.39\times10^{-07}$ |
| rs10830963 | 11 | *MTNR1B* | C/G | 0.039 (0.0038), $9.34\times10^{-25}$ | 68,612 | C/G | 0.039 (0.0040), $8.60\times10^{-23}$ |
| rs1552224 | 11 | *ARAP1* | C/A | 0.025 (0.004), $4.28\times10^{-10}$ | 75,207 | A/C | -0.017 (0.0043), $9.39\times10^{-05}$ |
| rs174555 | 11 | *FADS1-2-3* | T/C | -0.021 (0.0033), $8.97\times10^{-11}$ | 75,237 | T/C | -0.015 (0.0034), $1.32\times10^{-05}$ |
| rs5029909 | 20 | *FOXA2* | C/T | -0.044 (0.0079), $2.70\times10^{-08}$ | 68,584 | T/C | 0.032 (0.0092), $4.84\times10^{-04}$ |

**Table S2.** Comparison of HOMA-IR effects across genome-wide significant loci between inferred and published GWAS

| | | | HOMA-IR inferred | | | HOMA-IR published | |
|---|---|---|---|---|---|---|---|
| SNP | CHR | Locus name | Alleles (effect / other) | Effect (SE), p-value | N | Alleles (effect / other) | Effect (SE), p-value |
| rs2605101 | 1 | *LYPLAL1/ SLC30A10* | A/T | 0.019 (0.0034), $3.78{\times}10^{-08}$ | 75,238 | A/T | 0.016 (0.0042), $1.24{\times}10^{-04}$ |
| rs780093 | 2 | *GCKR* | C/T | 0.024 (0.0032), $1.06{\times}10^{-13}$ | 75,226 | T/C | -0.019 (0.0040), $1.31{\times}10^{-06}$ |
| rs10224545 | 7 | *PER4* | T/C | -0.031 (0.0056), $2.60{\times}10^{-08}$ | 71,951 | T/C | -0.015 (0.0069), $2.51{\times}10^{-02}$ |
| rs4240624 | 8 | *PPP1R3B* | A/G | -0.035 (0.0055), $2.73{\times}10^{-10}$ | 68,617 | A/G | -0.021 (0.0066), $1.66{\times}10^{-03}$ |
| rs2114912 | 12 | *IGF1* | T/G | 0.026 (0.0044), $3.08{\times}10^{-09}$ | 74,634 | T/G | 0.026 (0.0053), $1.33{\times}10^{-06}$ |

**Table S3.** Meta-analyses summary statistics sources

| | Reference | Year | Phenotype | Sample Size | Ancestry |
|---|---|---|---|---|---|
| 1 | Pattaro, C., Teumer, A., Gorski, M., Chu, A. Y., Li, M., Mijatovic, V., ... & Taliun, D. (2016). Genetic associations at 53 loci highlight cell types and biological pathways relevant for kidney function. Nature communications, 7. | 2015 | Estimated glomerular filtration rate based on serum creatinine (eGFRcrea) all/non-DM/DM, cystatin C (eGFRcys) all, chronic kidney disease (CKD) all | 133,413 (eGFRcrea all), 118,448 (eGFRcrea non-DM), 11,522 (eGFRcrea DM), 32,834 (eGFRcys), 117,165 (CKD) | European |
| 2 | Dehghan, A., Dupuis, J., Barbalic, M., Bis, J. C., Eiriksdottir, G., Lu, C., ... & Baumert, J. (2011). Meta-analysis of genome-wide association studies in> 80 000 subjects identifies multiple loci for C-reactive protein levels. Circulation, 123(7), 731-738. | 2010 | C-Reactive Protein (CRP) | 66,185 | European |
| 3 | Huang, J., Sabater-Lleal, M., Asselbergs, F. W., Tregouet, D., Shin, S. Y., Ding, J., ... & Smith, N. L. (2012). Genome-wide association study for circulating levels of PAI-1 provides novel insights into its regulation. Blood, 120(24), 4873-4881. | 2012 | Plasminogen Activator Inhibitor 1 (PAI-1) | 19,599 | European |
| 4 | Dastani, Z., Hivert, M. F., Timpson, N., Perry, J. R., Yuan, X., Scott, R. A., ... & Fuchsberger, C. (2012). Novel loci for adiponectin levels and their influence on type 2 diabetes and metabolic traits: a multi-ethnic meta-analysis of 45,891 individuals. PLoS Genet, 8(3), e1002607. | 2012 | Adiponectin (ADIP) | 29,347 | European |
| 5 | Paré, G., Ridker, P. M., Rose, L., Barbalic, M., Dupuis, J., Dehghan, A., ... & Chasman, D. I. (2011). Genome-wide association analysis of soluble ICAM-1 concentration reveals novel associations at the NFKBIK, PNPLA3, RELA, and SH2B3 loci. PLoS Genet, 7(4), e1001374. | 2011 | Intercellular Adhesion molecule 1 (ICAM-1) | 22,435 | North American women, Caucasian |
| 6 | Nalls, M. A., Couper, D. J., Tanaka, T., Van Rooij, F. J., Chen, M. H., Smith, A. V., ... & Wood, A. R. (2011). Multiple loci are associated with white blood cell phenotypes. PLoS Genet, 7(6), e1002113. | 2011 | White Blood Cell Counts (WBC) | 19,509 | European |

**Table S3.** Continued

| | Reference | Year | Phenotype | Sample Size | Ancestry |
|---|---|---|---|---|---|
| 7 | Schunkert, H., König, I. R., Kathiresan, S., Reilly, M. P., Assimes, T. L., Holm, H., ... & Absher, D. (2011). Large-scale association analysis identifies 13 new susceptibility loci for coronary artery disease. *Nature genetics*, *43*(4), 333-338. | 2011 | Coronary Artery Disease (CAD) | 22,233 cases and 64,762 controls | European |
| 8 | Prokopenko, I., Poon, W., Mägi, R., Prasad, R., Salehi, S. A., Almgren, P., ... & Stančáková, A. (2014). A central role for GRB10 in regulation of islet function in man. PLoS Genet, 10(4), e1004235. | 2014 | Corrected Insulin Response (CIR), ratio of the area under the curve for insulin and glucose (AUCins/AUCglu), disposition index (DI), insulin at 30 min (Ins30), incremental insulin at 30 min (Increm30), insulin response adjusted for BMI (Ins30adjBMI), area under the curve of insulin levels during OGTT (AUCIns). | 5,318 CIR, 4,789 CIRadjISI, 4,213 AUCins/AUCglu, 5,130 DI, 4,483 Ins30, 4,447 Increm30, 4,483 Ins30adjBMI, 4,324 AUCIns. | European |
| 9 | Strawbridge, R. J., Dupuis, J., Prokopenko, I., Barker, A., Ahlqvist, E., Rybin, D., ... & Nica, A. (2011). Genome-wide association identifies nine common variants associated with fasting proinsulin levels and provides new insights into the pathophysiology of type 2 diabetes. Diabetes, 60(10), 2624-2634. | 2011 | Fasting Proinsulin | 10,701 | European |
| 10 | Soranzo, N., Sanna, S., Wheeler, E., Gieger, C., Radke, D., Dupuis, J., ... & Sandhu, M. S. (2010). Common variants at 10 genomic loci influence hemoglobin A1C levels via glycemic and nonglycemic pathways. Diabetes, 59(12), 3229-3239. | 2010 | Hemoglobin A1C | 46,368 | European |
| 11 | Saxena, R., Hivert, M. F., Langenberg, C., Tanaka, T., Pankow, J. S., Vollenweider, P., ... & Kao, W. L. (2010). Genetic variation in GIPR influences the glucose and insulin responses to an oral glucose challenge. Nature genetics, 42(2), 142-148. | 2009 | 2hr glucose | 15,234 | European |

**Table S3.** Continued

| | Reference | Year | Phenotype | Sample Size | Ancestry |
|---|---|---|---|---|---|
| 12 | Newton-Cheh, C., Johnson, T., Gateva, V., Tobin, M. D., Bochud, M., Coin, L., ... & Papadakis, K. (2009). Genome-wide association study identifies eight loci associated with blood pressure. Nature genetics, 41(6), 666-676. | 2008 | Systolic and Diastolic blood pressure (SBP/DBP) | 34,433 | European |
| 13 | Shungin, D., Winkler, T. W., Croteau-Chonka, D. C., Ferreira, T., Locke, A. E., Mägi, R., ... & Workalemahu, T. (2015). New genetic loci link adipose and insulin biology to body fat distribution. Nature, 518(7538), 187-196. | 2015 | HIP, WHR, WC | ≈ 210,088 | European |
| 14 | Morris, A. P., Voight, B. F., Teslovich, T. M., Ferreira, T., Segre, A. V., Steinthorsdottir, V., ... & Prokopenko, I. (2012). Large-scale association analysis provides insights into the genetic architecture and pathophysiology of type 2 diabetes. Nature genetics, 44(9), 981. | 2012 | T2D | 12,171 cases and 56,862 controls | European |
| 15 | Walford, G. A., Gustafsson, S., Rybin, D., Stančáková, A., Chen, H., Liu, C. T., ... & Mägi, R. (2016). Genome-wide association study of the modified Stumvoll Insulin Sensitivity Index identifies BCL2 and FAM19A2 as novel insulin sensitivity loci. Diabetes, db160199. | 2016 | ISI | 16,753 | European |
| 16 | Teslovich, T. M., Musunuru, K., Smith, A. V., Edmondson, A. C., Stylianou, I. M., Koseki, M., ... & Johansen, C. T. (2010). Biological, clinical and population relevance of 95 loci for blood lipids. Nature, 466(7307), 707-713. | 2010 | TG, HDL-C, LDL-C, TC | Maximum sample size 100,184 for TC, 95,454 for LDL-C, 99,900 for HDL-C and 96,598 for TG | European |
| 17 | Locke, A. E., Kahali, B., Berndt, S. I., Justice, A. E., Pers, T. H., Day, F. R., ... & Croteau-Chonka, D. C. (2015). Genetic studies of body mass index yield new insights for obesity biology. Nature, 518(7538), 197-206. | 2015 | BMI | 234,069 | European |
| 18 | Shungin, D., Winkler, T. W., Croteau-Chonka, D. C., Ferreira, T., Locke, A. E., Mägi, R., ... & Workalemahu, T. (2015). New genetic loci link adipose and insulin biology to body fat distribution. Nature, 518(7538), 187-196. | 2015 | WHR, WC, HIP | ≈ 210,088 | European |

**Table S4.** 1000G summary statistics imputation results

| SNP | CHR | Locus name | Alleles (effect / other) | Effect (SE), p-value | N | P-value (imputed) | New lead variant (imputed) | P-value of the new lead variant | LD with new lead variant, $R^2$ | Distance from lead variant, bp |
|---|---|---|---|---|---|---|---|---|---|---|
| | | | | | HOMA-B | | | | | |
| rs560887 | 2 | *G6PC2* | C/T | -0.046 (0.0033), $4.74\times10^{-46}$ | 75,173 | $4.74\times10^{-46}$ | rs580670 | $2.35\times10^{-47}$ | 0.7009 | 12,973 |
| rs11708067 | 3 | *ADCY5* | A/G | -0.024 (0.0038), $2.50\times10^{-10}$ | 75,227 | $2.50\times10^{-10}$ | - | - | - | - |
| rs10258074 | 7 | *DGKB* | A/T | 0.019 (0.0031), $2.55\times10^{-10}$ | 75,202 | $2.04\times10^{-09}$ | rs2191349 | $5.41\times10^{-10}$ | 0.9881 | 93 |
| rs3757840 | 7 | *GCK* | G/T | 0.02 (0.0033), $1.98\times10^{-09}$ | 64,352 | $1.98\times10^{-09}$ | - | - | - | - |
| rs3802177 | 8 | *SLC30A8* | G/A | -0.021 (0.0036), $2.55\times10^{-09}$ | 68,044 | $2.55\times10^{-09}$ | rs35859536 | $9.48\times10^{-10}$ | 0.9854 | 6,450 |
| rs7034200 | 9 | *GLIS3* | A/C | -0.019 (0.003), $2.96\times10^{-10}$ | 74,669 | $2.96\times10^{-10}$ | rs57884925 | $2.85\times10^{-10}$ | 0.9685 | 3,931 |
| rs7903146 | 10 | *TCF7L2* | C/T | 0.03 (0.0034), $7.41\times10^{-19}$ | 75,212 | $7.41\times10^{-19}$ | rs10659211 | $6.76\times10^{-22}$ | 0.9103 | 24,232 |
| rs10830963 | 11 | *MTNR1B* | C/G | 0.039 (0.0038), $9.34\times10^{-25}$ | 68,612 | $2.73\times10^{-15}$ | rs10830956 | $5.23\times10^{-21}$ | 0.7217 | 27,697 |
| rs1552224 | 11 | *ARAP1[STAR D10]* | C/A | 0.025 (0.004), $4.28\times10^{-10}$ | 75,207 | $4.28\times10^{-10}$ | rs11603349 | $5.82\times10^{-11}$ | 0.9313 | 27,596 |
| rs174555 | 11 | *FADS1/2/3* | T/C | -0.021 (0.0033), $8.97\times10^{-11}$ | 75,237 | $8.97\times10^{-11}$ | - | - | - | - |
| rs5029909 | 20 | *FOXA2* | C/T | -0.044 (0.0079), $2.70\times10^{-08}$ | 68,584 | $2.70\times10^{-08}$ | - | - | - | - |

**Table S4.** Continued

| SNP | CHR | Locus name | Alleles (effect / other) | Effect (SE), p-value | N | P-value (imputed) | New lead variant (imputed) | P-value of the new lead variant | LD with new lead variant, $R^2$ | Distance from lead variant, bp |
|---|---|---|---|---|---|---|---|---|---|---|
| | | | | | HOMA-IR | | | | | |
| rs2605101 | 1 | *LYPLAL1/ SLC30A10* | A/T | 0.019 (0.0034), $3.78\times10^{-08}$ | 75,238 | $3.46\times10^{-07}$ | rs6669360 | $5.41\times10^{-08}$ | 0.5529 | 3,443 |
| rs780093 | 2 | *GCKR* | C/T | 0.024 (0.0032), $1.06\times10^{-13}$ | 75,226 | $1.06\times10^{-13}$ | rs11336847 | $3.08\times10^{-14}$ | 0.9796 | 6,390 |
| rs10224545 | 7 | *PER4* | T/C | -0.031 (0.0056), $2.60\times10^{-08}$ | 71,951 | $2.60\times10^{-08}$ | - | - | - | - |
| rs4240624 | 8 | *PPP1R3B* | A/G | -0.035 (0.0055), $2.73\times10^{-10}$ | 68,617 | $2.73\times10^{-10}$ | - | - | - | - |
| rs2114912 | 12 | *IGF1* | T/G | 0.026 (0.0044), $3.08\times10^{-09}$ | 74,634 | $3.08\times10^{-09}$ | - | - | - | - |

**Table S5.** Established T2D loci with their effect, standard errors (SE), significance (P), sample size (N) and minor allele frequency (MAF) in HOMA-B/-IR GWIS.

| rsID | LOCUS | EA | NEA | HOMA-B, effect (SE, P) | HOMA-IR, effect (SE, P) | N | MAF |
|---|---|---|---|---|---|---|---|
| rs10146997 | *NRXN3* | G | A | 0.0108 (0.004, 3.57e-03) | 0.0128 (0.004, 9.87e-04) | 75216 | 0.21 |
| rs10203174 | *THADA* | C | T | -0.0209 (0.005, 5.43e-05) | -0.0084 (0.005, 1.24e-01) | 75198 | 0.1 |
| rs10229583 | *PAX4* | G | A | 0.0013 (0.004, 7.15e-01) | -0.0032 (0.004, 3.94e-01) | 75194 | 0.24 |
| rs10401969 | *CILP2* | C | T | -3e-04 (0.007, 9.66e-01) | 0.0099 (0.007, 1.69e-01) | 68635 | 0.07 |
| rs10507349 | *RNF6* | G | A | 0.0045 (0.004, 2.08e-01) | -0.0024 (0.004, 5.33e-01) | 75173 | 0.23 |
| rs1061810 | *HSD17B12* | A | C | 0.0029 (0.003, 3.97e-01) | 0.0059 (0.004, 1.05e-01) | 75185 | 0.29 |
| rs10758593 | *GLIS3* | A | G | -0.0183 (0.003, 1.90e-09) | -0.0072 (0.003, 2.55e-02) | 75203 | 0.42 |
| rs10811661 | *CDKN2A/B* | T | C | -0.0125 (0.004, 2.11e-03) | 0.0034 (0.004, 4.39e-01) | 68591 | 0.18 |
| rs10830963 | *MTNR1B* | G | C | -0.039 (0.004, 9.34e-25) | 0.0184 (0.004, 4.40e-06) | 68612 | 0.29 |
| rs10842994 | *KLHDC5* | C | T | -0.0025 (0.004, 5.15e-01) | 7e-04 (0.004, 8.60e-01) | 75225 | 0.21 |
| rs10886471 | *GRK5* | C | T | 0.0051 (0.003, 1.05e-01) | -2e-04 (0.003, 9.43e-01) | 75137 | 0.5 |
| rs10923931 | *NOTCH2* | T | G | -0.0053 (0.005, 2.81e-01) | 1e-04 (0.005, 9.83e-01) | 73038 | 0.11 |
| rs11063069 | *CCND2* | G | A | -0.0025 (0.004, 5.65e-01) | 0.0037 (0.005, 4.08e-01) | 75147 | 0.22 |
| rs1111875 | *HHEX/IDE* | C | T | -0.0066 (0.003, 2.88e-02) | -0.003 (0.003, 3.51e-01) | 75236 | 0.43 |
| rs11123406 | *BCL2L11* | T | C | 2e-04 (0.003, 9.60e-01) | 7e-04 (0.003, 8.40e-01) | 75235 | 0.37 |
| rs1116357 | *CCDC85A* | G | A | 0.0029 (0.003, 3.43e-01) | 0.0045 (0.003, 1.60e-01) | 75228 | 0.49 |
| rs11257655 | *CDC123/CAMK1D* | T | C | -0.0067 (0.004, 7.83e-02) | 0.0029 (0.004, 4.67e-01) | 68554 | 0.22 |
| rs11603334 | *ARAP1* | G | A | -0.0246 (0.004, 8.24e-10) | -0.0074 (0.004, 8.08e-02) | 75206 | 0.16 |
| rs11634397 | *ZFAND6* | G | A | -0.0015 (0.003, 6.47e-01) | -6e-04 (0.004, 8.59e-01) | 75237 | 0.36 |
| rs11708067 | *ADCY5* | A | G | -0.0239 (0.004, 2.50e-10) | -0.0072 (0.004, 7.12e-02) | 75227 | 0.21 |
| rs1182443 | *MNX1* | G | A | -0.012 (0.004, 2.85e-03) | -0.0019 (0.004, 6.55e-01) | 74533 | 0.21 |
| rs12242953 | *VPS26A* | G | A | -0.0016 (0.006, 8.03e-01) | -0.0045 (0.007, 5.03e-01) | 75224 | 0.07 |
| rs12427353 | *HNF1A(TCF1)* | G | A | -0.0048 (0.004, 2.14e-01) | -0.0053 (0.004, 1.90e-01) | 75196 | 0.21 |
| rs12454712 | *BCL2* | T | C | 0.0171 (0.006, 2.00e-03) | 0.0157 (0.006, 8.15e-03) | 47310 | 0.36 |
| rs12497268 | *PSMD6* | G | C | -0.005 (0.004, 2.08e-01) | 0.0056 (0.004, 1.84e-01) | 75220 | 0.19 |
| rs12571751 | *ZMIZ1* | A | G | 6e-04 (0.003, 8.38e-01) | 4e-04 (0.003, 8.95e-01) | 75228 | 0.47 |
| rs12681990 | *KCNU1* | C | T | 0.0052 (0.004, 2.28e-01) | 0.011 (0.005, 1.50e-02) | 74652 | 0.17 |
| rs12899811 | *PRC1* | G | A | -0.0016 (0.003, 6.32e-01) | 0.0043 (0.003, 2.18e-01) | 75232 | 0.31 |
| rs12970134 | *MC4R* | A | G | 0.0078 (0.003, 2.46e-02) | 0.0111 (0.004, 2.41e-03) | 75198 | 0.25 |
| rs13233731 | *KLF14* | G | A | 0.0037 (0.003, 2.23e-01) | 0.0083 (0.003, 9.03e-03) | 75215 | 0.48 |
| rs13389219 | *GRB14* | A | C | 0.0109 (0.003, 4.41e-04) | 0.0135 (0.003, 3.67e-05) | 75225 | 0.41 |
| rs1359790 | *SPRY2* | G | A | -0.0135 (0.003, 7.09e-05) | -0.0095 (0.004, 8.14e-03) | 75098 | 0.27 |
| rs1421085 | *FTO* | C | T | 0.0145 (0.003, 3.56e-06) | 0.0174 (0.003, 1.38e-07) | 75230 | 0.42 |
| rs1496653 | *UBE2E2* | A | G | -0.0116 (0.004, 1.21e-03) | -0.0082 (0.004, 3.05e-02) | 75149 | 0.22 |
| rs163184 | *KCNQ1* | G | T | -0.0074 (0.003, 2.15e-02) | 0.0021 (0.003, 5.35e-01) | 75153 | 0.5 |
| rs16927668 | *PTPRD* | T | C | -0.0032 (0.004, 3.95e-01) | -0.004 (0.004, 3.13e-01) | 75216 | 0.21 |
| rs17106184 | *FAF1* | G | A | -0.0076 (0.005, 1.55e-01) | -5e-04 (0.006, 9.26e-01) | 75236 | 0.09 |
| rs17168486 | *DGKB* | T | C | -0.018 (0.004, 1.08e-05) | 0.0089 (0.004, 3.82e-02) | 71882 | 0.18 |
| rs1727313 | *MPHOSPH9* | C | T | 0.0026 (0.004, 4.84e-01) | 0.0037 (0.004, 3.36e-01) | 75227 | 0.21 |

| | | | | | | | |
|---|---|---|---|---|---|---|---|
| rs17301514 | *ST64GAL1* | A | G | -0.0065 (0.006, 3.01e-01) | -0.0014 (0.007, 8.37e-01) | 56174 | 0.12 |
| rs17791513 | *TLE4(CHCHD9)* | A | G | -0.0042 (0.006, 4.78e-01) | -0.0016 (0.006, 7.94e-01) | 64543 | 0.07 |
| rs17867832 | *GCC1* | T | G | -0.0064 (0.005, 2.41e-01) | -0.0084 (0.006, 1.44e-01) | 68915 | 0.09 |
| rs1799884 | *GCK* | T | C | -0.0236 (0.004, 2.32e-08) | 0.0208 (0.004, 3.30e-06) | 68074 | 0.16 |
| rs1801282 | *PPARG* | C | G | 0.0158 (0.005, 5.23e-04) | 0.0186 (0.005, 1.09e-04) | 75233 | 0.12 |
| rs1861612 | *DNER* | T | C | 2e-04 (0.003, 9.60e-01) | 0.0044 (0.003, 1.69e-01) | 75238 | 0.45 |
| rs2007084 | *AP3S2* | G | A | -0.0021 (0.007, 7.57e-01) | -4e-04 (0.007, 9.52e-01) | 60850 | 0.09 |
| rs2023681 | *MTMR3/HORMAD2* | G | A | -0.001 (0.005, 8.54e-01) | 0.007 (0.006, 2.22e-01) | 75239 | 0.09 |
| rs2050188 | *HLA-DRB5* | T | C | 0.0065 (0.003, 5.07e-02) | 0.0074 (0.003, 3.39e-02) | 75159 | 0.36 |
| rs2244020 | *HLA-B* | G | A | -0.0029 (0.003, 3.70e-01) | 0.0019 (0.003, 5.80e-01) | 72272 | 0.38 |
| rs2261181 | *HMGA2* | T | C | -0.0041 (0.005, 4.25e-01) | 0.0011 (0.005, 8.39e-01) | 75234 | 0.09 |
| rs2292626 | *PLEKHA1* | C | T | 0.0016 (0.003, 6.07e-01) | -0.0023 (0.003, 4.66e-01) | 75238 | 0.49 |
| rs2296172 | *MACF1* | G | A | 0.0019 (0.004, 6.04e-01) | 0.0075 (0.004, 5.16e-02) | 74891 | 0.21 |
| rs2334499 | *DUSP8* | T | C | -0.0039 (0.003, 2.05e-01) | -0.0032 (0.003, 3.36e-01) | 75192 | 0.41 |
| rs243088 | *BCL11A* | T | A | -9e-04 (0.003, 7.78e-01) | 0.0039 (0.003, 2.32e-01) | 74640 | 0.46 |
| rs2447090 | *SRR* | A | G | -0.0015 (0.003, 6.42e-01) | -0.0053 (0.003, 1.26e-01) | 64307 | 0.38 |
| rs2796441 | *TLE1* | G | A | 0.0036 (0.003, 2.75e-01) | 0.0029 (0.004, 4.12e-01) | 75210 | 0.39 |
| rs2867125 | *TMEM18* | C | T | 0.0063 (0.004, 1.12e-01) | 0.0127 (0.004, 2.53e-03) | 75232 | 0.17 |
| rs2925979 | *CMIP* | T | C | 1e-04 (0.003, 9.69e-01) | 0.0042 (0.004, 2.41e-01) | 75189 | 0.3 |
| rs2943640 | *IRS1* | C | A | 0.0127 (0.003, 4.74e-05) | 0.0153 (0.003, 3.78e-06) | 75218 | 0.37 |
| rs3132524 | *POU5F1/TCF19* | G | A | 0.0026 (0.003, 4.56e-01) | 0.0031 (0.004, 3.95e-01) | 75230 | 0.25 |
| rs329122 | *PHF15* | A | G | -0.0057 (0.003, 7.12e-02) | -0.005 (0.003, 1.31e-01) | 75233 | 0.42 |
| rs340874 | *PROX1* | C | T | -0.0123 (0.003, 7.52e-05) | -0.002 (0.003, 5.47e-01) | 75167 | 0.47 |
| rs3802177 | *SLC30A8* | G | A | -0.0212 (0.004, 2.55e-09) | 7e-04 (0.004, 8.46e-01) | 68044 | 0.31 |
| rs3812547 | *GPSM1* | A | G | -0.0152 (0.004, 7.06e-04) | -0.0036 (0.005, 4.45e-01) | 46507 | 0.36 |
| rs4299828 | *ZFAND3* | A | G | 6e-04 (0.004, 8.65e-01) | -6e-04 (0.004, 8.89e-01) | 75230 | 0.2 |
| rs4430796 | *HNF1B(TCF2)* | G | A | -0.0093 (0.004, 2.06e-02) | -0.009 (0.004, 3.49e-02) | 54542 | 0.47 |
| rs4458523 | *WFS1* | G | T | -0.0049 (0.003, 1.09e-01) | 1e-04 (0.003, 9.64e-01) | 75192 | 0.41 |
| rs4502156 | *C2CD4A* | T | C | -0.0086 (0.003, 5.88e-03) | 0.0054 (0.003, 1.01e-01) | 75192 | 0.43 |
| rs459193 | *ANKRD55* | G | A | 0.009 (0.003, 9.41e-03) | 0.0185 (0.004, 4.76e-07) | 75230 | 0.25 |
| rs4865796 | *ARL15* | A | G | 0.0082 (0.003, 1.16e-02) | 0.0112 (0.003, 1.14e-03) | 75232 | 0.32 |
| rs516946 | *ANK1* | C | T | -0.0115 (0.003, 9.63e-04) | -0.0074 (0.004, 4.28e-02) | 75161 | 0.25 |
| rs5215 | *KCNJ11* | C | T | 0.0012 (0.003, 7.04e-01) | -0.005 (0.003, 1.24e-01) | 75155 | 0.39 |
| rs576674 | *KL* | G | A | -0.0147 (0.005, 1.48e-03) | -0.0028 (0.005, 5.68e-01) | 68623 | 0.16 |
| rs6017317 | *HNF4A* | G | T | -0.011 (0.004, 5.01e-03) | -0.01 (0.004, 1.53e-02) | 75224 | 0.17 |
| rs6723108 | *TMEM163* | T | G | -0.0076 (0.003, 1.49e-02) | -0.0114 (0.003, 5.39e-04) | 75045 | 0.45 |
| rs6795735 | *ADAMTS9* | C | T | -0.0026 (0.003, 4.00e-01) | 0.0022 (0.003, 5.05e-01) | 75229 | 0.43 |
| rs6808574 | *LPP* | C | T | -0.0034 (0.003, 2.69e-01) | 0.0044 (0.003, 1.78e-01) | 75205 | 0.4 |
| rs6813195 | *TMEM154* | C | T | -0.0075 (0.004, 3.76e-02) | -0.0119 (0.004, 1.82e-03) | 68582 | 0.29 |
| rs6878122 | *ZBED3* | G | A | -0.0028 (0.004, 4.52e-01) | 0.0057 (0.004, 1.47e-01) | 67346 | 0.27 |
| rs7177055 | *HMG20A* | A | G | -0.0031 (0.003, 3.49e-01) | 0.0051 (0.004, 1.48e-01) | 75233 | 0.29 |
| rs7202877 | *BCAR1* | T | G | -0.0067 (0.005, 1.75e-01) | 0.0035 (0.005, 5.02e-01) | 75181 | 0.1 |
| rs7258890 | *GIPR* | C | T | -0.0027 (0.004, 4.71e-01) | 6e-04 (0.004, 8.81e-01) | 75223 | 0.29 |
| rs731839 | *PEPD* | G | A | 0.0118 (0.003, 2.79e-04) | 0.016 (0.003, 3.30e-06) | 75142 | 0.33 |
| rs7403531 | *RASGRP1* | T | C | -0.0053 (0.004, 1.52e-01) | 0.0019 (0.004, 6.21e-01) | 75215 | 0.23 |
| rs7569522 | *RBMS1* | A | G | 5e-04 (0.003, 8.77e-01) | 0.0053 (0.003, 1.17e-01) | 68569 | 0.45 |

| | | | | | | | |
|---|---|---|---|---|---|---|---|
| rs7651090 | *IGF2BP2* | G | A | -0.011 (0.003, 9.68e-04) | -0.0015 (0.004, 6.78e-01) | 68594 | 0.31 |
| rs7674212 | *CISD2* | G | T | -0.0047 (0.003, 1.29e-01) | -9e-04 (0.003, 7.76e-01) | 75187 | 0.42 |
| rs780094 | *GCKR* | C | T | 8e-04 (0.003, 7.86e-01) | 0.0238 (0.003, 1.47e-13) | 75208 | 0.39 |
| rs7845219 | *TP53INP1* | T | C | -0.0076 (0.003, 1.15e-02) | -4e-04 (0.003, 9.06e-01) | 75229 | 0.5 |
| rs7903146 | *TCF7L2* | T | C | -0.0303 (0.003, 7.41e-19) | -0.0134 (0.004, 1.93e-04) | 75212 | 0.3 |
| rs791595 | *MIR129-LEP* | A | G | -0.0025 (0.004, 5.34e-01) | 0.0033 (0.004, 4.47e-01) | 68626 | 0.18 |
| rs7955901 | *TSPAN8/LGR5* | C | T | -0.0055 (0.003, 7.04e-02) | -0.0012 (0.003, 7.02e-01) | 75232 | 0.45 |
| rs7985179 | *MIR17HG* | T | A | 0.0079 (0.004, 2.41e-02) | 0.0114 (0.004, 2.18e-03) | 75239 | 0.25 |
| rs8090011 | *LAMA1* | G | C | -0.0024 (0.003, 4.70e-01) | -6e-04 (0.003, 8.58e-01) | 67134 | 0.37 |
| rs825476 | *CCDC92* | T | C | 0.006 (0.003, 4.49e-02) | 0.0065 (0.003, 4.00e-02) | 75233 | 0.42 |
| rs849135 | *JAZF1* | G | A | -0.005 (0.003, 9.63e-02) | -0.0028 (0.003, 3.83e-01) | 75225 | 0.49 |
| rs9309245 | *ASB3* | G | C | 0.001 (0.003, 7.47e-01) | -4e-04 (0.003, 9.14e-01) | 75236 | 0.34 |
| rs9368222 | *CDKAL1* | A | C | -0.0156 (0.003, 2.98e-06) | -0.0064 (0.004, 7.00e-02) | 75205 | 0.28 |
| rs9502570 | *SSR1/RREB1* | A | G | 0.0116 (0.004, 1.20e-03) | 0.0023 (0.004, 5.45e-01) | 75168 | 0.26 |
| rs9648716 | *BRAF* | T | A | -0.0103 (0.005, 3.60e-02) | -0.0072 (0.005, 1.62e-01) | 75214 | 0.11 |
| rs9674498 | *GLP2R* | G | T | 0.0038 (0.003, 2.41e-01) | 0.0076 (0.003, 2.64e-02) | 75226 | 0.31 |
| rs9940149 | *ITFG3* | G | A | -0.0036 (0.004, 3.84e-01) | -2e-04 (0.004, 9.67e-01) | 68628 | 0.17 |

**Table S6.** Established FG/FI loci with their effect, standard errors (SE), significance (P), sample size (N) and minor allele frequency (MAF) in HOMA-B/-IR GWIS.

| rsID | LOCUS | EA | NEA | Associated phenotype | HOMA-B, effect (SE, P) | HOMA-IR, effect (SE, P) | N | MAF |
|---|---|---|---|---|---|---|---|---|
| rs10195252 | *COBLL1/GRB14* | T | C | FI | 0.0106 (0.003, 5.86e-04) | 0.0135 (0.003, 3.19e-05) | 75199 | 0.41 |
| rs10747083 | *P2RX2* | A | G | FG | -0.0062 (0.004, 7.86e-02) | 0.0046 (0.004, 2.22e-01) | 74820 | 0.33 |
| rs10758593 | *GLIS3* | A | G | FG | -0.0183 (0.003, 1.90e-09) | -0.0072 (0.003, 2.55e-02) | 75203 | 0.42 |
| rs10811661 | *CDKN2A/B* | T | C | FG | -0.0125 (0.004, 2.11e-03) | 0.0034 (0.004, 4.39e-01) | 68591 | 0.18 |
| rs10830963 | *MTNR1B* | G | C | FG | -0.039 (0.004, 9.34e-25) | 0.0184 (0.004, 4.40e-06) | 68612 | 0.29 |
| rs10885122 | *ADRA2A* | G | T | FG | -0.0245 (0.005, 4.10e-07) | -0.0034 (0.005, 5.10e-01) | 75220 | 0.11 |
| rs11071657 | *FAM148B/VPS13C/ C2CD4A/B* | A | G | FG | -0.0067 (0.003, 4.41e-02) | 0.0039 (0.004, 2.63e-01) | 68605 | 0.37 |
| rs11603334 | *ARAP1* | G | A | FG | -0.0246 (0.004, 8.24e-10) | -0.0074 (0.004, 8.08e-02) | 75206 | 0.16 |
| rs11605924 | *CRY2* | A | C | FG | -0.0135 (0.003, 7.21e-06) | 0.0037 (0.003, 2.42e-01) | 75188 | 0.48 |
| rs11619319 | *PDX1* | G | A | FG | -0.0165 (0.004, 6.13e-06) | -0.002 (0.004, 6.07e-01) | 75224 | 0.22 |
| rs1167800 | *HIP1* | A | G | FI | 0.0148 (0.003, 1.79e-05) | 0.0162 (0.004, 8.01e-06) | 62225 | 0.45 |
| rs11708067 | *ADCY5* | A | G | FG | -0.0239 (0.004, 2.50e-10) | -0.0072 (0.004, 7.12e-02) | 75227 | 0.21 |
| rs11715915 | *AMT* | C | T | FG | 0.0012 (0.003, 7.28e-01) | 0.0109 (0.004, 1.92e-03) | 73791 | 0.31 |
| rs11920090 | *SLC2A2* | T | A | FG | -0.0174 (0.004, 8.07e-05) | 0.0017 (0.005, 7.10e-01) | 75238 | 0.13 |
| rs13179048 | *PCSK1* | C | A | FG | -0.0088 (0.003, 6.77e-03) | 0.0067 (0.003, 5.21e-02) | 75214 | 0.31 |
| rs1371614 | *DPYSL5* | T | C | FG | -0.0064 (0.004, 7.06e-02) | 0.0047 (0.004, 2.12e-01) | 75135 | 0.25 |
| rs1421085 | *FTO* | C | T | FI | 0.0145 (0.003, 3.56e-06) | 0.0174 (0.003, 1.38e-07) | 75230 | 0.42 |
| rs1483121 | *OR4S1* | G | A | FG | -0.0097 (0.005, 4.10e-02) | 0.0061 (0.005, 2.31e-01) | 62728 | 0.14 |
| rs1530559 | *YSK4* | A | G | FI | 0.01 (0.003, 4.24e-03) | 0.0143 (0.004, 9.81e-05) | 75236 | 0.46 |
| rs17168486 | *DGKB* | T | C | FG | -0.018 (0.004, 1.08e-05) | 0.0089 (0.004, 3.82e-02) | 71882 | 0.18 |
| rs174550 | *FADS1* | T | C | FG | -0.0197 (0.003, 4.67e-10) | -0.0051 (0.003, 1.26e-01) | 75237 | 0.34 |
| rs1799884 | *GCK* | T | C | FG | -0.0236 (0.004, 2.32e-08) | 0.0208 (0.004, 3.30e-06) | 68074 | 0.16 |
| rs2657879 | *GLS2* | G | A | FG | -0.0029 (0.004, 4.81e-01) | 0.0056 (0.004, 2.05e-01) | 68622 | 0.18 |
| rs2745353 | *RSPO3* | T | C | FI | 0.0154 (0.003, 4.53e-07) | 0.0136 (0.003, 2.41e-05) | 75223 | 0.49 |
| rs2785980 | *LYPLAL1* | T | C | FI | 0.012 (0.003, 1.82e-04) | 0.0162 (0.003, 1.77e-06) | 75231 | 0.33 |
| rs2943640 | *IRS1* | C | A | FI | 0.0127 (0.003, 4.74e-05) | 0.0153 (0.003, 3.78e-06) | 75218 | 0.37 |
| rs340874 | *PROX1* | C | T | FG | -0.0123 (0.003, 7.52e-05) | -0.002 (0.003, 5.47e-01) | 75167 | 0.47 |
| rs35767 | *IGF1* | G | A | FI | 0.0187 (0.004, 1.09e-05) | 0.0226 (0.005, 5.57e-07) | 68021 | 0.16 |
| rs3783347 | *WARS* | G | T | FG | -0.0073 (0.004, 6.24e-02) | 0.0045 (0.004, 2.77e-01) | 75170 | 0.21 |
| rs3802177 | *SLC30A8* | G | A | FG | -0.0212 (0.004, 2.55e-09) | 7e-04 (0.004, 8.46e-01) | 68044 | 0.31 |
| rs3829109 | *DNLZ* | G | A | FG | -0.0108 (0.005, 2.06e-02) | 4e-04 (0.005, 9.37e-01) | 59012 | 0.29 |
| rs4646949 | *C6orf107/UHRF1BP1* | T | G | FI | 0.0122 (0.003, 4.77e-04) | 0.0161 (0.004, 1.14e-05) | 75217 | 0.26 |
| rs4691380 | *PDGFC* | C | T | FI | 0.0109 (0.003, 7.62e-04) | 0.0139 (0.003, 4.63e-05) | 75234 | 0.33 |
| rs4841132 | *PPP1R3B* | A | G | FG/FI | 0.0165 (0.005, 1.37e-03) | 0.0342 (0.005, 3.22e-10) | 75219 | 0.09 |
| rs4865796 | *ARL15* | A | G | FI | 0.0082 (0.003, 1.16e-02) | 0.0112 (0.003, 1.14e-03) | 75232 | 0.32 |
| rs560887 | *G6PC2* | C | T | FG | -0.0465 (0.003, 4.74e-46) | 0.0047 (0.003, 1.74e-01) | 75173 | 0.3 |
| rs576674 | *KL* | G | A | FG | -0.0147 (0.005, 1.48e-03) | -0.0028 (0.005, 5.68e-01) | 68623 | 0.16 |
| rs6048205 | *FOXA2* | A | G | FG | -0.0398 (0.007, 3.12e-08) | -0.0077 (0.008, 3.10e-01) | 70262 | 0.06 |

| | | | | | | | | |
|---|---|---|---|---|---|---|---|---|
| rs6072275 | *TOP1* | A | G | FG | -0.0071 (0.004, 8.19e-02) | 0.0078 (0.004, 6.88e-02) | 75235 | 0.16 |
| rs6450057 | *PELO* | C | T | FI | 0.0089 (0.003, 3.61e-03) | 0.0089 (0.003, 5.90e-03) | 75204 | 0.44 |
| rs6943153 | *GRB10* | T | C | FG | 0.0043 (0.003, 1.80e-01) | 0.0137 (0.003, 4.93e-05) | 75221 | 0.32 |
| rs7258890 | *GIPR* | C | T | FG | -0.0027 (0.004, 4.71e-01) | 6e-04 (0.004, 8.81e-01) | 75223 | 0.29 |
| rs731839 | *PEPD* | G | A | FI | 0.0118 (0.003, 2.79e-04) | 0.016 (0.003, 3.30e-06) | 75142 | 0.33 |
| rs7651090 | *IGF2BP2* | G | A | FG | -0.011 (0.003, 9.68e-04) | -0.0015 (0.004, 6.78e-01) | 68594 | 0.31 |
| rs780094 | *GCKR* | C | T | FG/FI | 8e-04 (0.003, 7.86e-01) | 0.0238 (0.003, 1.47e-13) | 75208 | 0.39 |
| rs7903146 | *TCF7L2* | T | C | FG | -0.0303 (0.003, 7.41e-19) | -0.0134 (0.004, 1.93e-04) | 75212 | 0.3 |
| rs7944584 | *MADD* | A | T | FG | -0.0148 (0.003, 2.20e-05) | 0.0032 (0.004, 3.82e-01) | 75201 | 0.28 |
| rs9368222 | *CDKAL1* | A | C | FG | -0.0156 (0.003, 2.98e-06) | -0.0064 (0.004, 7.00e-02) | 75205 | 0.28 |
| rs9502570 | *SSR1/RREB1* | A | G | FG | 0.0116 (0.004, 1.20e-03) | 0.0023 (0.004, 5.45e-01) | 75168 | 0.26 |
| rs9884482 | *TET2* | C | T | FI | 0.013 (0.003, 4.61e-05) | 0.0141 (0.003, 2.84e-05) | 71820 | 0.39 |

**Table S7.** Established HbA1c loci with their effect, standard errors (SE), significance (P), sample size (N) and minor allele frequency (MAF) in HOMA-B/-IR GWIS.

| rsID | LOCUS | EA | NEA | Classification | HOMA-B, effect (SE, P) | HOMA-IR, effect (SE, P) | N | MAF |
|---|---|---|---|---|---|---|---|---|
| rs1046896 | *FN3KRP* | T | C | Unclassified | 9e-04 (0.003, 7.83e-01) | 0.0039 (0.003, 2.64e-01) | 75225 | 0.32 |
| rs10774625 | *ATXN2* | G | A | Erythrocytic | -0.0038 (0.003, 2.17e-01) | -0.0023 (0.003, 4.93e-01) | 75185 | 0.48 |
| rs10823343 | *HK1* | A | G | Unclassified | -0.0067 (0.004, 8.17e-02) | -0.0049 (0.004, 2.27e-01) | 68494 | 0.26 |
| rs10830963 | *MTNR1B* | G | C | Glycemic | -0.039 (0.004, 9.34e-25) | 0.0184 (0.004, 4.40e-06) | 68612 | 0.29 |
| rs11086054 | *MYO9B* | A | T | Unclassified | 0.003 (0.003, 3.81e-01) | 0.0051 (0.004, 1.64e-01) | 68574 | 0.31 |
| rs11224302 | *CNTN5* | C | T | Erythrocytic | 0.001 (0.005, 8.42e-01) | -0.0053 (0.005, 3.28e-01) | 75221 | 0.11 |
| rs11248914 | *ITFG3* | T | C | Erythrocytic | -7e-04 (0.003, 8.20e-01) | 0.0013 (0.003, 6.94e-01) | 75218 | 0.34 |
| rs11558471 | *SLC30A8* | A | G | Glycemic | -0.0197 (0.003, 3.12e-09) | 0.001 (0.004, 7.84e-01) | 74470 | 0.31 |
| rs11603334 | *ARAP1* | G | A | Glycemic | -0.0246 (0.004, 8.24e-10) | -0.0074 (0.004, 8.08e-02) | 75206 | 0.16 |
| rs11619319 | *PDX1* | G | A | Glycemic | -0.0165 (0.004, 6.13e-06) | -0.002 (0.004, 6.07e-01) | 75224 | 0.22 |
| rs11708067 | *ADCY5* | A | G | Glycemic | -0.0239 (0.004, 2.50e-10) | -0.0072 (0.004, 7.12e-02) | 75227 | 0.21 |
| rs11964178 | *C6orf183* | A | G | Erythrocytic | 0.005 (0.003, 1.02e-01) | 0.0055 (0.003, 8.71e-02) | 75210 | 0.43 |
| rs12132919 | *TMEM79* | A | C | Erythrocytic | 0.0021 (0.003, 5.42e-01) | -6e-04 (0.004, 8.73e-01) | 75194 | 0.29 |
| rs12621844 | *FOXN2* | T | C | Unclassified | 0.0036 (0.003, 2.45e-01) | 0.0076 (0.003, 2.16e-02) | 75228 | 0.39 |
| rs13134327 | *FREM3* | A | G | Glycemic | -0.0079 (0.003, 1.54e-02) | -0.0021 (0.003, 5.48e-01) | 75206 | 0.32 |
| rs13387347 | *G6PC2* | T | C | Glycemic | -0.012 (0.003, 1.35e-04) | -0.002 (0.003, 5.41e-01) | 74615 | 0.45 |
| rs1467311 | *KLF4* | G | A | Unclassified | 0.0022 (0.003, 4.90e-01) | -0.001 (0.003, 7.62e-01) | 75221 | 0.34 |
| rs1558902 | *FTO* | A | T | Unclassified | 0.0145 (0.003, 3.26e-06) | 0.0173 (0.003, 1.49e-07) | 75230 | 0.42 |
| rs17256082 | *SCRN3* | C | T | Unclassified | -0.0018 (0.003, 5.83e-01) | -0.0018 (0.003, 6.00e-01) | 75180 | 0.37 |
| rs174577 | *FADS2* | C | A | Glycemic | -0.019 (0.003, 2.58e-09) | -0.0046 (0.003, 1.71e-01) | 75226 | 0.34 |
| rs17509001 | *ATAD2B* | C | T | Unclassified | 0.0071 (0.005, 1.21e-01) | 0.0099 (0.005, 3.95e-02) | 75234 | 0.13 |
| rs17533903 | *MYO9B* | A | G | Erythrocytic | 0.0032 (0.004, 4.47e-01) | 0.003 (0.004, 4.98e-01) | 70516 | 0.22 |
| rs17747324 | *TCF7L2* | C | T | Glycemic | -0.0328 (0.004, 2.71e-17) | -0.0138 (0.004, 7.50e-04) | 75196 | 0.23 |
| rs1800562 | *HFE* | G | A | Erythrocytic | 1e-04 (0.007, 9.89e-01) | -0.0025 (0.007, 7.26e-01) | 68618 | 0.05 |
| rs198846 | *HFE* | G | A | Erythrocytic | 0.0024 (0.004, 5.91e-01) | 0.0033 (0.005, 4.85e-01) | 75197 | 0.14 |
| rs2073285 | *TMC6* | C | T | Unclassified | -0.0019 (0.005, 6.86e-01) | -0.0021 (0.005, 6.64e-01) | 46186 | 0.2 |
| rs2110073 | *PHB2* | T | C | Unclassified | 0.0123 (0.005, 2.06e-02) | 0.0126 (0.006, 2.48e-02) | 71133 | 0.09 |
| rs2191349 | *DGKB* | T | G | Glycemic | -0.0187 (0.003, 5.41e-10) | 0.002 (0.003, 5.39e-01) | 75224 | 0.47 |
| rs2237896 | *KCNQ1* | G | A | Glycemic | -0.0144 (0.008, 5.69e-02) | -0.0057 (0.008, 4.79e-01) | 68615 | 0.05 |
| rs2375278 | *SYF2* | A | G | Unclassified | -0.0092 (0.004, 3.65e-02) | -0.0076 (0.005, 9.83e-02) | 61698 | 0.17 |
| rs2383208 | *MTAP* | A | G | Glycemic | -0.0145 (0.004, 2.22e-04) | 0.002 (0.004, 6.28e-01) | 75164 | 0.18 |
| rs2408955 | *SENP1* | T | G | Erythrocytic | -0.0038 (0.003, 2.05e-01) | -0.0032 (0.003, 3.09e-01) | 74663 | 0.48 |
| rs267738 | *CERS2* | T | G | Unclassified | -0.0134 (0.004, 3.35e-04) | -0.0067 (0.004, 8.88e-02) | 73633 | 0.21 |
| rs282587 | *ATP11A* | G | A | Unclassified | -0.0036 (0.005, 4.40e-01) | 7e-04 (0.005, 8.92e-01) | 74667 | 0.13 |
| rs3782123 | *BET1L* | C | A | Unclassified | 7e-04 (0.004, 8.47e-01) | 0.0043 (0.004, 2.61e-01) | 70242 | 0.31 |
| rs3824065 | *GCK* | C | T | Glycemic | -0.0143 (0.003, 9.94e-06) | 0.0102 (0.003, 2.82e-03) | 75053 | 0.45 |
| rs4607517 | *GCK* | A | G | Glycemic | -0.0242 (0.004, 4.07e-09) | 0.0205 (0.004, 2.07e-06) | 74674 | 0.17 |
| rs4737009 | *ANK1* | A | G | Erythrocytic | -0.0087 (0.004, 2.08e-02) | -0.0039 (0.004, 3.25e-01) | 75215 | 0.24 |
| rs4745982 | *HK1* | T | G | Erythrocytic | 0.0103 (0.011, 3.57e-01) | 0.0026 (0.012, 8.24e-01) | 45217 | 0.08 |

| | | | | | | | | |
|---|---|---|---|---|---|---|---|---|
| rs4783565 | *CDH3* | A | G | Erythrocytic | -0.0067 (0.004, 5.96e-02) | -0.0057 (0.004, 1.33e-01) | 75158 | 0.28 |
| rs4820268 | *TMPRSS6* | G | A | Erythrocytic | 0.0042 (0.003, 1.89e-01) | 0.0069 (0.003, 4.25e-02) | 68613 | 0.44 |
| rs4894799 | *FNDC3B* | A | G | Unclassified | 0.0079 (0.003, 1.09e-02) | 0.0098 (0.003, 2.97e-03) | 75232 | 0.37 |
| rs560887 | *G6PC2* | C | T | Glycemic | -0.0465 (0.003, 4.74e-46) | 0.0047 (0.003, 1.74e-01) | 75173 | 0.3 |
| rs576674 | *KL* | G | A | Glycemic | -0.0147 (0.005, 1.48e-03) | -0.0028 (0.005, 5.68e-01) | 68623 | 0.16 |
| rs579459 | *ABO* | C | T | Glycemic | -0.0045 (0.004, 2.19e-01) | 0.0019 (0.004, 6.27e-01) | 75225 | 0.22 |
| rs592423 | *CITED2* | A | C | Erythrocytic | 0.0032 (0.003, 3.10e-01) | 0.0067 (0.003, 4.40e-02) | 68554 | 0.45 |
| rs6474359 | *ANK1* | T | C | Unclassified | 0.0036 (0.01, 7.06e-01) | -0.0024 (0.01, 8.09e-01) | 63710 | 0.03 |
| rs6980507 | *SLC20A2* | A | G | Erythrocytic | -8e-04 (0.003, 7.93e-01) | -0.0017 (0.003, 6.06e-01) | 68628 | 0.39 |
| rs7040409 | *C9orf47* | C | G | Erythrocytic | 0.0039 (0.007, 5.65e-01) | 6e-04 (0.007, 9.33e-01) | 68611 | 0.06 |
| rs7616006 | *SYN2* | A | G | Erythrocytic | 0.0013 (0.003, 6.76e-01) | 0.0016 (0.003, 6.29e-01) | 75207 | 0.43 |
| rs7756992 | *CDKAL1* | G | A | Glycemic | -0.0151 (0.003, 5.03e-06) | -0.0063 (0.003, 7.30e-02) | 75227 | 0.28 |
| rs8192675 | *SLC2A2* | T | C | Glycemic | -0.0134 (0.003, 5.51e-05) | -3e-04 (0.003, 9.28e-01) | 75239 | 0.3 |
| rs837763 | *CDT1* | T | C | Erythrocytic | -0.0051 (0.004, 1.58e-01) | -0.0037 (0.004, 3.33e-01) | 55770 | 0.44 |
| rs857691 | *SPTA1* | T | C | Erythrocytic | 0.0014 (0.003, 6.85e-01) | 0.002 (0.004, 5.71e-01) | 75226 | 0.26 |
| rs9604573 | *GAS6* | T | C | Unclassified | 0.0113 (0.004, 2.05e-03) | 0.0084 (0.004, 2.99e-02) | 68889 | 0.26 |
| rs9818758 | *USP4* | A | G | Unclassified | 0.0012 (0.004, 7.54e-01) | -0.0073 (0.004, 6.99e-02) | 75235 | 0.19 |
| rs9914988 | *ERAL1* | A | G | Erythrocytic | 0 (0.004, 9.93e-01) | 0.0034 (0.004, 3.93e-01) | 75213 | 0.19 |

**Table S8.** Genetic correlations ($r_g$) between *derived* HOMA-B and HOMA-IR and 13 glycaemic, 17 cardiometabolic traits, 5 inflammation markers, and T2D, including their standard errors (SEs), z-statistics and p-values (p indicates p-value, yielded by LD Score regression, $p_{fdr}$ indicates p-value, obtained after correction for multiple testing). Asterisk (*) indicates adjustment for BMI in primary GWAS meta-analysis. Table also reports SNP-heritability ($h^2$) and LD Score intercept (Intercept) for the phenotypes with their corresponding SEs.

| | | HOMA-B | | | | | HOMA-IR | | | | | SNP-$h^2$ | | | |
|---|---|---|---|---|---|---|---|---|---|---|---|---|---|---|---|
| | | Genetic Correlation | | | | | Genetic Correlation | | | | | | | | |
| | | $r_g$ | se | z | p | $p_{fdr}$ | $r_g$ | se | z | p | $p_{fdr}$ | $h^2$ | se | Intercept | se |
| T2D | T2D | 0.05 | 0.10 | 0.50 | 0.62 | 0.71 | 0.56 | 0.09 | 6.24 | $4.43\times10^{-10}$ | $2.31\times10^{-9}$ | 0.10 | 0.01 | 1.01 | 0.01 |
| Glycaemic Traits | Fasting Glucose (FG) | -0.32 | 0.14 | -2.34 | 0.02 | 0.05 | 0.49 | 0.08 | 6.33 | $2.45\times10^{-10}$ | $1.37\times10^{-9}$ | 0.09 | 0.02 | 1.02 | 0.01 |
| | Fasting Insulin (FI) | 0.80 | 0.04 | 18.12 | $2.40\times10^{-73}$ | $8.77\times10^{-72}$ | 0.98 | 0.005 | 202.87 | <0.001 | <0.001 | 0.06 | 0.01 | 1.04 | 0.01 |
| | $HbA1_C$ | -0.02 | 0.13 | -0.16 | 0.87 | 0.93 | 0.28 | 0.11 | 2.61 | $8.9\times10^{-3}$ | 0.024 | 0.06 | 0.01 | 1.00 | 0.01 |
| | Fasting Proinsulin (FPI) | 0.03 | 0.17 | 0.16 | 0.87 | 0.93 | 0.22 | 0.12 | 1.89 | 0.06 | 0.12 | 0.20 | 0.09 | 0.98 | 0.01 |
| | Insulin Sensitivity Index (ISI) | -0.90 | 0.19 | -4.67 | $2.97\times10^{-6}$ | $1.20\times10^{-5}$ | -1.04 | 0.18 | -5.83 | $5.56\times10^{-9}$ | $2.71\times10^{-8}$ | 0.10 | 0.03 | 0.99 | 0.01 |
| | Insulin Sensitivity Index (ISI)* | -0.60 | 0.16 | -3.82 | $1.00\times10^{-4}$ | $3.32\times10^{-4}$ | -0.63 | 0.13 | -4.76 | $1.91\times10^{-6}$ | $8.18\times10^{-6}$ | 0.12 | 0.03 | 0.98 | 0.01 |
| | Disposition Index (DI) | 0.26 | 0.24 | 1.07 | 0.28 | 0.45 | -0.26 | 0.25 | -1.04 | 0.30 | 0.45 | 0.12 | 0.09 | 1.00 | 0.01 |
| | Corrected Insulin Response (CIR) adjusted for Insulin Sensitivity Index (ISI) | 0.56 | 0.46 | 1.24 | 0.22 | 0.37 | 0.04 | 0.29 | 0.14 | 0.89 | 0.93 | 0.07 | 0.10 | 1.01 | 0.01 |
| | Corrected Insulin Response (CIR) | 0.53 | 0.26 | 2.04 | 0.04 | 0.09 | 0.15 | 0.22 | 0.68 | 0.50 | 0.61 | 0.13 | 0.09 | 1.00 | 0.01 |
| | Incr30 | 0.55 | 0.35 | 1.56 | 0.12 | 0.23 | 0.56 | 0.36 | 1.54 | 0.12 | 0.23 | 0.10 | 0.10 | 1.01 | 0.01 |
| | AUC for insulin | 0.98 | 1.32 | 0.74 | 0.46 | 0.60 | 0.83 | 1.13 | 0.73 | 0.47 | 0.60 | 0.05 | 0.11 | 1.01 | 0.01 |
| | Ins30* | 0.97 | 0.80 | 1.21 | 0.23 | 0.38 | 0.57 | 0.54 | 1.06 | 0.29 | 0.45 | 0.06 | 0.09 | 1.01 | 0.01 |
| | 2 hour Glucose* | 0.05 | 0.15 | 0.32 | 0.75 | 0.84 | 0.06 | 0.14 | 0.41 | 0.69 | 0.78 | 0.11 | 0.03 | 0.99 | 0.01 |

**Table S8.** Continued

| | | HOMA-B | | | | | HOMA-IR | | | | | SNP-$h^2$ | | | |
|---|---|---|---|---|---|---|---|---|---|---|---|---|---|---|---|
| | | Genetic Correlation | | | | | Genetic Correlation | | | | | | | | |
| | | $r_g$ | se | z | p | $p_{fdr}$ | $r_g$ | se | z | p | $p_{fdr}$ | $h^2$ | se | Intercept | se |
| Lipids | Triglycerides (TG) | 0.35 | 0.07 | 5.34 | $9.23\times10^{-8}$ | $4.21\times10^{-7}$ | 0.42 | 0.10 | 4.01 | $6.12\times10^{-5}$ | $2.13\times10^{-4}$ | 0.15 | 0.03 | 0.86 | 0.02 |
| | High Density Lipoprotein Cholesterol (HDL-C) | -0.36 | 0.08 | -4.23 | $2.34\times10^{-5}$ | $8.98\times10^{-5}$ | -0.53 | 0.08 | -6.76 | $1.39\times10^{-11}$ | $9.20\times10^{-11}$ | 0.11 | 0.02 | 0.93 | 0.04 |
| | Low Density Lipoprotein Cholesterol (LDL-C) | 0.15 | 0.08 | 1.74 | 0.08 | 0.17 | 0.09 | 0.08 | 1.11 | 0.27 | 0.43 | 0.09 | 0.02 | 0.96 | 0.04 |
| | Total Cholesterol (TC) | 0.09 | 0.07 | 1.27 | 0.20 | 0.37 | 0.005 | 0.07 | 0.07 | 0.94 | 0.95 | 0.12 | 0.02 | 0.93 | 0.03 |
| Obesity | Hip Circumference (HIP) | 0.39 | 0.06 | 6.55 | $5.82\times10^{-11}$ | $3.54\times10^{-10}$ | 0.53 | 0.06 | 9.16 | $5.06\times10^{-20}$ | $4.61\times10^{-19}$ | 0.15 | 0.01 | 0.75 | 0.01 |
| | Waist Circumference (WC) | 0.52 | 0.06 | 9.26 | $2.14\times10^{-20}$ | $2.23\times10^{-19}$ | 0.70 | 0.05 | 14.06 | $6.44\times10^{-45}$ | $1.57\times10^{-43}$ | 0.14 | 0.01 | 0.74 | 0.01 |
| | Waist to Hip Ratio (WHR) | 0.50 | 0.06 | 8.70 | $3.32\times10^{-18}$ | $2.69\times10^{-17}$ | 0.66 | 0.05 | 12.65 | $1.16\times10^{-36}$ | $2.11\times10^{-35}$ | 0.10 | 0.01 | 0.83 | 0.01 |
| | BMI | 0.43 | 0.05 | 8.63 | $6.15\times10^{-18}$ | $4.49\times10^{-17}$ | 0.62 | 0.05 | 12.43 | $1.73\times10^{-35}$ | $2.52\times10^{-34}$ | 0.14 | 0.01 | 0.65 | 0.01 |
| T2D complications | Systolic Blood Preasure (SBP) | -0.08 | 0.11 | -0.68 | 0.50 | 0.61 | 0.10 | 0.10 | 0.94 | 0.35 | 0.51 | 0.11 | 0.02 | 0.95 | 0.01 |
| | Diastolic Dlood Preasure (DBP) | -0.11 | 0.12 | -0.98 | 0.33 | 0.48 | 0.01 | 0.10 | 0.05 | 0.96 | 0.96 | 0.13 | 0.02 | 0.94 | 0.01 |
| | Hypertension (HTN) | -0.01 | 0.13 | -0.09 | 0.92 | 0.95 | 0.16 | 0.13 | 1.24 | 0.21 | 0.37 | 0.15 | 0.03 | 0.96 | 0.01 |
| | Coronary Artery Disease (CAD) | 0.09 | 0.10 | 0.86 | 0.39 | 0.55 | 0.19 | 0.10 | 1.89 | 0.06 | 0.12 | 0.07 | 0.01 | 1.03 | 0.01 |
| | eGFR creatinine Diabetes Mellitus | -0.09 | 0.18 | -0.52 | 0.60 | 0.71 | 0.03 | 0.18 | 0.19 | 0.85 | 0.93 | 0.09 | 0.04 | 1.00 | 0.01 |
| | eGFR creatinine non Diabetes Mellitus | -0.05 | 0.06 | -0.79 | 0.43 | 0.59 | -0.05 | 0.07 | -0.70 | 0.49 | 0.61 | 0.11 | 0.01 | 0.98 | 0.01 |
| | eGFR creatinine all | -0.05 | 0.06 | -0.78 | 0.43 | 0.59 | -0.04 | 0.07 | -0.53 | 0.59 | 0.71 | 0.11 | 0.01 | 0.98 | 0.01 |
| | eGFR cystatin C all | -0.31 | 0.13 | -2.49 | 0.01 | 0.03 | -0.27 | 0.11 | -2.34 | 0.02 | 0.05 | 0.16 | 0.07 | 0.95 | 0.01 |
| | Chronic Kidney Disease (CKD) | 0.31 | 0.14 | 2.23 | 0.03 | 0.06 | 0.23 | 0.14 | 1.59 | 0.11 | 0.22 | 0.02 | 0.01 | 1.02 | 0.01 |

**Table S8.** Continued

| | | HOMA-B | | | | | HOMA-IR | | | | | SNP-$h^2$ | | | |
|---|---|---|---|---|---|---|---|---|---|---|---|---|---|---|---|
| | | Genetic Correlation | | | | | Genetic Correlation | | | | | | | | |
| | | $r_g$ | se | z | p | $p_{fdr}$ | $r_g$ | se | z | p | $p_{fdr}$ | $h^2$ | se | Intercept | se |
| Inflammation | White Blood Cell counts (WBC) | 0.14 | 0.19 | 0.74 | 0.46 | 0.60 | 0.23 | 0.18 | 1.26 | 0.21 | 0.37 | 0.05 | 0.02 | 1.00 | 0.01 |
| | ICAM-1 | 0.14 | 0.17 | 0.82 | 0.41 | 0.58 | 0.03 | 0.17 | 0.16 | 0.88 | 0.93 | 0.74 | 0.50 | 1.04 | 0.03 |
| | C-Reactive Protein (CRP) | 0.28 | 0.09 | 3.25 | $1.20\times10^{-3}$ | $3.65\times10^{-3}$ | 0.33 | 0.11 | 3.16 | $1.60\times10^{-3}$ | $4.67\times10^{-3}$ | 0.13 | 0.03 | 1.00 | 0.02 |
| | PAI-1 | 0.78 | 0.23 | 3.36 | $8.00\times10^{-4}$ | $2.54\times10^{-3}$ | 0.92 | 0.23 | 4.01 | $6.13\times10^{-5}$ | $2.13\times10^{-4}$ | 0.06 | 0.03 | 1.00 | 0.01 |
| | ADIPONECTIN | -0.24 | 0.11 | -2.16 | 0.03 | 0.07 | -0.30 | 0.10 | -2.85 | $4.40\times10^{-3}$ | 0.012 | 0.11 | 0.02 | 0.96 | 0.01 |

**Table S9.** Genetic correlation ($r_g$) between inferred and published HOMA-B and HOMA-IR, the LD score regression intercepts, SNP-heritability and their standard errors (SE).

| | Published | Inferred Global Correction | Inferred Local Correction |
|---|---|---|---|
| | | HOMA-B | |
| $r_g$ (SE), p-value | - | 1.12 (0.07), $4.30\times10^{-66}$ | 1.14 (0.07), $2.22\times10^{-56}$ |
| Intercept (SE) | 0.99 (0.007) | 1.043 (0.008) | 1.05 (0.008) |
| $h^2$ (SE) | 0.08 (0.014) | 0.058 (0.01) | 0.055 (0.01) |
| | | HOMA-IR | |
| $r_g$ (SE), p-value | - | 1.18 (0.07), $3.58\times10^{-58}$ | 1.18 (0.07), $1.40\times10^{-58}$ |
| Intercept (SE) | 1.003 (0.007) | 1.03 (0.008) | 1.03 (0.008) |
| $h^2$ (SE) | 0.07 (0.01) | 0.059 (0.008) | 0.060 (0.008) |

**Figure S1.** Manhattan Plots of published meta-analysis results

HOMA-B

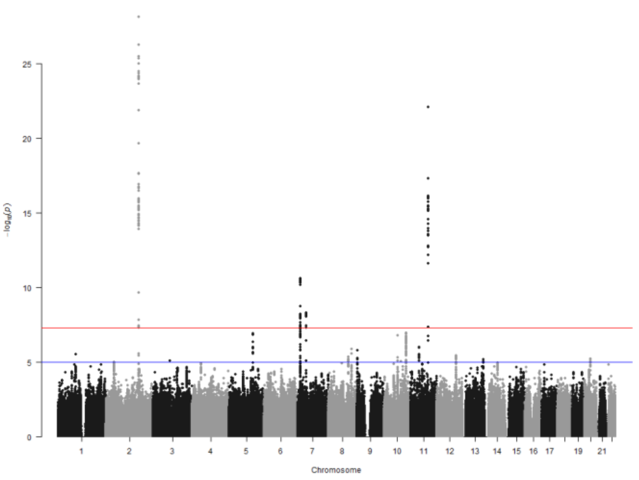


HOMA-IR

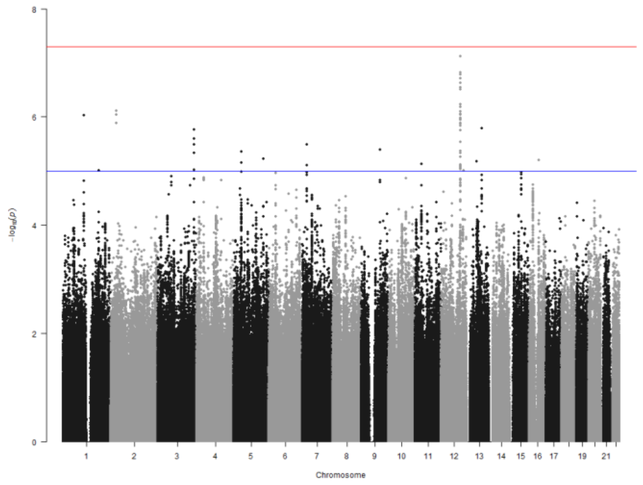


**Figure S2.** Manhattan Plots of inferred meta-analysis results

HOMA-B

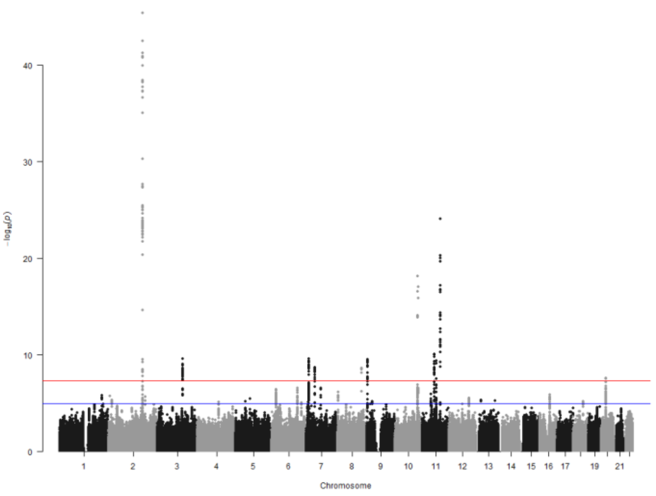


HOMA-IR

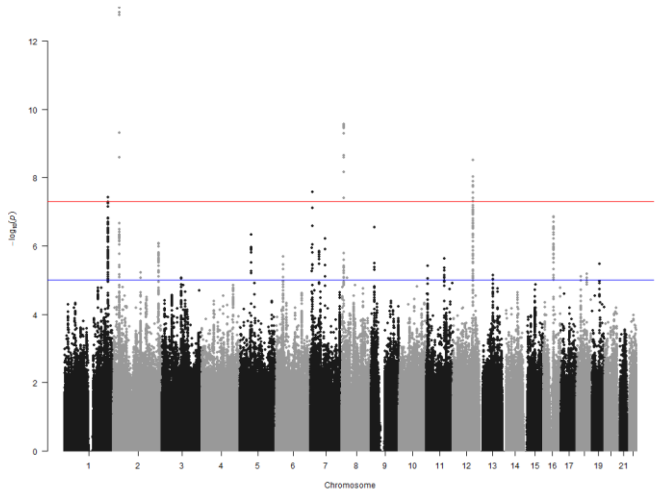

**Figure S3.** QQ plots of published meta-analysis results

HOMA-B

HOMA-IR

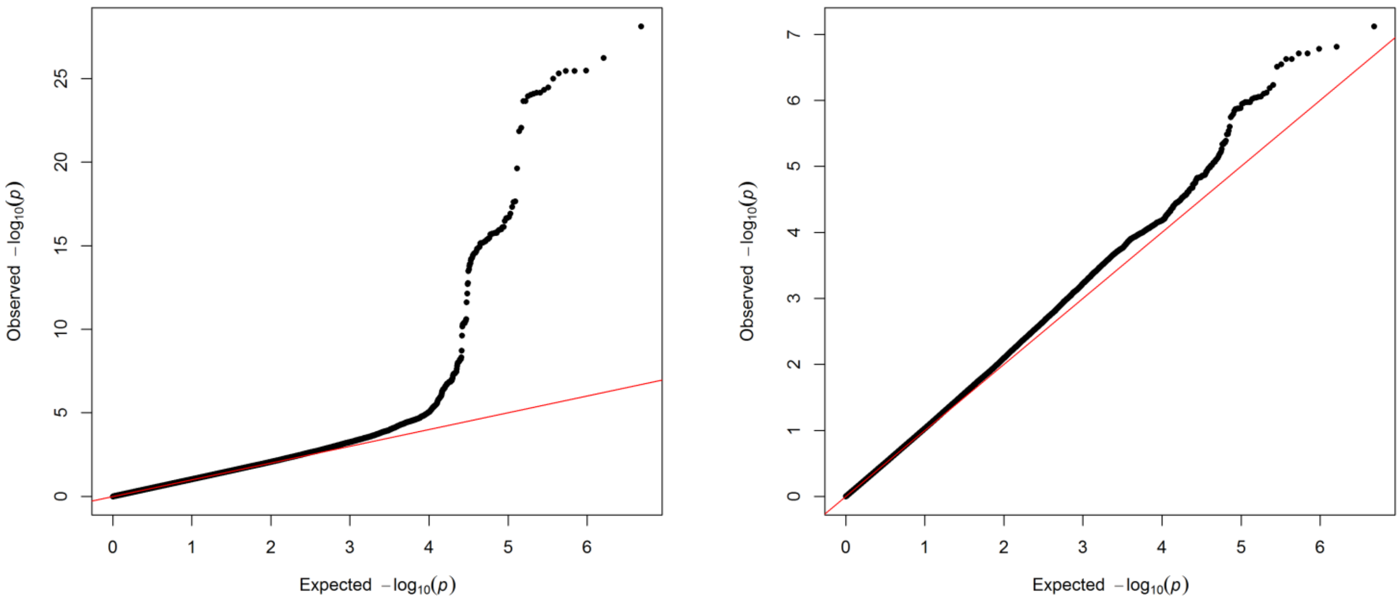


**Figure S4.** QQ plots of inferred meta-analysis results

HOMA-B

HOMA-IR

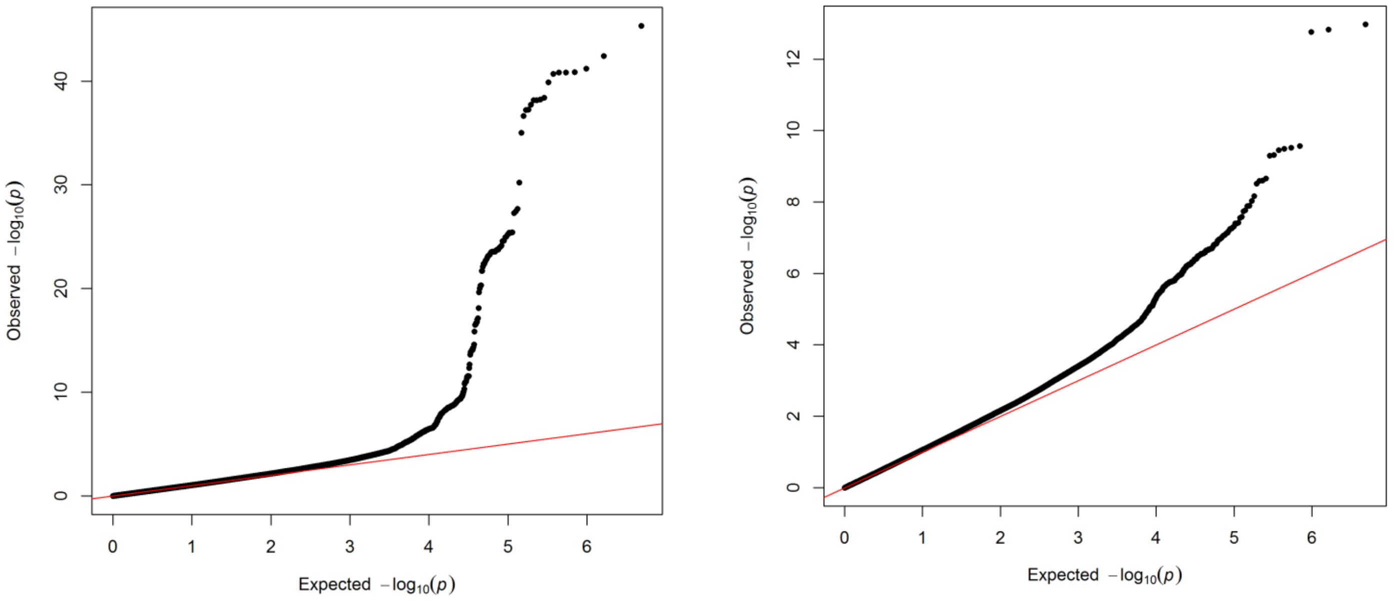

**Figure S5.** Manhattan and QQ plots, showing GWIS results for HOMA-B (a)/-IR (b). Novel loci are highlighted in purple/red colour. The X-axis denotes chromosome and base-pair positions of SNPs against –log10(P) plotted on the Y-axis. The red horizontal line indicates genome-wide significance level ($P<5\times10^{-8}$), and the blue line represents suggestive significance level ($P<1\times10^{-5}$). HOMA-B/IR GWAS significant loci are labelled by the name of nearest gene, established FG(a)/FI(b) loci are also represented in blue/green colour, respectively.

a) HOMA-B

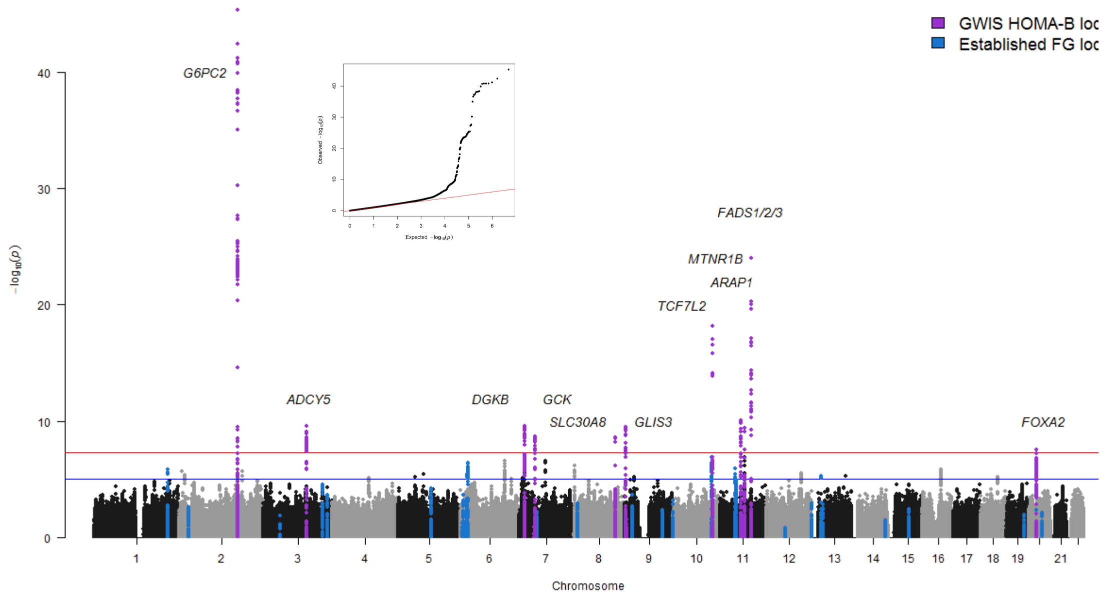

**Figure S5.** Continued

b) HOMA-IR

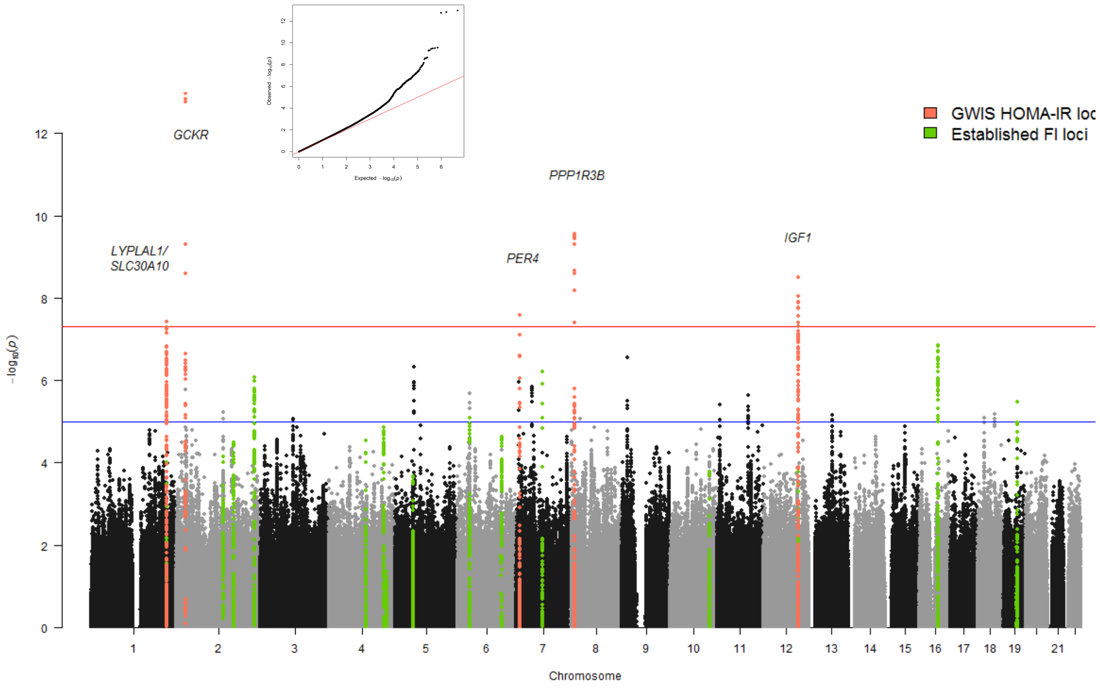

**Figure S6.** FG (a) and FI (b) average estimates from each participating cohort, plotted against –log10(P) for each HOMA-B/-IR locus lead SNP that was genome-wide significant in main analysis. The horizontal line indicates genome-wide significance level (P<5×10$^{-8}$). Vertical lines indicate the FG/FI means and dashed lines indicate 1*SDs* from either of FG/FI means.

**a** **b**

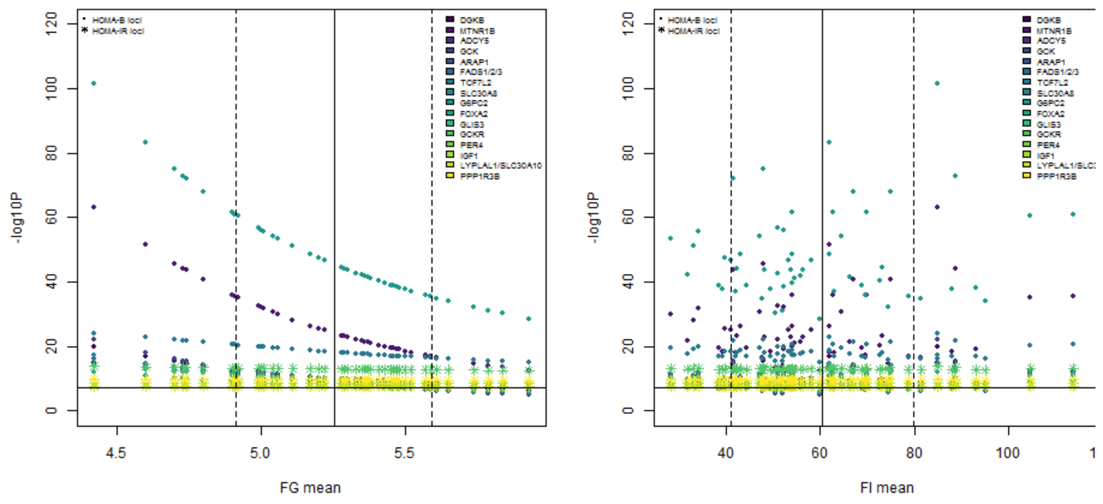

**Figure S7.**

HOMA-B HOMA-IR

a. Peaks

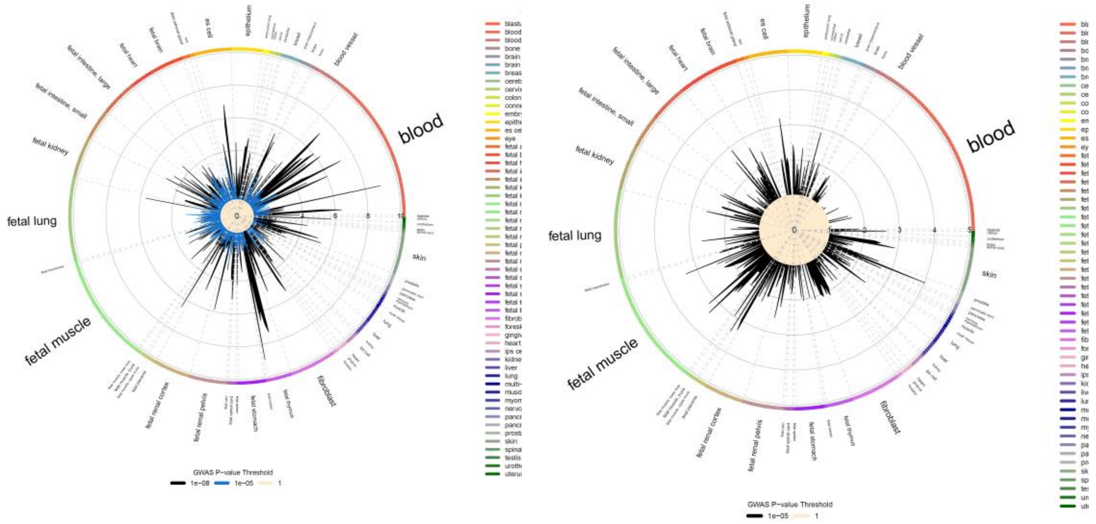


b. Footprints

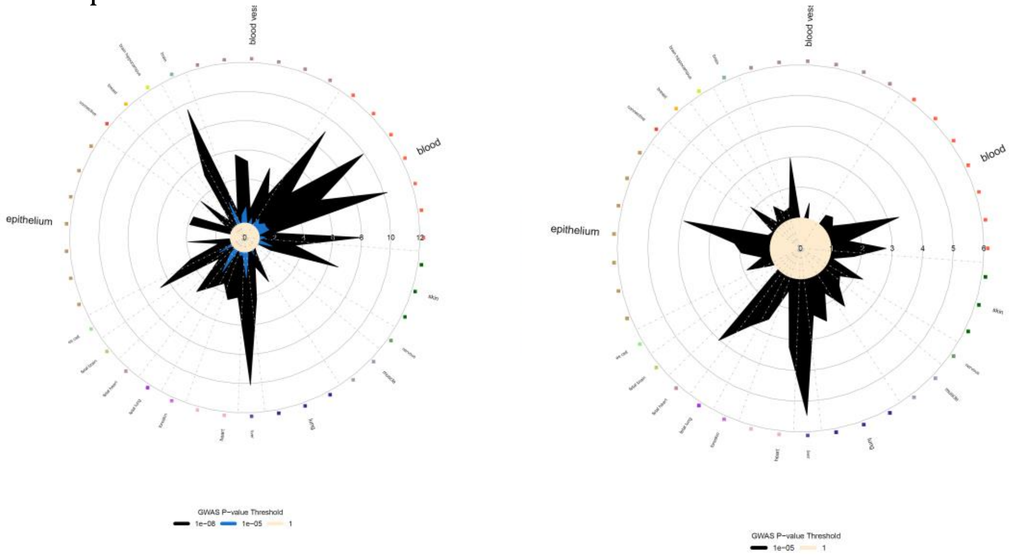


c. Chromatin states

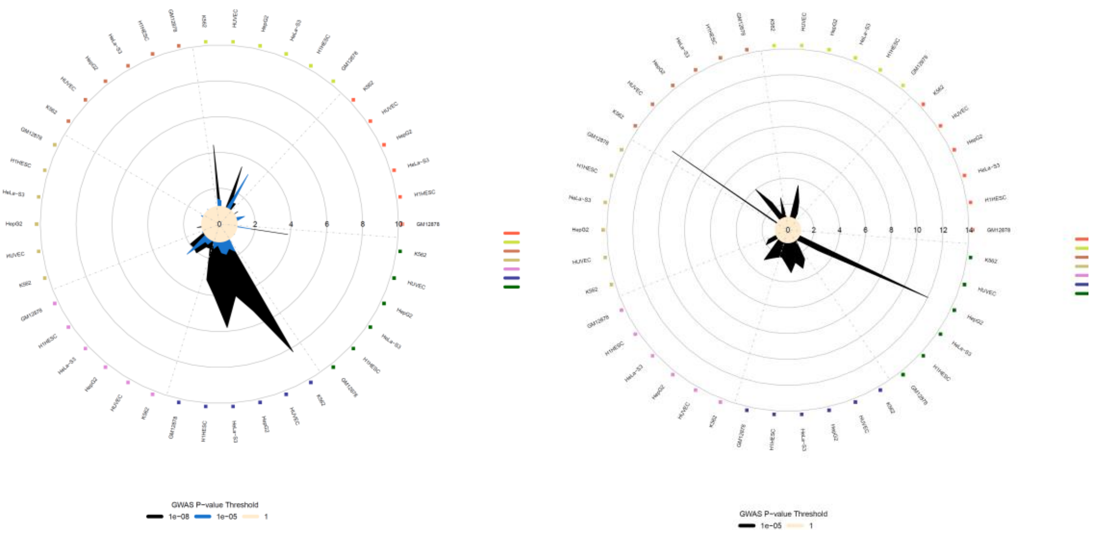

**Figure S7. Continued.**

HOMA-B HOMA-IR

d. Histone modifications

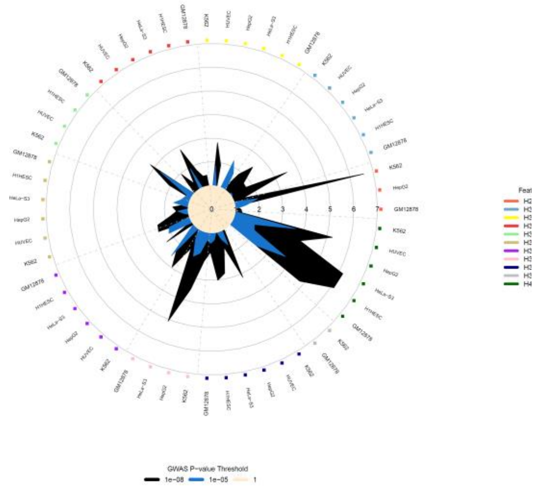


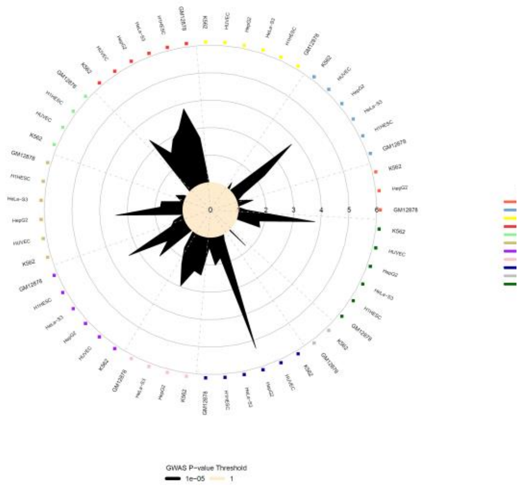


e. TFBS

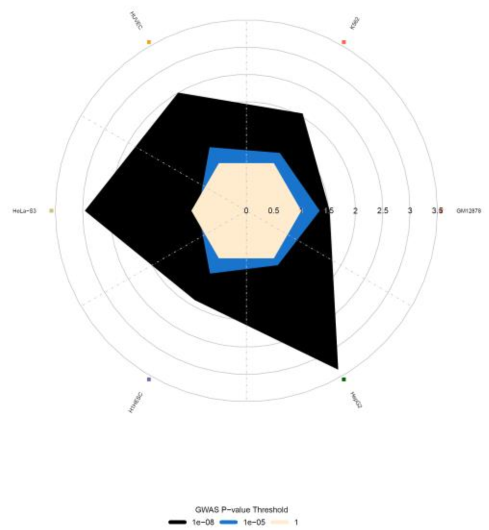


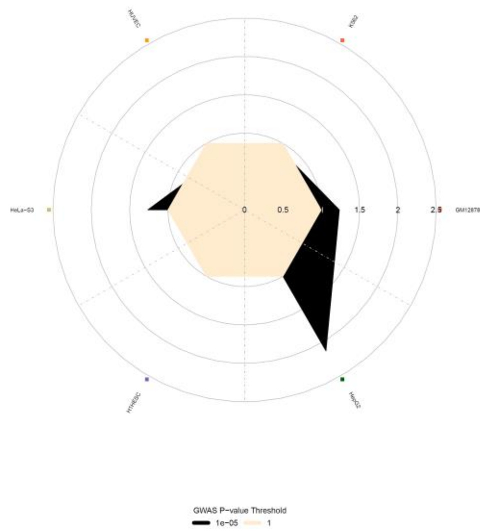


f. FAIRE

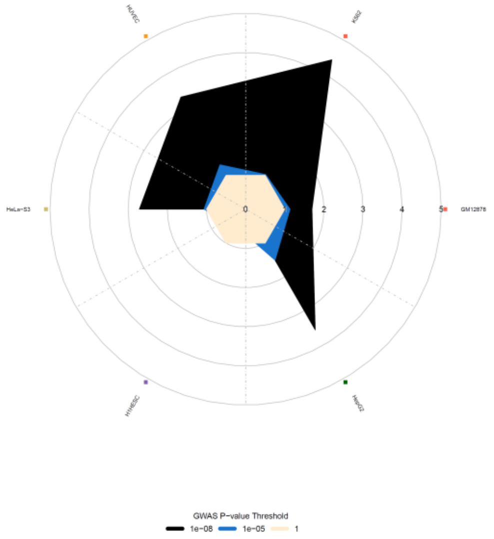


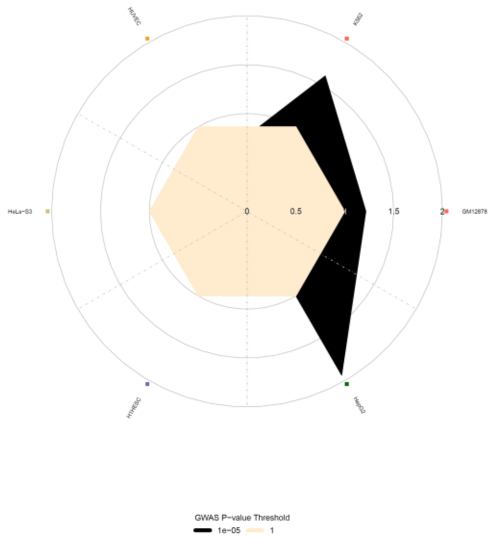

**Figure S8.** Effect size, –log10P and SE comparison between published and inferred meta-analysis results for 'global' (dark color) and 'local' (light color) correction.

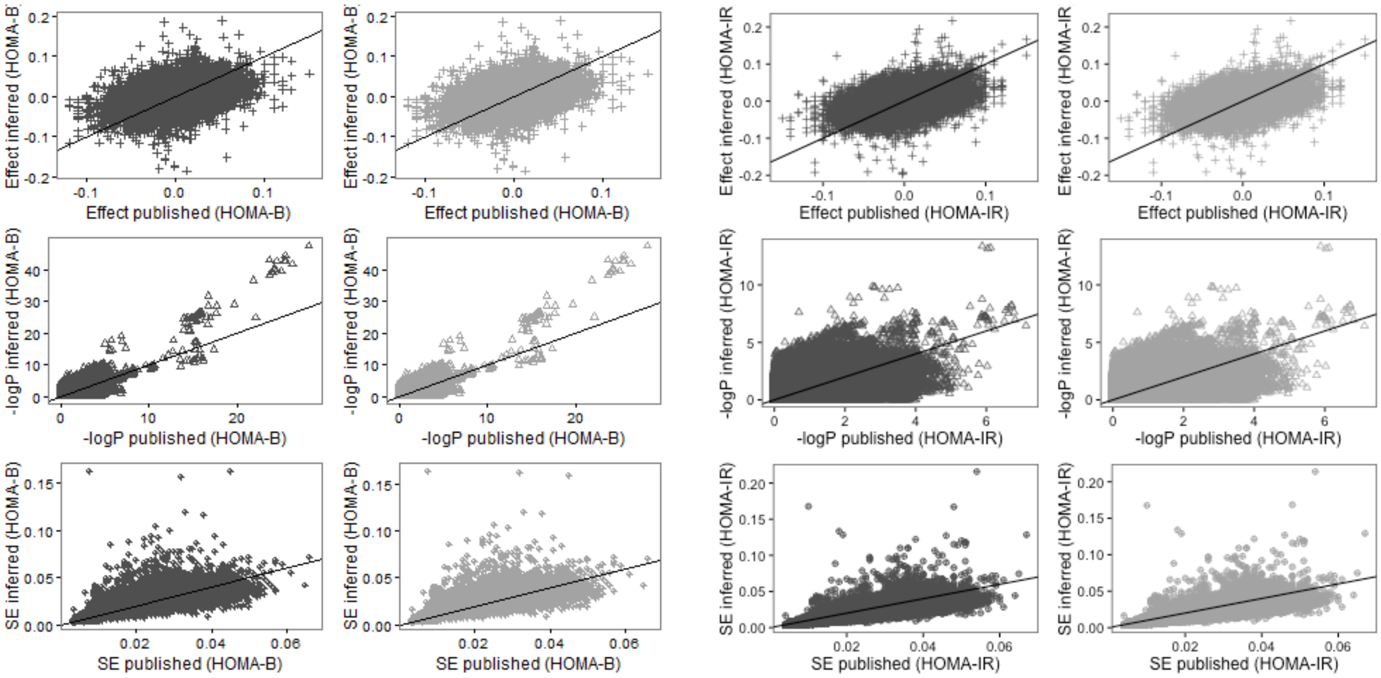


**Figure S9.** Difference in SE, effect size and Z-score plotted against N for 'global' (dark color) and 'local' (light color) correction.

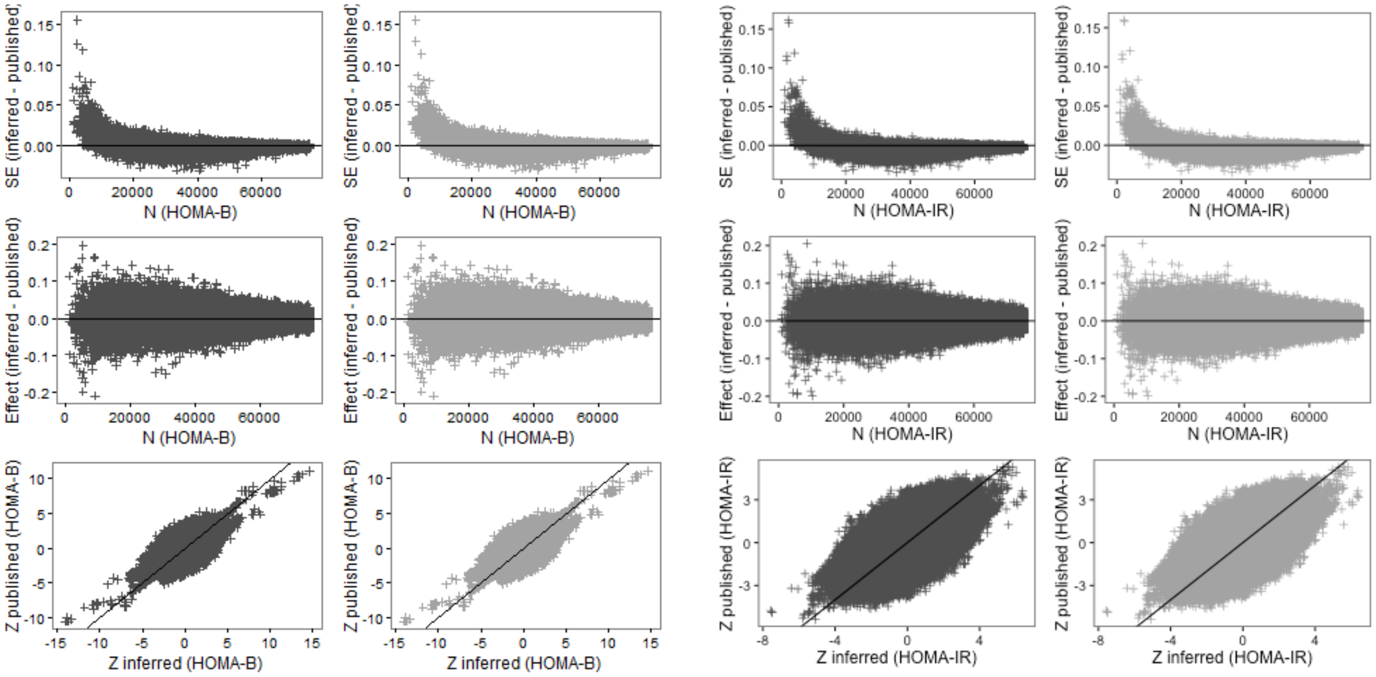

**Figure S10.** Distributions of the difference between inferred and published meta-analysis SE, for all SNPs (left) and for SNPs with smaller difference (right)

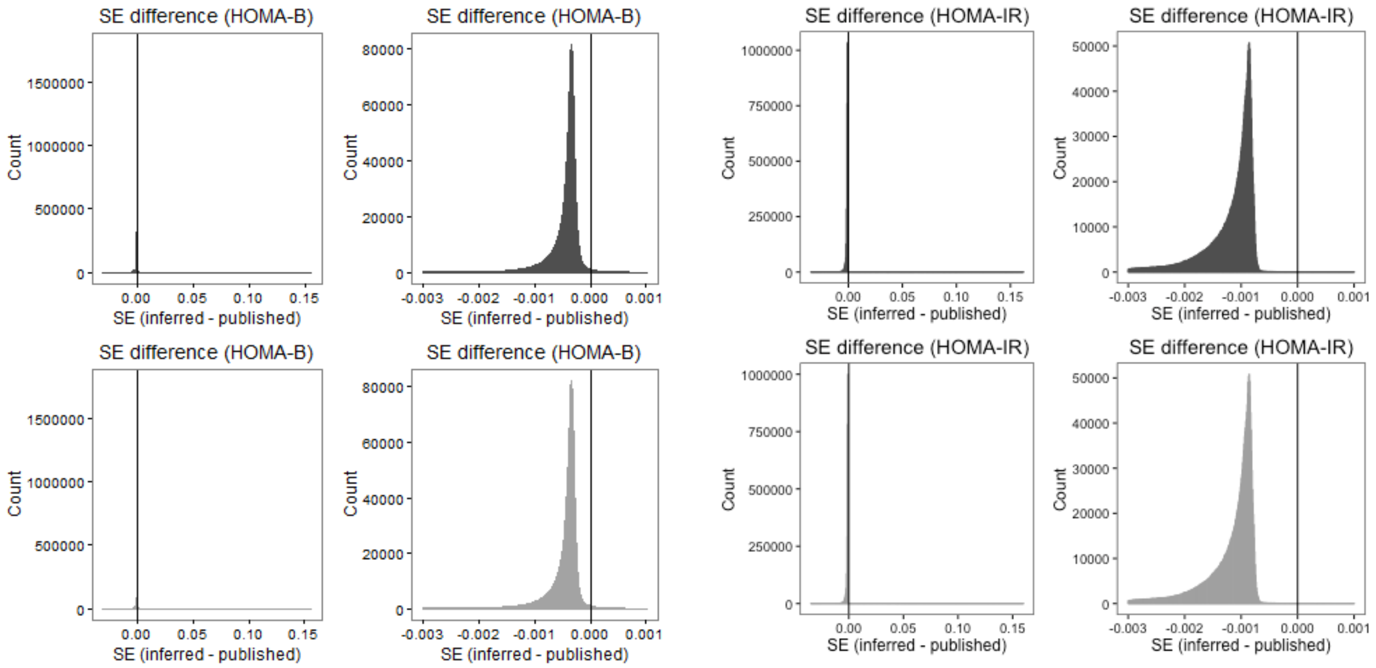


**Figure S11.** Distributions of the difference between inferred and published meta-analysis effect sizes, for all SNPs (left) and for SNPs with smaller difference (right)

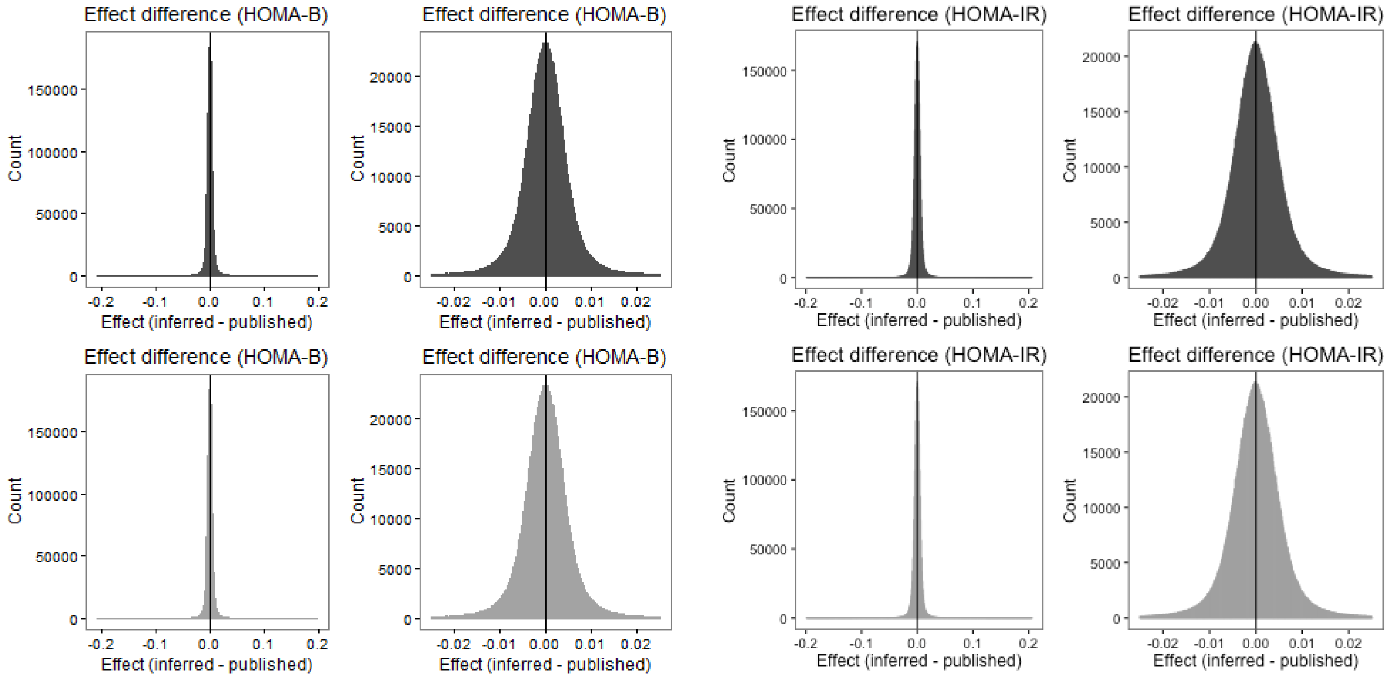

**Figure S12.** Distributions of the difference between inferred and published meta-analysis Z scores and difference in effect sizes plotted against difference in SE's for 'global' (dark color) and 'local' (light color) versions.

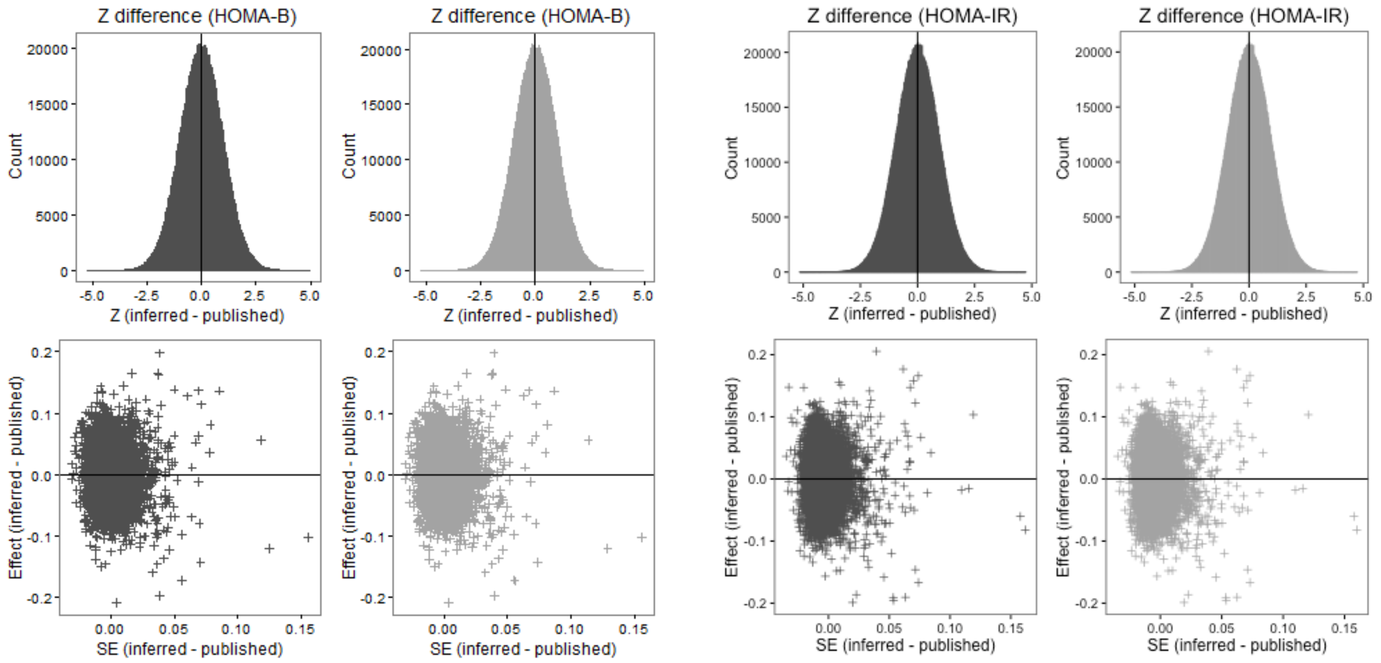


**Figure S13.** SE differences plotted against MAF in published and inferred meta-analysis for 'global' (dark color) and 'local' (light color) correction

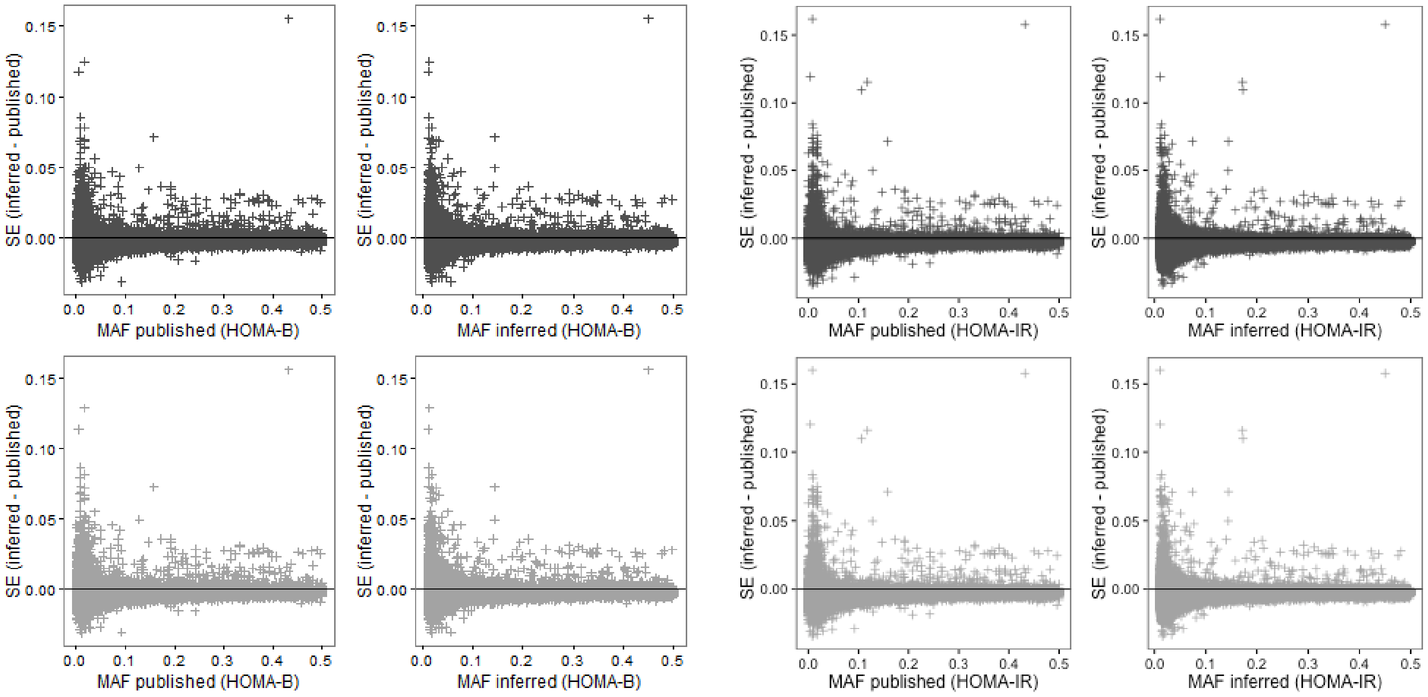

**Figure S14.** Difference in SE (left) and effect sizes (right) between inferred and published GWAS plotted against N. Dashed line indicated difference in SE < -0.003. Datasets are filtered with N > 35,000 and MAF > 0.01

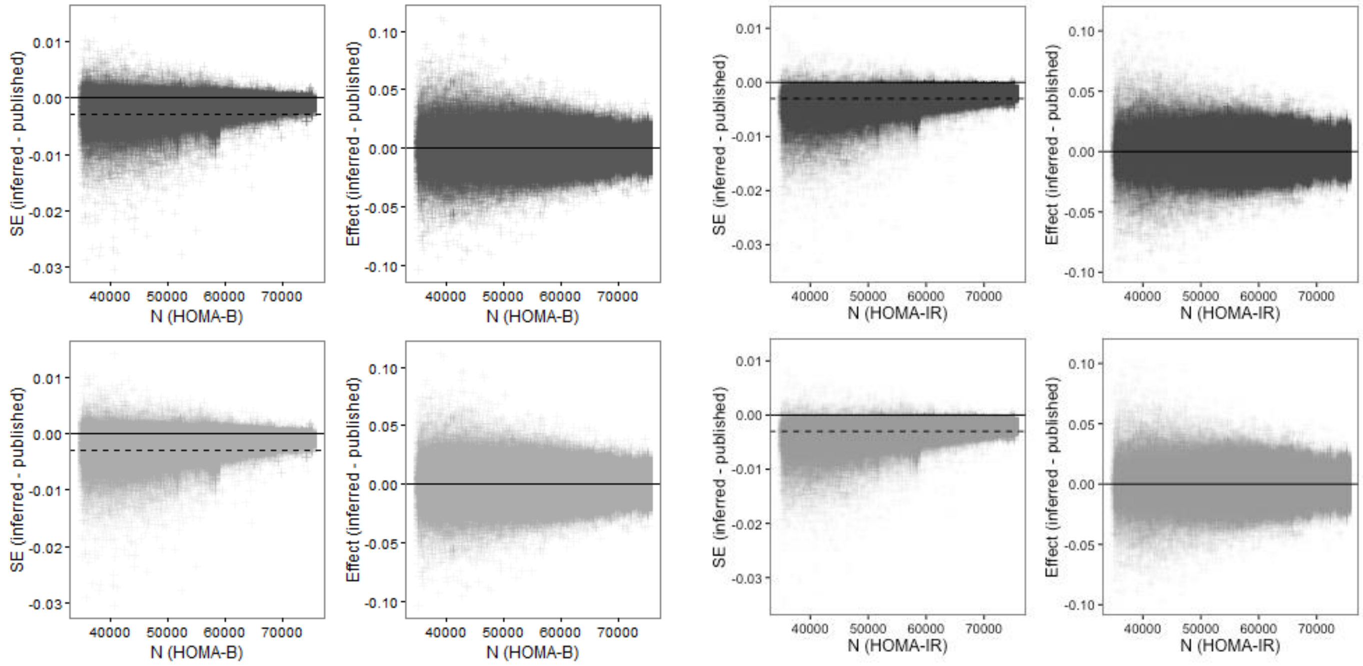


**Figure S15.** Difference in SE between inferred and published GWAS plotted against SE published (left) and inferred (right). Dashed line indicated difference in SE < -0.003. Datasets are filtered with N > 35,000 and MAF > 0.01

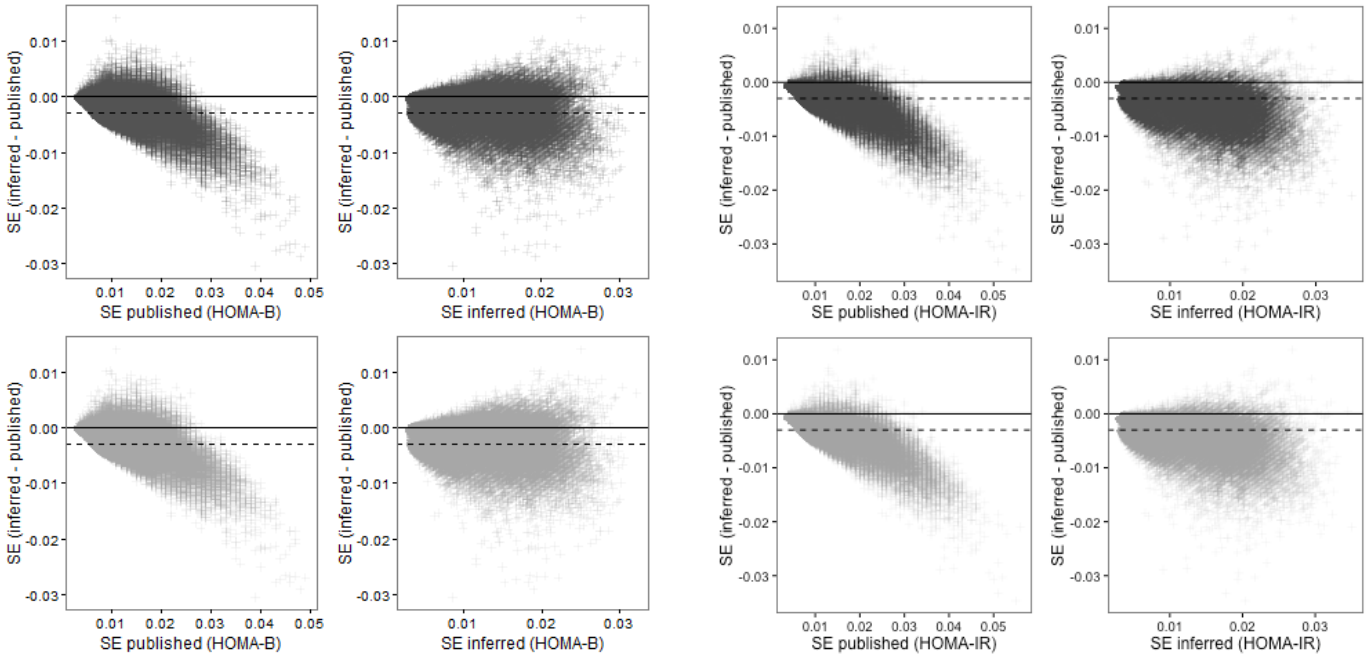

**Figure S16.** -log$_{10}$P of inferred against published meta-analysis results plotted for all SNPs (left) and SNPs, which SE difference between inferred and published GWAS less than -0.003 (right).

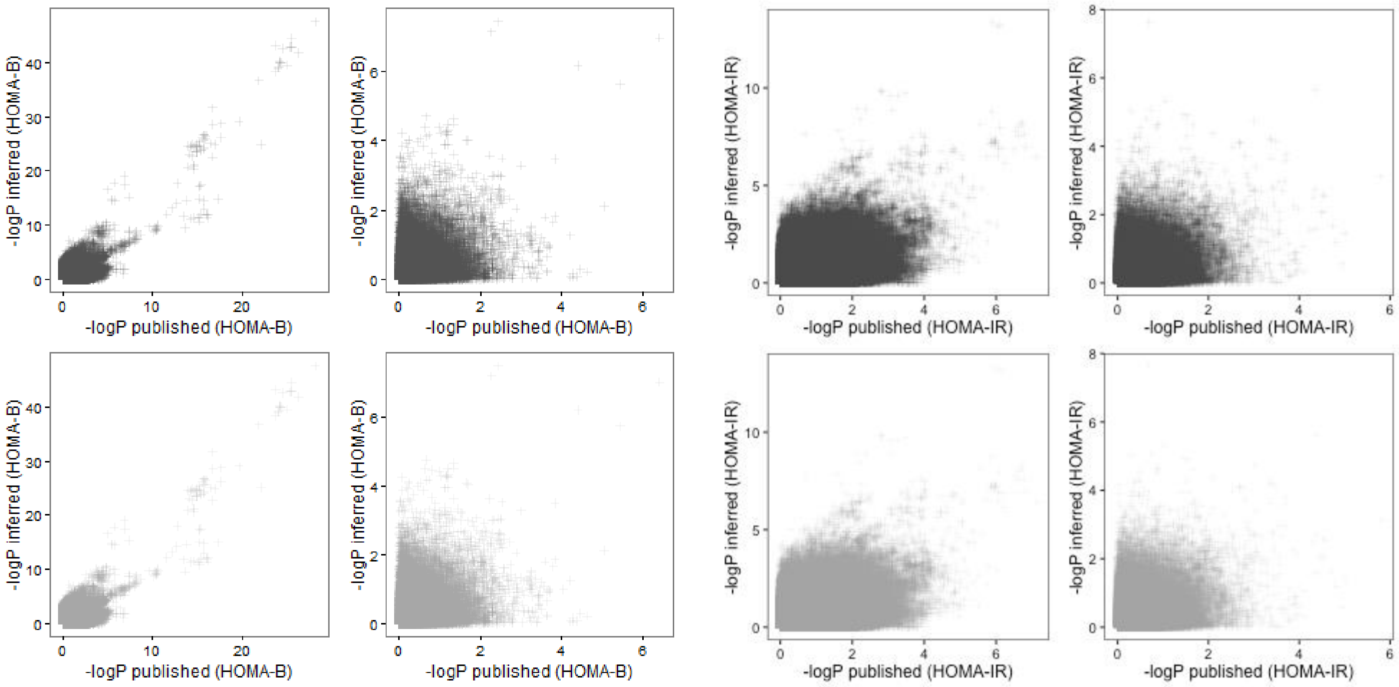

**Figure S17.** Effect size of inferred against published meta-analysis results plotted for all SNPs (left) and SNPs, which SE difference between inferred and published GWAS less than -0.003 (right).

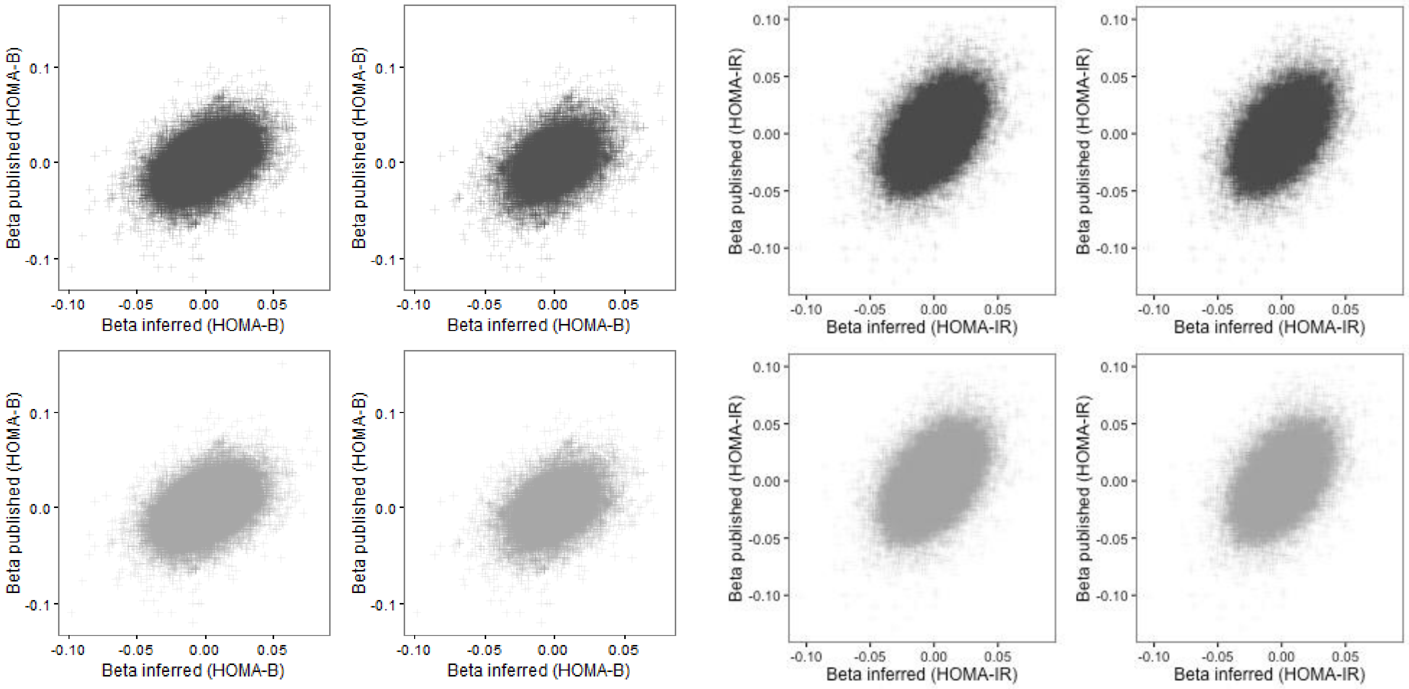

**Figure S18.** Distribution of SE's of the published (upper panel) and inferred (lower panel) results stratified by SE difference < -0.003. Low (light gray) vs. large (dark gray) SEs difference between inferred and published GWAS.

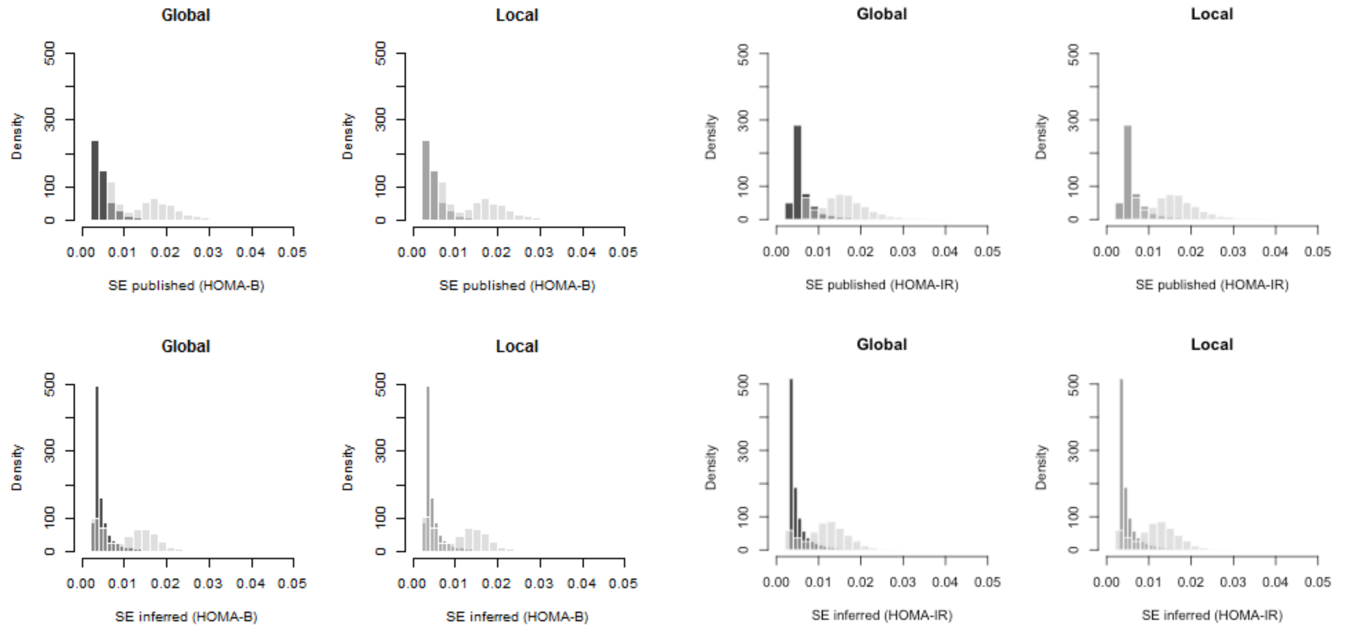


**Figure S19.** Comparison of SE's and effect sizes between 'global' and 'local' correction versions. Upper panel represent the whole dataset, whereas lower panel represent the dataset filtered with N > 35,000 and MAF > 0.01.

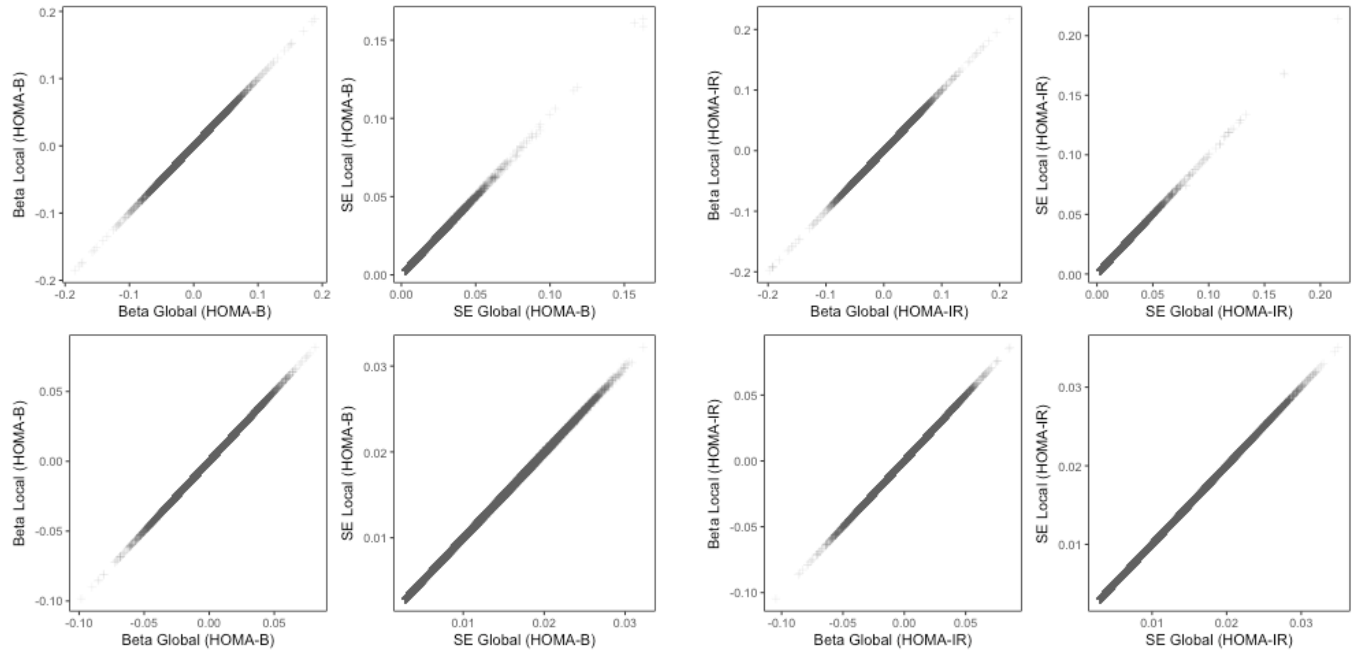

**Figure S20.** Region plots for each of the leading SNPs in HOMA-B GWIS results

a

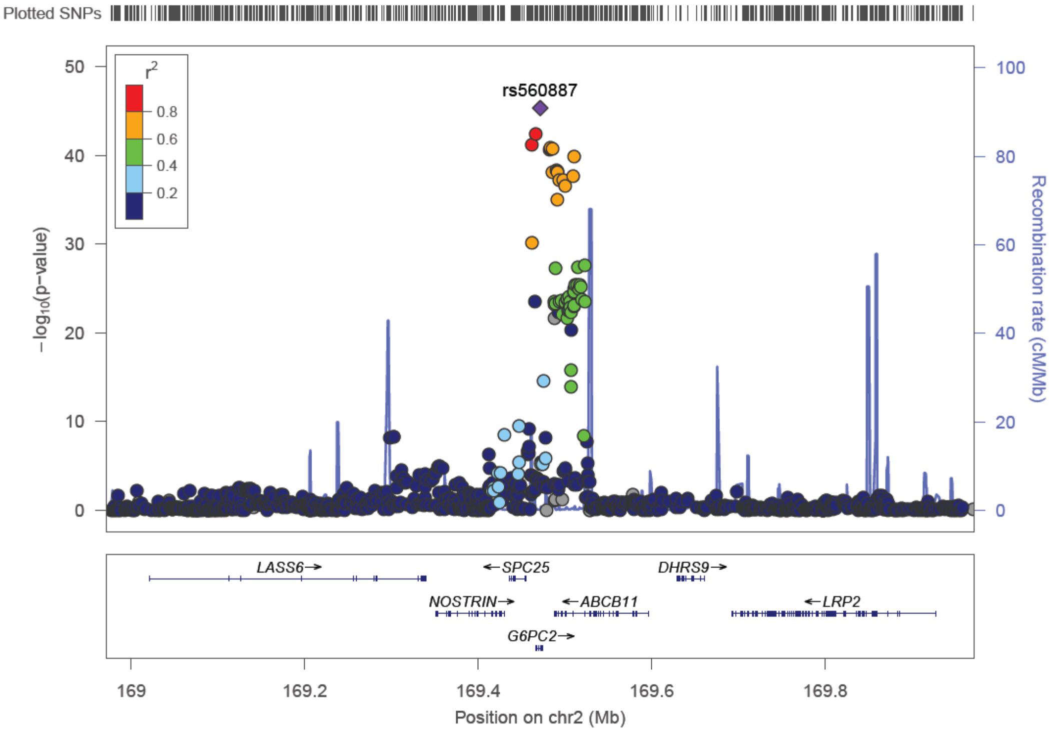


b

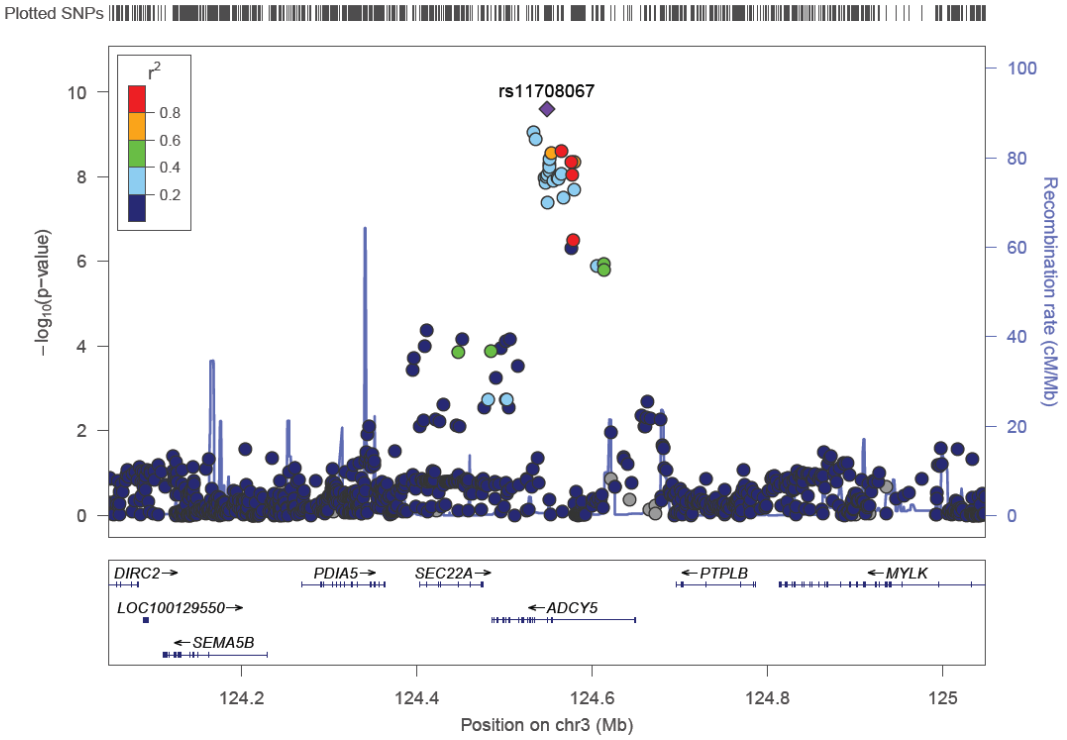

c

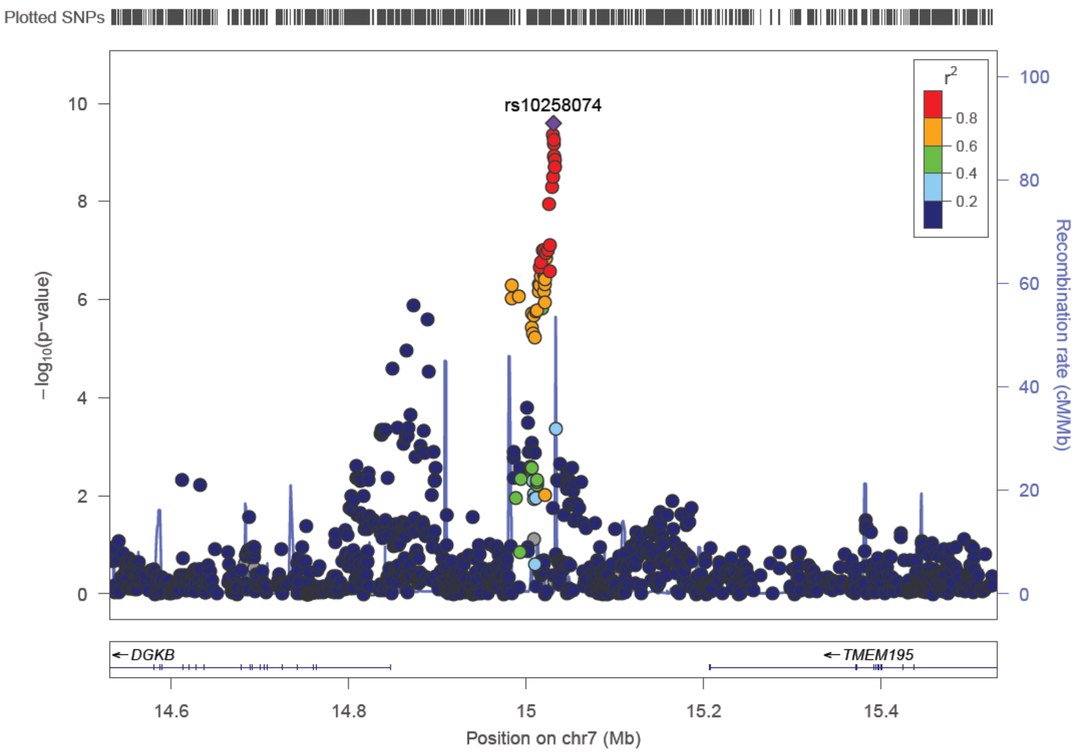

Plotted SNPs
rs10258074
r2
0.8
0.6
0.4
0.2
10
8
6
4
2
0
100
80
60
40
20
0
−log10(p−value)
Recombination rate (cM/Mb)
DGKB
TMEM195
14.6
14.8
15
15.2
15.4
Position on chr7 (Mb)


d

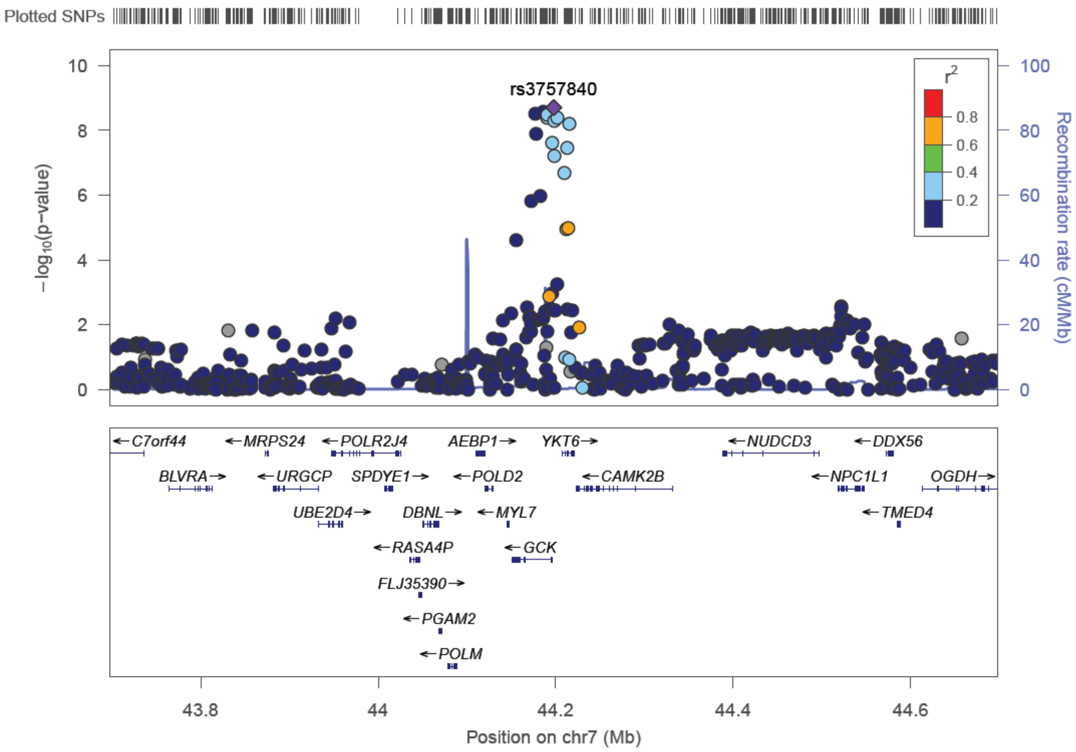

Plotted SNPs
rs3757840
r2
0.8
0.6
0.4
0.2
10
8
6
4
2
0
100
80
60
40
20
0
−log10(p−value)
Recombination rate (cM/Mb)
C7orf44
MRPS24
POLR2J4
AEBP1
YKT6
NUDCD3
DDX56
BLVRA
URGCP
SPDYE1
POLD2
CAMK2B
NPC1L1
OGDH
UBE2D4
DBNL
MYL7
TMED4
RASA4P
GCK
FLJ35390
PGAM2
POLM
43.8
44
44.2
44.4
44.6
Position on chr7 (Mb)

e

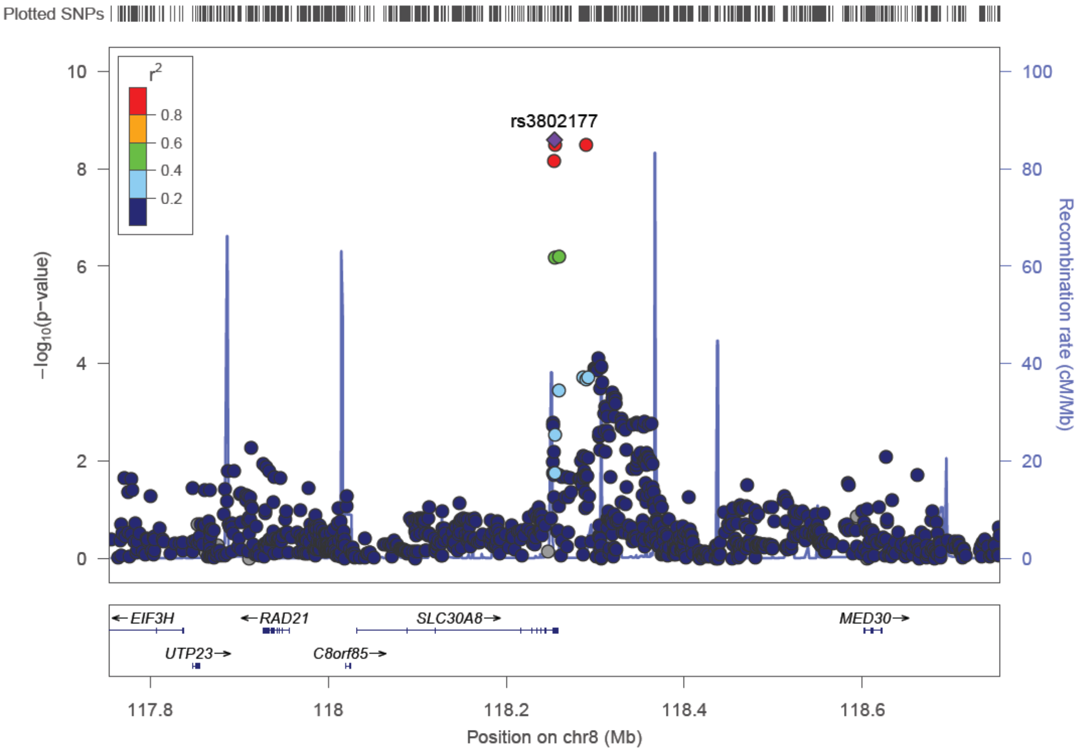

Plotted SNPs
rs3802177
r2
0.8
0.6
0.4
0.2
−log10(p−value)
Recombination rate (cM/Mb)
EIF3H
RAD21
SLC30A8
MED30
UTP23
C8orf85
117.8
118
118.2
118.4
118.6
Position on chr8 (Mb)


f

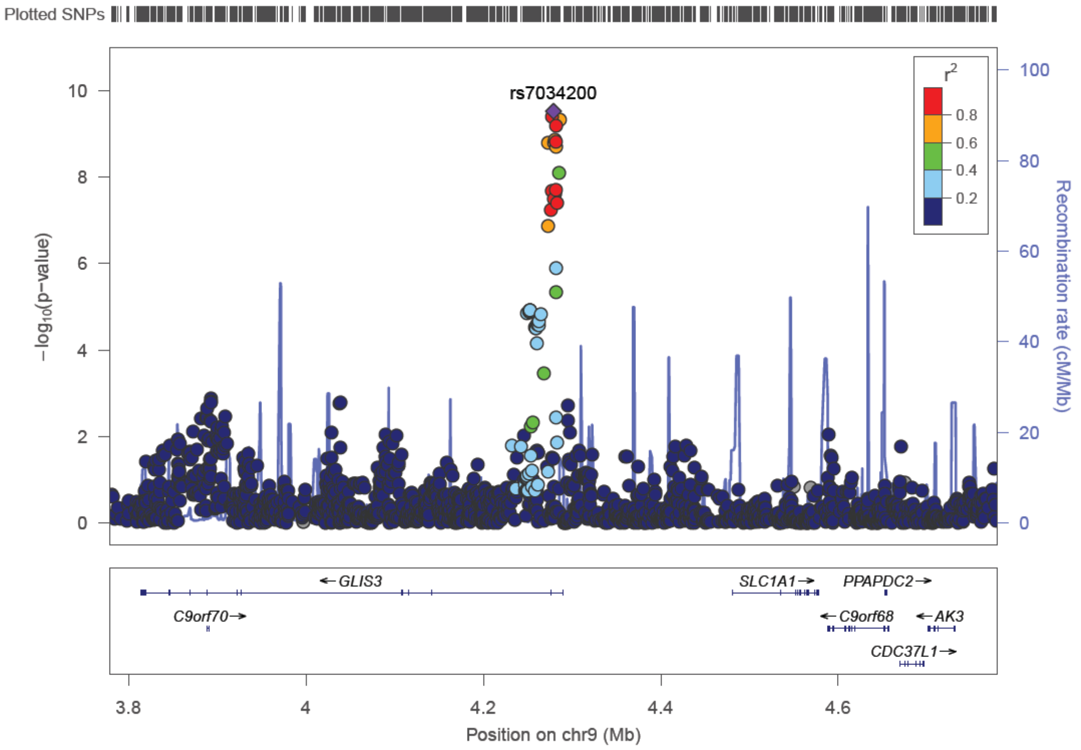

Plotted SNPs
rs7034200
r2
0.8
0.6
0.4
0.2
−log10(p−value)
Recombination rate (cM/Mb)
GLIS3
SLC1A1
PPAPDC2
C9orf70
C9orf68
AK3
CDC37L1
3.8
4
4.2
4.4
4.6
Position on chr9 (Mb)

g

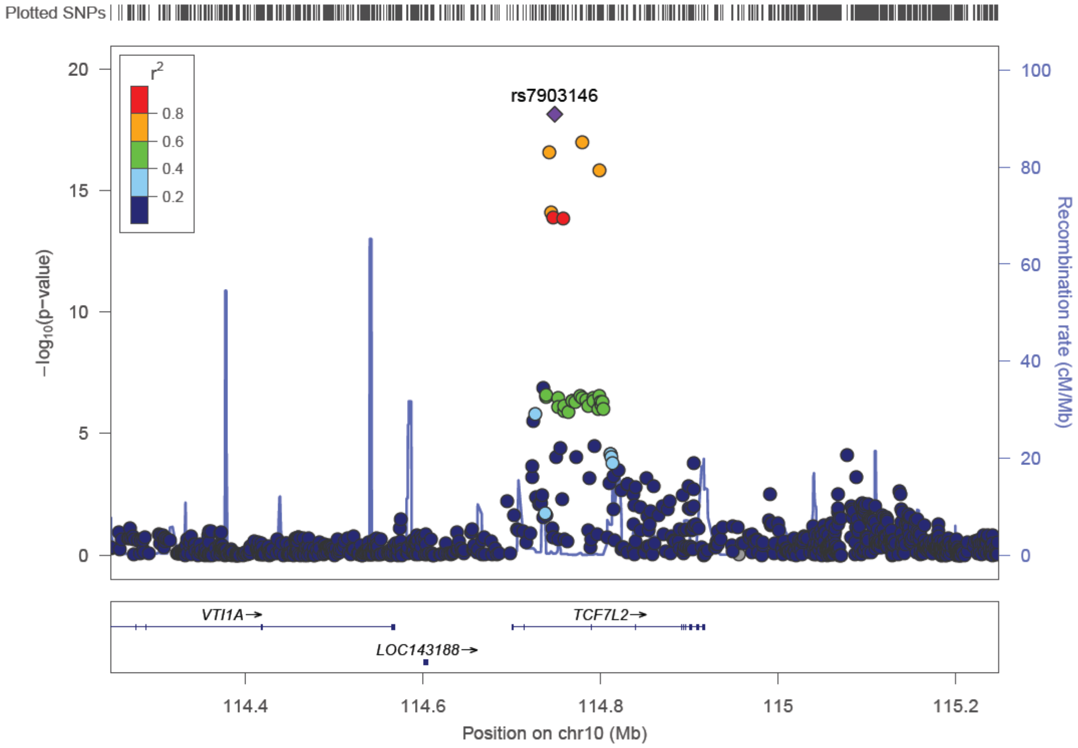
Plotted SNPs
rs7903146
r2
0.8
0.6
0.4
0.2
−log10(p−value)
Recombination rate (cM/Mb)
VTI1A
TCF7L2
LOC143188
Position on chr10 (Mb)

h

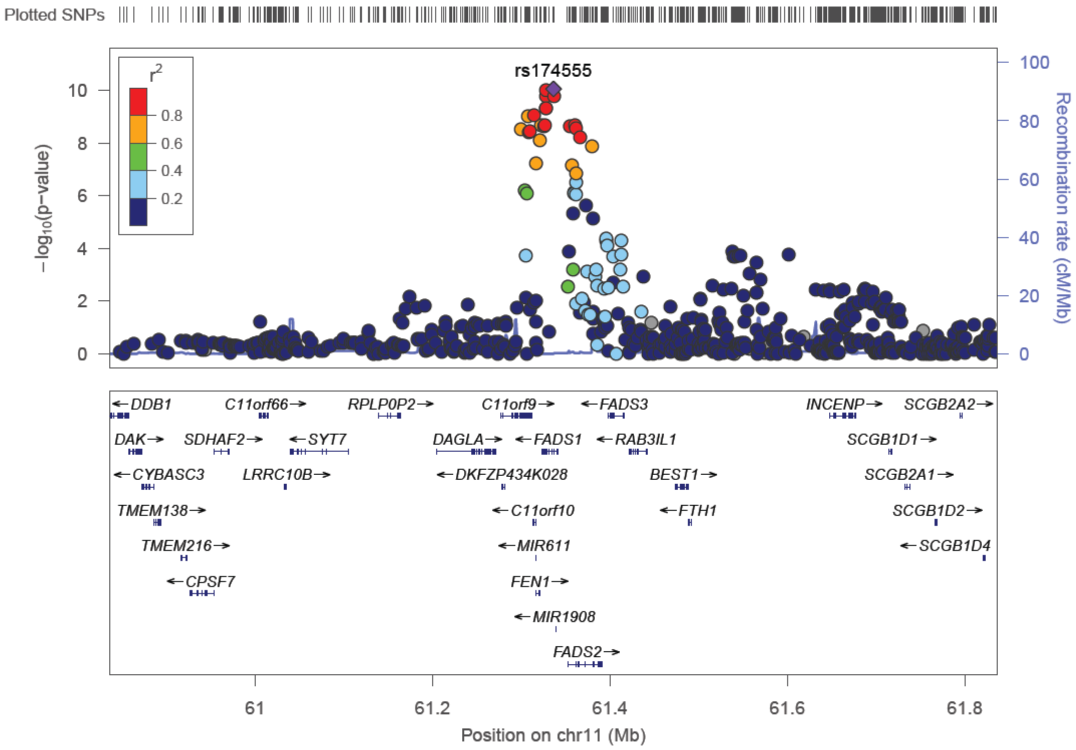
Plotted SNPs
rs174555
r2
−log10(p−value)
Recombination rate (cM/Mb)
DDB1
C11orf66
RPLP0P2
C11orf9
FADS3
INCENP
SCGB2A2
DAK
SDHAF2
SYT7
DAGLA
FADS1
RAB3IL1
SCGB1D1
CYBASC3
LRRC10B
DKFZP434K028
BEST1
SCGB2A1
TMEM138
C11orf10
FTH1
SCGB1D2
TMEM216
MIR611
SCGB1D4
CPSF7
FEN1
MIR1908
FADS2
Position on chr11 (Mb)

i

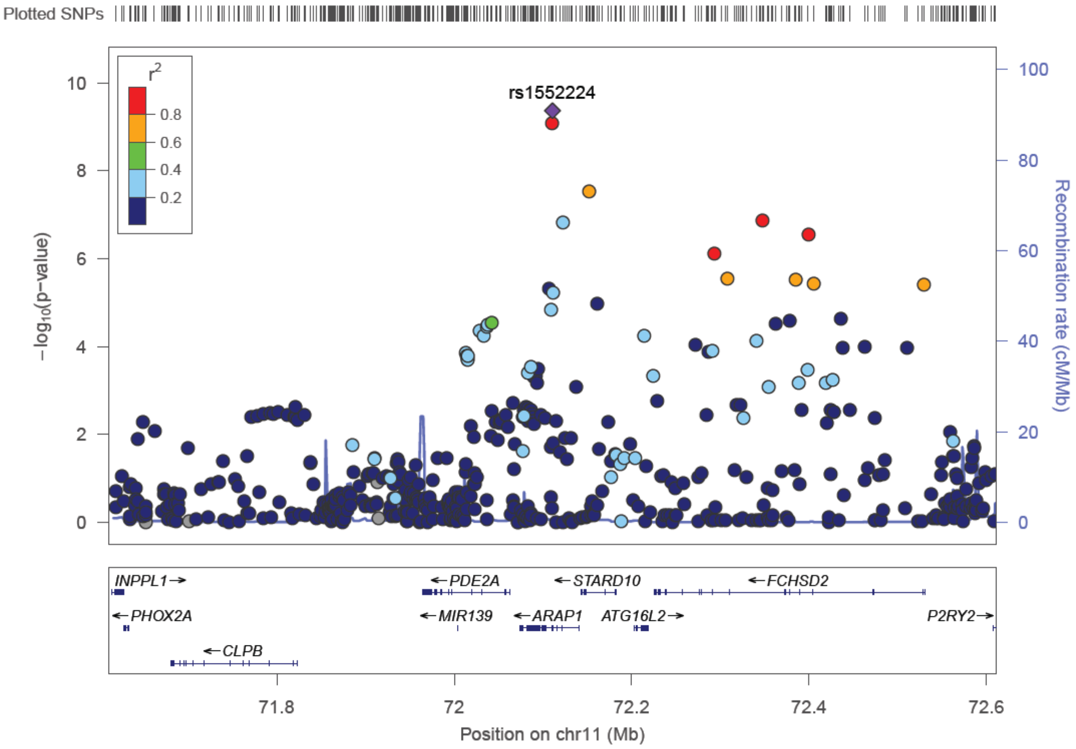
Plotted SNPs
rs1552224
r2
0.8
0.6
0.4
0.2
−log10(p−value)
Recombination rate (cM/Mb)
INPPL1→
←PDE2A
←STARD10
←FCHSD2
←PHOX2A
←MIR139
←ARAP1
ATG16L2→
P2RY2→
←CLPB
71.8
72
72.2
72.4
72.6
Position on chr11 (Mb)

j

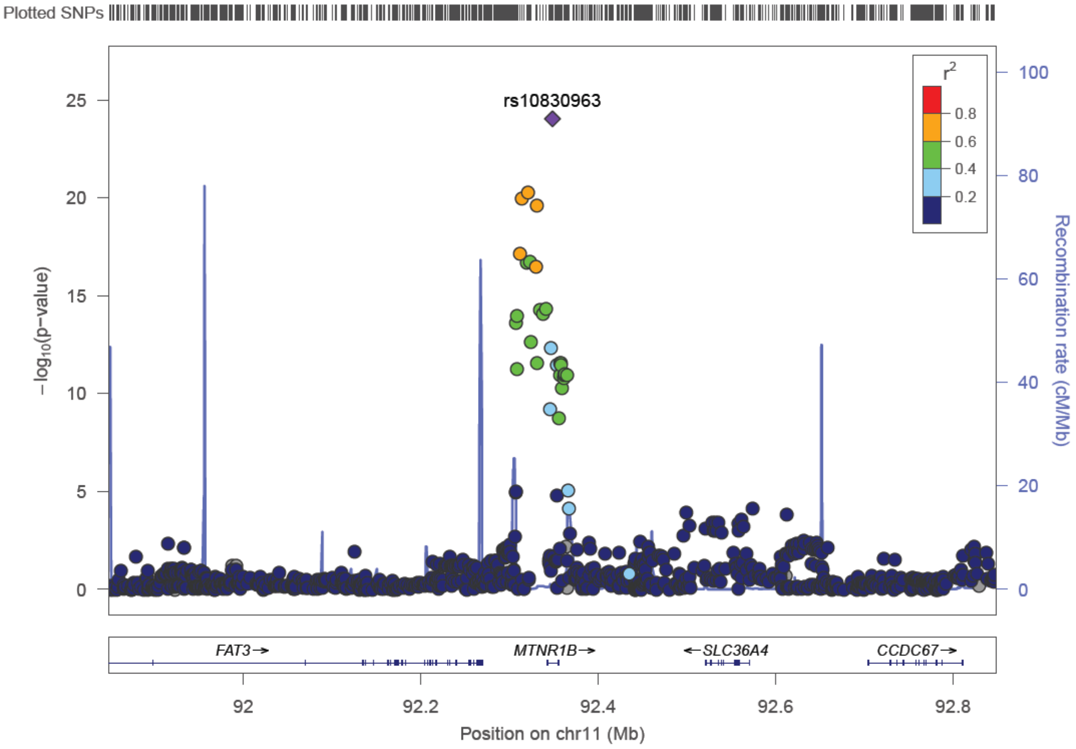
Plotted SNPs
rs10830963
r2
0.8
0.6
0.4
0.2
−log10(p−value)
Recombination rate (cM/Mb)
FAT3→
MTNR1B→
←SLC36A4
CCDC67→
92
92.2
92.4
92.6
92.8
Position on chr11 (Mb)

k

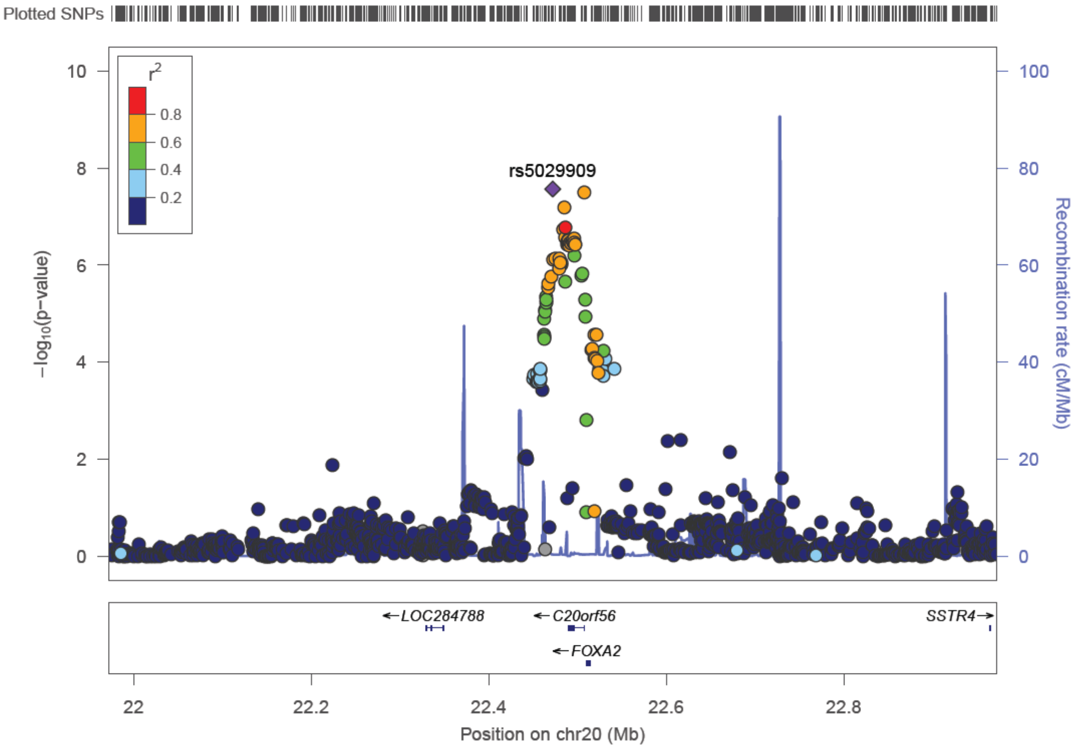
Plotted SNPs
10
8
6
4
2
0
−log10(p−value)
r2
0.8
0.6
0.4
0.2
rs5029909
100
80
60
40
20
0
Recombination rate (cM/Mb)
←LOC284788
←C20orf56
←FOXA2
SSTR4→
22
22.2
22.4
22.6
22.8
Position on chr20 (Mb)

**Figure S21.** Region plots for each of the leading SNPs in HOMA-IR GWIS results.

a

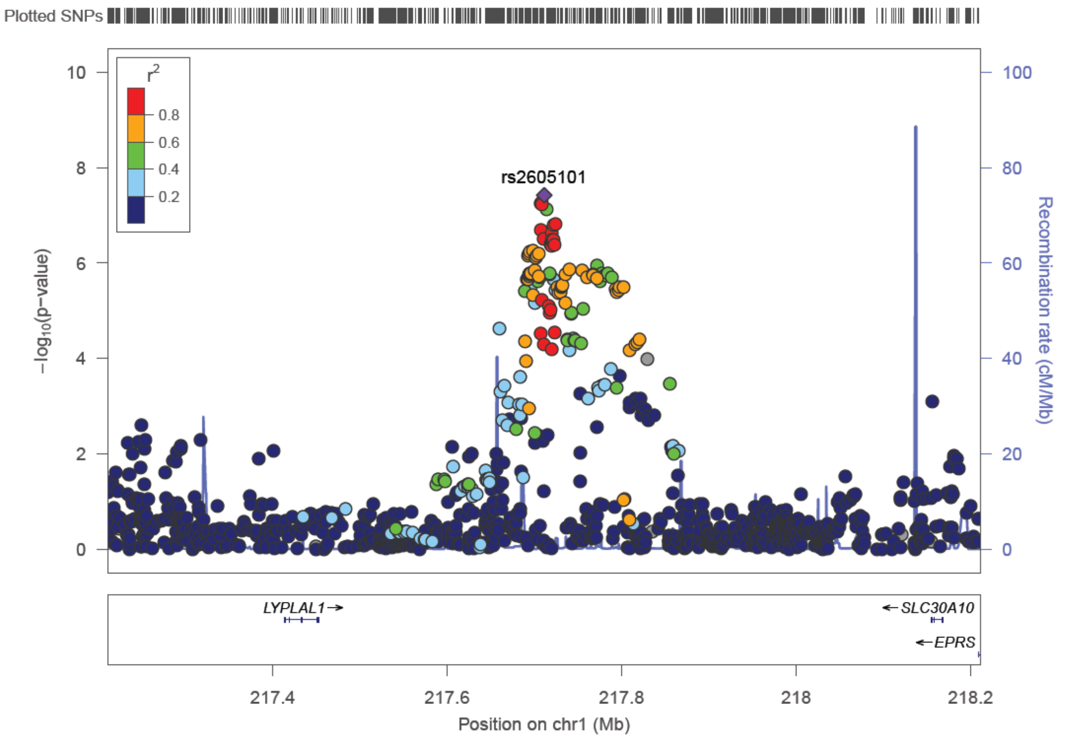


b

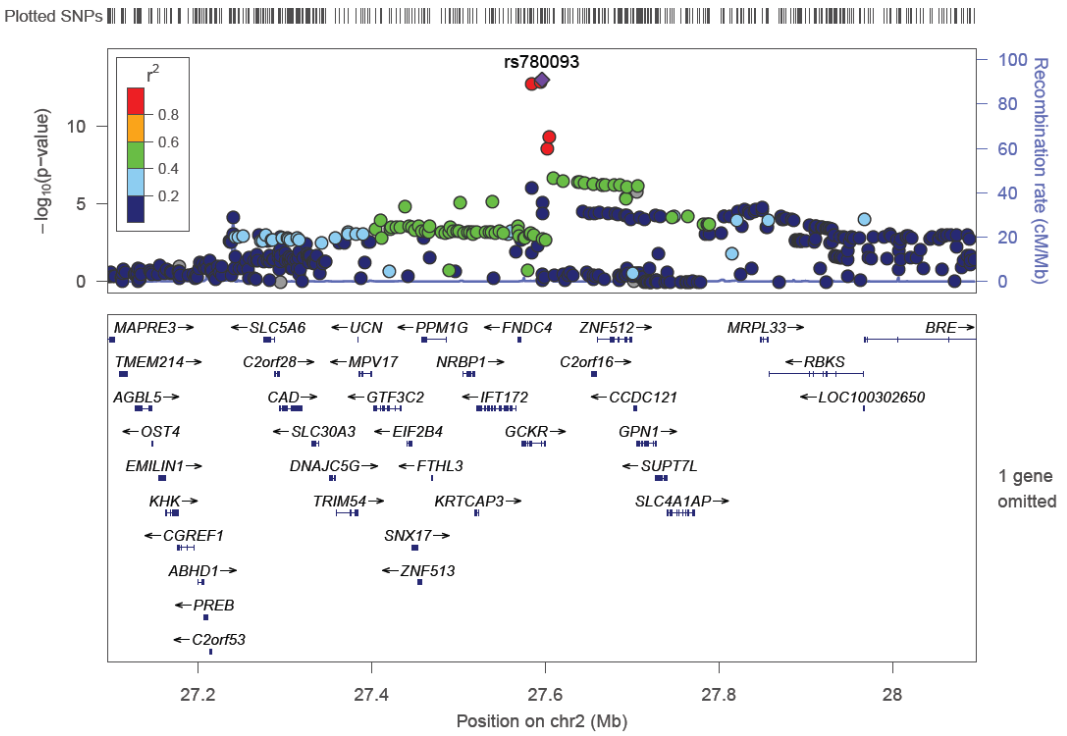

c

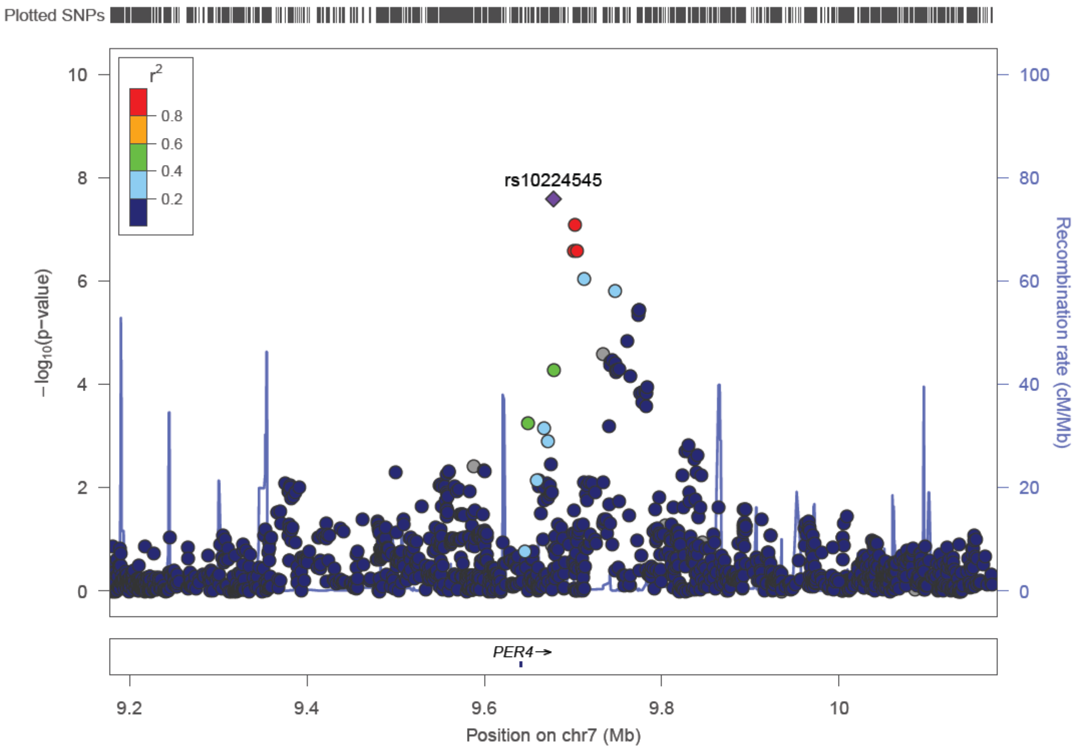
Plotted SNPs
r2
0.8
0.6
0.4
0.2
rs10224545
−log10(p−value)
Recombination rate (cM/Mb)
PER4→
Position on chr7 (Mb)

d

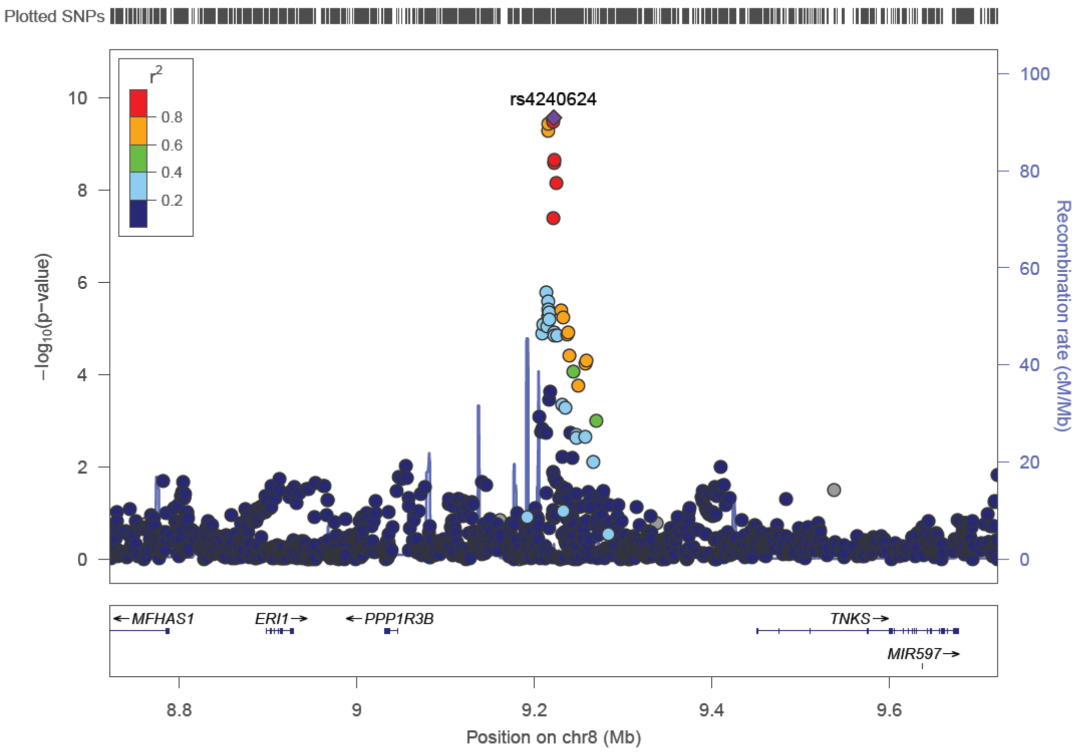
Plotted SNPs
r2
0.8
0.6
0.4
0.2
rs4240624
−log10(p−value)
Recombination rate (cM/Mb)
←MFHAS1
ERI1→
←PPP1R3B
TNKS→
MIR597→
Position on chr8 (Mb)

e

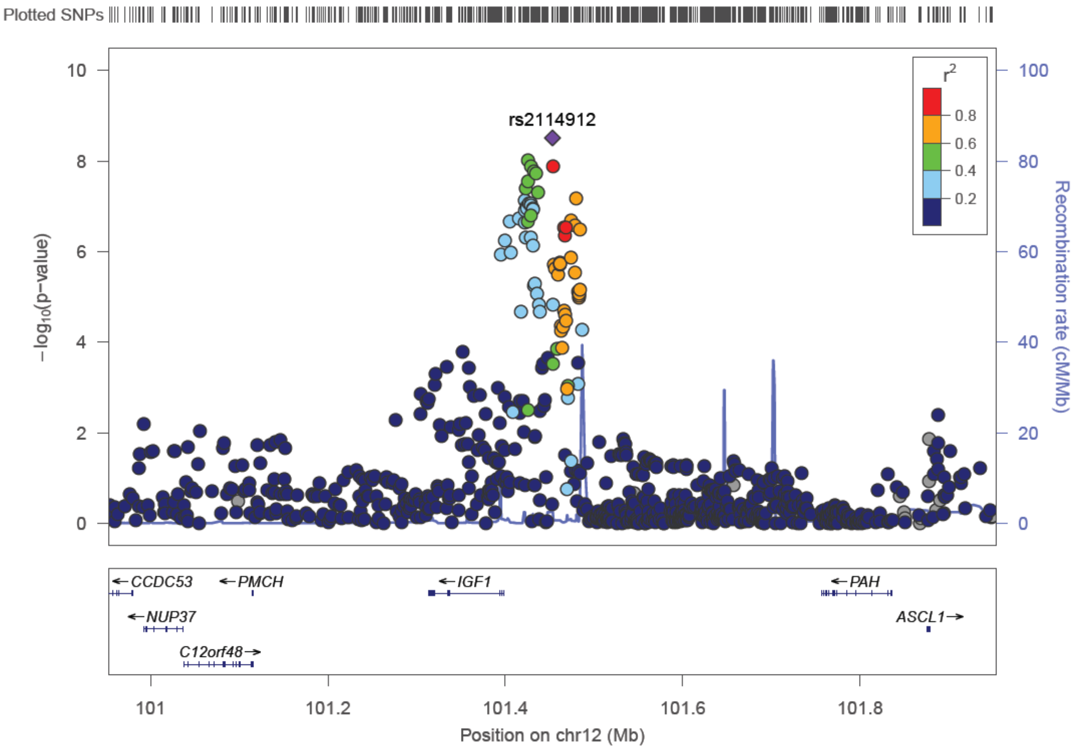

Plotted SNPs
10
8
6
4
2
0
−log10(p−value)
rs2114912
r2
0.8
0.6
0.4
0.2
100
80
60
40
20
0
Recombination rate (cM/Mb)
CCDC53
PMCH
IGF1
PAH
NUP37
ASCL1
C12orf48
101
101.2
101.4
101.6
101.8
Position on chr12 (Mb)

**MAGIC (the Meta-Analyses of Glucose and Insulin-related traits Consortium) investigators**

Vasiliki Lagou[1-3], Reedik Mägi[4], Jouke-Jan J Hottenga[5-6], Harald Grallert[7], John RB Perry[1,8], Nabila Bouatia-Naji[9-11], Letizia Marullo[12], Denis Rybin[13], Rick Janssen[14], Josine L Min[15-16], Antigone S Dimas[17-18], Joao Fadista[19], Maria Stathopoulou[20], Aaron Isaacs[21-23], Sara M Willems[24], Pau Navarro[25], Toshiko Tanaka[26], Anne U Jackson[27], May E Montasser[28], Jeff R O'Connell[28], Lawrence F Bielak[29], Rebecca J Webster[30], Richa Saxena[31-34], Jeanette S Andrews[35], Beate St Pourcain[36], Nicholas J Timpson[37], Perttu Salo[38], So-Youn Shin[39], Najaf Amin[40], Albert V Smith[41-43], Guo Li[44-45], Niek Verweij[46], Anuj Goel[1], Ian Ford[47], Paul CD Johnson[47-48], Toby Johnson[49-50], Karen Kapur[51], Gudmar Thorleifsson[52], Rona Strawbridge[53-54], Laura J Rasmussen-Torvik[55], Tõnu Esko[4,56], Evelin Mihailov[56], Tove Fall[57], Andrea Groop[58-60], Fraser Ross[61], Anubha Mahajan[62], Stavroula Kanoni[63], Vilmantas Giedraitis[64], Marcus E Kleber[65-66], Günther Silbernagel[67], Julia Meyer[68], Martina Müller-Nurasyid[69-71], Andrea Ganna[58-60], Antti-Pekka Sarin[72-73], Loic Yengo[9-10], Dmitry Shungin[74-76], Jian'an Luan[77], Momoko Horikoshi[1,78-79], An Ping[80], Sanna Serena[81-82,82], Yvonne Boettcher[83-84], N W Rayner[1,63,78], Ilja M Nolte[85], Tatijana Zemunik[86], Erik van Iperen[87], Peter Kovacs[88], Nicholas D Hastie[25], Sarah H Wild[61], Stela McLachlan[61], Susan Campbell[25], Ozren Polasek[86], Olga Carlson[89], Josephine Egan[89], Wieland Kiess[84,90], Gonneke Willemsen[5], Johanna Kuusisto[91], Markku Laakso[91], Maria Dimitriou[92], Andrew A Hicks[93], Rainer Rauramaa[94-95], Stefania Bandinelli[96], Barbara Thorand[97], Yongmei Liu[98], Iva Miljkovic[99], Lars Lind[100], Alex Doney[101], Markus

Perola[4,38,102], Aroon Hingorani[103], Mika Kivimaki[103], Meena Kumari[103-104], Amanda J Bennett[78], Christopher J Groves[78], Mark I McCarthy[1,78,105], Christian Herder[106-107], Ulf de Faire[108], Stephan JL Bakker[109], Matti Uusitupa[110], Colin NA Palmer [101], J W Jukema[111], Naveed Sattar[112], Anneli Pouta [113-114], Harold Snieder[85,115], Eric Boerwinkle[116-117], James S Pankow[118], Patrik K Magnusson[119], Ulrika Krus[120], Eco JCN de Geus[5-6], Matthias Blüher[83-84], Bruce HR Wolffenbuttel[115,121], Michael A Province[80], Goncalo R Abecasis[82,122], James B Meigs[123-124], Kees G Hovingh[125], Jaana Lindström[126], James F Wilson[61,127], Alan F Wright[127], George V Dedousis[92], Stefan R Bornstein[128], Peter EH Schwarz[128], Anke Tönjes[83-84], Bernhard R Winkelmann[129], Bernhard O Boehm[130], Winfried März[66,131], Andres Metspalu[4,56], Jackie F Price[61], Panos Deloukas[63,132-133], Antje Körner[84,90], Timo A Lakka[94,134], Sirkka M Keinanen-Kiukaanniemi[135-136], Timo E Saaristo[137-138], Richard N Bergman[139], Jaakko Tuomilehto[140-143], Nicholas J Wareham[77], Claudia Langenberg[77], Satu Männistö[144], Paul W Franks[75,145-146], Caroline Hayward[25], Veronique Vitart[25], Jaako Kaprio[147], Sophie Visvikis-Siest[20], Beverley Balkau[148-149], David Altshuler[31-32,124], Igor Rudan[61], Michael Stumvoll[83-84], Harry Campbell[61], Cornelia M van Duijn[21,150], Christian Gieger[151-153], Thomas Illig[151,154-155], Luigi Ferrucci[26], Nancy L Pedersen[119], Peter P Pramstaller[93,156-157], Michael Boehnke[27], Timothy M Frayling[8], Alan R Shuldiner[28,158], Patricia A Peyser[29], Lyle J Palmer[159], Brenda W Penninx[160], Pierre Meneton[161], Tamara B Harris[162], Gerjan Navis[109], Pim van der Harst[46,163], George Davey Smith[37], Nita G Forouhi[77], Ruth JF Loos[77,164], Veikko Salomaa[165], Nicole Soranzo[39], Dorret I Boomsma[5], Beverley Balkau[148-149], Leif Groop[166-168], Tiinamaija Tuomi[169-171], Albert Hofman[40,172], Patricia B Munroe[49-50], Vilmundur Gudnason[41,173], David S Siscovick[44-45,174], Hugh Watkins[1], Cecile Lecoeur[9-10], Peter Vollenweider[175], Kari

Stefansson[52,176], Marjo-Riitta Jarvelin [177-178], Anders Hamsten[53-54,179], George Nicholson[180], Fredrik Karpe[78,105], Emmanouil T Dermitzakis[18], Cecilia M Lindgren[1,78,181], Philippe Froguel[9-10,182], Valeriya Lyssenko[168,183], Richard M Watanabe[184-186], Erik Ingelsson[57,187], Jose C Florez[32,188], Josée Dupuis[189-190], Inês Barroso[191-192], Andrew P Morris[1,4,193], Inga Prokopenko[1,78,182].

**Affiliations:**

1)Wellcome Centre for Human genetics, University of Oxford, Oxford, United Kingdom; 2)Laboratory for Neuroimmunology, Department of Neurosciences, KU Leuven, Leuven, Belgium; VIB Center for Brain & Disease Research, Leuven, Belgium; 3)Laboratory for Translational Immunology, Department of Immunology and Microbiology, KU Leuven, Leuven, Belgium; 4)Estonian Genome Center, University of Tartu, Tartu, Estonia; 5)Department of Biological Psychology, Vrije Universiteit, Amsterdam, the Netherlands; 6)Amsterdam Public Health research institute, VU University medical center, Amsterdam, the Netherlands; 7)Research Unit Molecular Epidemiology, Helmholtz Zentrum München, German Research Center for Environmental Health, Neuherberg, Germany; 8)Genetics of Complex Traits, Peninsula Medical School, University of Exeter, United Kingdom; 9)University of Lille Nord de France, Lille, France; 10)CNRS UMR8199, Institut Pasteur de Lille, Lille, France; 11)INSERM U970, Paris Cardiovascular Research Center PARCC, 75006 Paris, France; 12)Department of Life Sciences and Biotechnology, University of Ferrara, Ferrara, Italy; 13)Boston University Data Coordinating Center, Boston, Massachusetts, USA; 14)Department of Psychiatry, VU University Medical Center Amsterdam, Amsterdam, the Netherlands; 15)MRC Integrative Epidemiology

Unit, University of Bristol, Bristol, United Kingdom; 16)Bristol Medical School, University of Bristol, Bristol, United Kingdom; 17)Biomedical Sciences Research Center "Alexander Fleming", Vari, Greece; 18)Department of Genetic Medicine and Development, University of Geneva Medical School, Geneva, Switzerland; 19)Department of Epidemiology Research, Statens Serum Institut, Copenhagen, Denmark; 20)UMR INSERM U1122; Interactions Gène-Environnement en Physiopathologie Cardio-Vasculaire (IGE-PCV), Université de Lorraine, Nancy, France; 21)Genetic Epidemiology Unit, Department of Epidemiology, Erasmus Medical Center, Rotterdam, the Netherlands; 22)CARIM School for Cardiovascular Diseases, Maastricht Centre for Systems Biology (MaCSBio), Maastricht University, Maastricht, the Netherlands; 23)Department of Biochemistry, Maastricht University, Maastricht, the Netherlands; 24)Genetic Epidemiology Unit, Department of Epidemiology, Erasmus University Medical Center, Rotterdam, the Netherlands; 25)MRC Human Genetics Unit, MRC Institute of Genetics and Molecular Medicine,University of Edinburgh, Western General Hospital, Edinburgh, United Kingdom; 26)Clinical Research Branch, National Institute on Aging, Baltimore, Maryland, USA; 27)Department of Biostatistics and Center for Statistical Genetics, University of Michigan, Ann Arbor, Michigan, USA; 28)Division of Endocrinology, Diabetes, and Nutrition, Department of Medicine, University of Maryland, School of Medicine, Baltimore, Maryland, USA; 29)Department of Epidemiology, University of Michigan, Ann Arbor, Michigan, USA; 30)Laboratory for Cancer Medicine, Harry Perkins Institute of Medical Research, University of Western Australia Centre for Medical Research, Nedlands, Australia; 31)Broad Institute of Harvard and Massachusetts Institute of Technology (MIT), Cambridge, Massachusetts, USA; 32)Center for

Human Genetic Research, Massachusetts General Hospital, Boston, Massachusetts, USA; 33)Department of Genetics, Harvard Medical School, Boston, Massachusetts, USA; 34)Departmentartment of Anesthesia, Critical Care and Pain Medicine, MGH, Boston, USA; 35)Department of Biostatistical Sciences, Division of Public Health Sciences, Wake Forest University School of Medicine, Winston-Salem, North Carolina, USA; 36)School of Social and Community Medicine, University of Bristol, Bristol, United Kingdom; 37)MRC CAiTE Centre, School of Social and Community Medicine, University of Bristol, Bristol, United Kingdom; 38)Public Health Genomics Unit, Department of Chronic Disease Prevention, the National Institute for Health and Welfare, Helsinki, Finland; 39)Wellcome Trust Sanger Institute, Wellcome Trust Genome Campus, Hinxton, United Kingdom; 40)Department of Epidemiology Erasmus MC, Rotterdam, the Netherlands; 41)Icelandic Heart Association, Kopavogur, Iceland; 42)Faculty of MedicineUniversity of Iceland, Reykjavik, Iceland; 43)Department of Biostatistics, School of Public Health,University of Michigan, Ann Arbor, Michigan, USA; 44)Cardiovascular Health Research Unit, University of Washington, Seattle, Washington, USA; 45)Department of Medicine, University of Washington, Seattle, Washington, USA; 46)Department of Cardiology, University Medical Center Groningen, University of Groningen, Groningen, the Netherlands; 47)Robertson Centre for Biostatistics, University of Glasgow, Glasgow, United Kingdom; 48)Institute of Biodiversity, Animal Health & Comparative Medicine, University of Glasgow, Glasgow, United Kingdom; 49)Clinical Pharmacology, William Harvey Research Institute, Barts and The London School of Medicine and Dentistry, Queen Mary University of London, London, United Kingdom; 50)NIHR Barts Cardiovascular Biomedical Research Unit, Barts and The London School of

Medicine and Dentistry, Queen Mary University of London, London, United Kingdom; 51)Department of Medical Genetics, University of Lausanne, Lausanne, Switzerland; 52)deCODE Genetics, Reykjavik, Iceland; 53)Cardiovascular Medicine Unit, Department of Medicine, Solna, Karolinska Institutet, Stockholm, Sweden; 54)Center for Molecular Medicine, Karolinska University Hospital Solna, Stockholm, Sweden; 55)Department of Preventive Medicine, Northwestern University Feinberg School of Medicine, Chicago, Illinois, USA; 56)Insitute of Molecular and Cell Biology, University of Tartu, Tartu, Estonia; 57)Department of Medical Sciences, Molecular Epidemiology and Science for Life Laboratory, Uppsala University, Uppsala, Sweden; 58)Analytic and Translational Genetics Unit, Massachusetts General Hospital, Boston, Massachusetts, USA; 59)Program in Medical and Population Genetics, Broad Institute of MIT and Harvard, Cambridge, Massachussets, USA; 60)Stanley Center for Psychiatric Research, Broad Institute of MIT and Harvard, Cambridge, Massachussets, USA; 61)Usher Institute of Population Health Sciences and Informatics, University of Edinburgh, Edinburgh, United Kingdom; 62)Wellcome Centre for Human Genetics, University of Oxford, Oxford, United Kingdom; 63)Wellcome Trust Sanger Institute, Hinxton, United Kingdom; 64)Department of Public Health and Caring Sciences, Uppsala Universitet, Uppsala, Sweden; 65)LURIC Study nonprofit LLC, Freiburg, Germany; 66)Mannheim Institute of Public Health, Social and Preventive Medicine, Medical Faculty of Mannheim, University of Heidelberg, Mannheim, Germany; 67)Division of Angiology, Department of Internal Medicine, Medical University of Graz, Austria; 68)Institute of Genetic Epidemiology, Helmholtz Zentrum München, German Research Center for Environmental Health, Neuherberg, Germany; 69)Institute of Medical Informatics, Biometry and

Epidemiology, Chair of Epidemiology and Chair of Genetic Epidemiology, Ludwig-Maximilians-Universität, Munich, Germany; 70)Department of Medicine I, University Hospital Grosshadern, Ludwig-Maximilians-University, Munich, Germany; 71)Institute of Genetic Epidemiology, Helmholtz Zentrum München, German Research Center for Environmental Health, Neuherberg, Germany; 72)Institute for Molecular Medicine Finland, FIMM, University of Helsinki, Finland; 73)Public Health Genomics Unit, National Institute for Health and Welfare, Helsinki, Finland; 74)Department of Public Health & Clinical Medicine, Umeå University, Umeå, Sweden; 75)Department of Clinical Sciences, Genetic and Molecular Epidemiology Unit, Skåne University Hospital Malmö, Malmö, Sweden; 76)Department of Odontology, Umeå University, Umeå, Sweden; 77)MRC Epidemiology Unit, University of Cambridge School of Clinical Medicine, Cambridge, United Kingdom; 78)Oxford Centre for Diabetes, Endocrinology and Metabolism, University of Oxford, Oxford, United Kingdom; 79)RIKEN, Center for Integrative Medical Sciences, Laboratory for Endocrinology, Metabolism and kidney disease, Yokohama, Japan; 80)Division of Statistical Genomics, Washington University School of Medicine, St. Louis, Missouri, USA; 81)Istituto di Ricerca Genetica e Biomedica, CNR, Monserrato, Italy; 82) ; 83)University of Leipzig, Department of Medicine, Leipzig, Germany; 84)University of Leipzig, IFB AdiposityDiseases, Leipzig, Germany; 85)Department of Epidemiology, University Medical Center Groningen, University of Groningen, Groningen, the Netherlands; 86)Faculty of Medicine, University of Split, Split, Croatia; 87)Department of Clinical Epidemiology and Biostatistics, Academic Medical Center, University of Amsterdam, Amsterdam, the Netherlands; 88)Interdisciplinary Center for Clinical Research, University of Leipzig, Leipzig,

Germany; 89)Laboratory of Clinical Investigation, National Institute of Aging, Baltimore, Maryland, USA; 90)Pediatric Research Center, Department of Women´s & Child Health, University of Leipzig, Leipzig, Germany; 91)Department of Medicine, University of Eastern Finland and Kuopio University Hospital, Kuopio, Finland; 92)Department of Dietetics-Nutrition, Harokopio University, Athens, Greece; 93)Center for Biomedicine, European Academy Bozen/Bolzano (EURAC), Bolzano, Italy - Affiliated Institute of the University of Lübeck, Lübeck, Germany; 94)Kuopio Research Institute of Exercise Medicine, Kuopio, Finland; 95)Department of Clinical Physiology and Nuclear Medicine, Kuopio University Hospital, Kuopio, Finland; 96)Geriatric Unit, Azienda Sanitaria Firenze (ASF), Florence, Italy; 97)Institute of Epidemiology II, Helmholtz Zentrum München, German Research Center for Environmental Health, Neuherberg, Germany; 98)Department of Epidemiology and Prevention, Division of Public Health Sciences, Wake Forest University School of Medicine, Winston-Salem, North Carolina, USA; 99)Department of Epidemiology, Center for Aging and Population Health, University of Pittsburgh, Pittsburgh, Pennsylvania, USA; 100)Department of Medical Sciences, Uppsala University, Akademiska sjukhuset, Uppsala, Sweden; 101)Pat McPherson Centre for Pharmacogenetics and Pharmacogenomics, Division of Molecular and Clinical Medicine, Ninewells Hospital and Medical School, University of Dundee, Dundee, United Kingdom; 102)Institute for Molecular Medicine Finland FIMM, University of Helsinki, Finland; 103)Department of Epidemiology and Public Health, University College London, London, United Kingdom; 104)University of Essex, Wivenhoe Park, Colchester, Essex, United Kingdom; 105)Oxford National Institute for Health Research Biomedical Research Centre, Churchill Hospital, Oxford, United

Kingdom; 106)Institute for Clinical Diabetology, German Diabetes Center, Leibniz Center for Diabetes Research, Heinrich Heine University Düsseldorf, Düsseldorf, Germany; 107)German Center for Diabetes Research (DZD), München-Neuherberg, Germany; 108)Division of Cardiovascular Epidemiology, Institute of Environmental Medicine, Karolinska Institutet, Stockholm, Sweden; 109)Department of Internal Medicine, University Medical Center Groningen, University of Groningen, the Netherlands; 110)Institute of Public Health and Clinical Nutrition, University of Eastern Finland, Kuopio, Finland; 111)Department of Cardiology C5-P, Leiden University Medical Center, Leiden, the Netherlands; 112)Institute of Cardiovascular and Medical Sciences, University of Glasgow, Glasgow, United Kingdom; 113)National Institute for Health and Welfare, Oulu, Finland; 114)Department of Clinical Sciences/Obstetrics and Gynecology, University of Oulu, Oulu, Finland; 115)Lifelines Cohort Study and Biobank, Groningen, the Netherlands; 116)IMM Center for Human Genetics, University of Texas Health Science Center at Houston, Houston, Texas, USA; 117)Division of Epidemiology, School of Public Health, University of Texas Health Science Center at Houston, Houston, Texas, USA; 118)Division of Epidemiology and Community Health, School of Public Health, University of Minnesota, Minneapolis, Minnesota, USA; 119)Department of Medical Epidemiology and Biostatistics, Karolinska Institutet, Stockholm, Sweden; 120)Department of Clinical Sciences, Diabetes and Endocrinology Research Unit, University Hospital Malmö, Lund University, Malmö, Sweden ; 121)Department of Endocrinology, University Medical Center Groningen, University of Groningen, Groningen, the Netherlands; 122)Center for Statistical Genetics, Department of Biostatistics, University of Michigan, Ann Arbor,

Michigan, USA; 123)General Medicine Division, Massachusetts General Hospital, Boston, Massachusetts, USA; 124)Department of Medicine, Harvard Medical School, Boston, Massachusetts, USA; 125)Department Vascular Medicine, Academic Medical Center, Amsterdam, the Netherlands; 126)National Institute for Health and Welfare, Diabetes Prevention Unit, Helsinki, Finland; 127)MRC Human Genetics Unit, MRC Institute of Genetics and Molecular Medicine, University of Edinburgh, Western General Hospital, Edinburgh, United Kingdom; 128)Department of Medicine III, University of Dresden, Medical Faculty Carl Gustav Carus, Dresden, Germany; 129)Cardiology Group, Frankfurt-Sachsenhausen, Germany; 130)Division of Endocrinology and Diabetes, Department of Medicine, University Hospital, Ulm, Germany; 131)Synlab Academy, Mannheim, Germany; 132)William Harvey Research Institute, Barts and The London School of Medicine and Dentistry, Queen Mary University of London, London, United Kingdom; 133)Princess Al-Jawhara Al-Brahim Centre of Excellence in Research of Hereditary Disorders (PACER-HD), King Abdulaziz University, Jeddah, Saudi Arabia; 134)Institute of Biomedicine/Physiology, University of Eastern Finland, Kuopio Campus, Kuopio, Finland; 135)Faculty of Medicine, Institute of Health Sciences, University of Oulu, Oulu, Finland; 136)Unit of General Practice, Oulu University Hospital, Oulu, Finland; 137)Finnish Diabetes Association, Tampere, Finland; 138)Pirkanmaa Hospital District, Tampere, Finland; 139)Diabetes and Obesity Research Institute, Cedars-Sinai Medical Center, Los Angeles, California, USA; 140)Department of Chronic Disease Prevention, National Institute for Health and Welfare, Helsinki, Finland; 141)Dasman Diabetes Institute, Dasman, Kuwait; 142)Centre for Vascular Prevention, Danube-University Krems, Krems, Austria; 143)Diabetes Research

Group, King Abdulaziz University, Jeddah, Saudi Arabia; 144)Department of Public Health Solutions, National Institute for Health and Welfare, Helsinki, Finland; 145)Department of Nutrition, Harvard School of Public Health, Boston, Massachusetts, USA; 146)Department of Public Health & Clinical Medicine, Units of Medicine and Nutritional Research, Umeå University, Umeå, Sweden; 147)Department of Public Health, University of Helsinki, Helsinki, Finland; 148)Inserm, CESP Center for Research in Epidemiology and Public Health, U1018, Epidemiology of diabetes, obesity and chronic kidney disease over the lifecourse, Villejuif, France; 149)University Paris Sud 11, UMRS 1018, Villejuif, France; 150)Centre for Medical Systems Biology, Leiden, the Netherlands; 151)Research Unit of Molecular Epidemiology, Helmholtz Zentrum München, German Research Center for Environmental Health, Neuherberg, Germany; 152)Institute of Epidemiology II, Helmholtz Zentrum München, German Research Center for Environmental Health, Neuherberg, Germany; 153)German Center for Diabetes Research (DZD), Neuherberg, Germany; 154)Hannover Unified Biobank, Hannover Medical School, Hannover, Germany; 155)Institute of Human Genetics, Hannover Medical School, Hannover, Germany; 156)Department of Neurology, General Central Hospital, Bolzano, Italy; 157)Department of Neurology, University of Lübeck, Lübeck, Germany; 158)Geriatric Research and Education Clinical Center, Veterans Administration Medical Center, Baltimore, Maryland; 159)School of Public Health, University of Adelaide, Adelaide, Australia; 160)Department of Psychiatry, VU University Medical Center, Amsterdam, the Netherlands; 161)U872 Institut National de la Santé et de la Recherche Médicale, Centre de Recherche des Cordeliers, 75006 75006 Paris, France; 162)Geriatric Epidemiology Section, Laboratory of

Epidemiology, Demography, and Biometry, National Institute on Aging, Bethesda, Maryland; 163)Department of Genetics, University Medical Center Groningen, University of Groningen, Groningen, the Netherlands; 164)The Charles Bronfman Institute for Personalized Medicine, Icahn School of Medicine at Mount Sinai, New York, NY,  USA; 165)Chronic Disease Epidemiology and Prevention Unit, Department of Chronic Disease Prevention, National Institute for Health and Welfare, Helsinki, Finland; 166)Institute for Molecular Medicine, Helsinki, Finland ; 167)Department of Medicine, Helsinki University Hospital, University of Helsinki, Helsinki, Finland; 168)Department of Clinical Sciences, Diabetes and Endocrinology Research Unit, University Hospital Malmö, Lund University, Malmö, Sweden; 169)Endocrinology, Abdominal Centre, University of Helsinki and Helsinki University Hospital, Helsinki, Finland; 170)Diabetes and Obesity Research Program, University of Helsinki and Folkhälsan Research Center, Helsinki, Finland; 171)Finnish Institute for Molecular Medicine, University of Helsinki, Helsinki, Finland; 172)Netherlands Consortium for healthy ageing, the Hague, the Netherlands; 173)Faculty of Medicien University of Iceland, Reykjavik, Iceland; 174)Department of Epidemiology, University of Washington, Seattle, Washington, USA; 175)Department of Medicine, University Hospital Lausanne, Lausanne, Switzerland; 176)Faculty of Medicine, University of Iceland, Reykjavík, Iceland; 177)Department of Epidemiology and Biostatistics and HPA-MRC Center, School of Public Health, Imperial College London, London, United Kingdom; 178)Institue of Health Sciences, University of Oulu, Finland; 179)Department of Cardiology, Karolinska University Hospital Solna, Stockholm, Sweden; 180)Department of Statistics, University of Oxford, Oxford, United Kingdom; 181)Big Data Institute, Li Ka Shing Centre for Health Information and

Discovery, University of Oxford, Oxford, United Kingdom; 182)Department of Medicine, Imperial College London, London, United Kingdom; 183)Department of Clinical Science, University of Bergen, Bergen, Norway; 184)Department of Preventive Medicine, Keck School of Medicine of USC, Los Angeles, California, USA; 185)Department of Physiology & Neuroscience, Keck School of Medicine of USC, Los Angeles, California, USA; 186)USC Diabetes and Obesity Research Institute, Los Angeles, California, USA; 187)Department of Medicine, Division of Cardiovascular Medicine, Stanford University School of Medicine, Stanford, California, USA; 188)Diabetes Research Center, Diabetes Unit, Massachusetts General Hospital, Boston, Massachusetts, USA; 189)Department of Biostatistics, Boston University School of Public Health, Boston, Massachusetts, USA; 190)National Heart, Lung, and Blood Institute's Framingham Heart Study, Framingham, Massachusetts, USA; 191)Wellcome Trust Sanger Institute, Wellcome Trust Genome Campus,Hinxton, United Kingdom; 192)University of Cambridge Metabolic Research Laboratories and NIHR Cambridge Biomedical Research Centre, Wellcome Trust-MRC Institute of Metabolic Science, Cambridge, United Kingdom; 193)Department of Biostatistics, University of Liverpool, Liverpool, United Kingdom;
